# ACE: a versatile contrastive learning framework for single-cell mosaic integration

**DOI:** 10.1101/2024.11.28.625798

**Authors:** Xuhua Yan, Jinmiao Chen, Ruiqing Zheng, Min Li

## Abstract

The integration of single-cell multi-omics datasets is critical for deciphering cellular heterogeneities. Mosaic integration, the most general integration task, poses a greater challenge regarding disparity in modality abundance across datasets. Here, we present ACE, a mosaic integration framework that assembles two types of strategies to handle this problem: modality-alignment based strategy (ACE-align) and regression-based strategy (ACE-spec). ACE-align utilizes a novel contrastive learning objective for explicit modality alignment to uncover the shared latent representations behind modalities. ACE-spec combines the modality-alignment results and modality-specific representations to construct complete multi-omics representations for all datasets. Extensive experiments across various mosaic integration scenarios demonstrate the superiority of ACE’s two strategies over existing methods. Application of ACE-spec to bi-modal and tri-modal integration scenarios showcases that ACE-spec is able to enhance the representation of cellular heterogeneities for datasets with incomplete modalities. The source code of ACE can be accessed at https://github.com/CSUBioGroup/ACE-main.

## Introduction

Single-cell RNA sequencing (scRNA-seq) allows for quantifying RNA molecular traits at the single-cell level, enabling applications from cellular heterogeneity characterization [1] to regulatory network inference [2]. However, biological processes in cells involve multiple molecular types, such as DNA, RNA, and protein, resulting in an intrinsic need for multimodal understanding. Advances in single-cell sequencing have enabled technologies that can capture multiple molecule types. For example, cellular indexing of transcriptomes and epitopes (CITE-seq) captures RNA expression and cell surface protein abundance [3], 10x Genomics Multiome captures RNA expression alongside DNA fragments associated with regions of open chromatin [4] and DOGMA-seq [5] simultaneously measures chromatin accessibility, gene expressions, and protein abundances. To gain biological insights from such rich resources, data integration has emerged as a key challenge.

Ricard et al. [6] categorize data integration tasks on single-cell data into four scenarios: horizontal, vertical, diagonal and mosaic. Horizontal integration, also termed batch correction, refers to the scenario in which all data batches share the same modality. Vertical integration refers to the scenario where data batch is measured with multiple modalities. Diagonal integration refers to the scenario where data batches do not share any modality. Mosaic integration refers to the most general scenario where different batches are profiled with various modalities, like a grid that’s concatenated using various horizontal, vertical and diagonal scenarios. Among the four scenarios, mosaic integration is the most challenging one.

Similar to horizontal integration, mosaic integration also has the goal of preserving cellular heterogeneities and aligning data batches [7]. The distinction is that in mosaic integration, the inter-batch difference not only exist within the same modality but also come from difference in modality abundance. Different modalities may carry different information content and different batches can be measured with different modalities. Existing mosaic integration methods mainly adopt regression-based strategy to solve this problem. In specific, they attempt to recover the unmeasured modality for each batch using their measured modalities. For example, StabMap sets the batch measured with multiple modalities as reference (bridge batch) and calculates its low-dimensional embeddings which contain information from multiple modalities [8]. Then, on the bridge batch, it trains regression models to predict these embeddings based on single-modality profile and applies these regression models to other batches with single modality only. The core idea of Cobolt [9] is similar, which uses multi-modal variational auto-encoder (MVAE) [10] to embed all batches first and then trains XGBoost [11] to predict the reference embeddings from single modality. CLUE [12] also adopts the MVAE framework, but it doesn’t predict the reference embeddings directly. Instead, CLUE predicts the unmeasured modality based on the measured information and aggregates all modalities for each batch. Some of mosaic integration methods adopt non-negative matrix factorization to decompose each batch into cell factors and feature-specific factors, such as UINMF [13] and scMoMaT [7]. Those cell factors are expected to represent shared heterogeneities across all batches.

Here, we develop a novel mosaic integration framework, Align and CompletE (ACE), which builds on a new integration strategy and can be flexibly adjusted to encompass the output of regression-based methods. Specifically, we propose to build a modality-aligned latent space in which cells with similar biological signals are concentrated rather than batch labels or modality labels. In other words, we aim to extract the shared latent representations behind modalities. To achieve this target, we utilize contrastive learning to align modalities and propose a new learning objective to address the modality gap phenomena caused by the commonly used InfoNCE loss [14, 15]. Then, we can construct consensus latent representations across batches based on modality-aligned outputs. However, we also realize that modality alignment can erase the unique biological variations in each modality, especially when the disparity in the information content between modalities is significant. In this case, we think regression-based strategy are more suitable. Accordingly, our method can be flexibly adapted to encompass regression-based outcomes: we construct modality-specific embeddings by learning independent modality encoders and then recover the embeddings of missing modalities through a cross-modality matching approach based on prior alignment results. Together, we propose a mosaic integration framework which can provide modality alignment-based outputs and regression-based outputs, facilitating its use in various integration scenarios. For convenience, we refer the alignment-based outputs as ACE-align, and regression-based outputs as ACE-spec.

We evaluated the two types of outputs on four integration scenarios which cover different modality compositions (e.g., CITE-seq, Multiome-seq), different number of modalities (2 or 3 modalities), existence of horizontal batch effects, different cell type composition across batches. Experimental results show that both outputs of ACE’s framework can achieve superior mosaic integration performance. Moreover, we showcased that ACE-spec could enhance representations of cellular heterogeneities, help refine cell type annotations, and reveal differences between cell subpopulations. Finally, we validated the robustness of our framework across various aspects, including the loss function, sensitivity to hyperparameters, choice of batch correction methods and clustering methods.

## Method

### Overview of ACE

Let 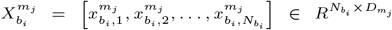 denotes inputs of one data batch *b*_*i*_ measured with modality *m*_*j*_ where 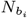 denotes the number of cells and 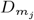 denotes the input dimensions of modality 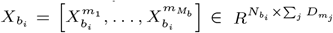 denotes one data batch *b*_*i*_ measured with 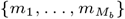 modalities and 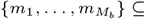 {RNA, ATAC, Histone mark, Protein}. The general target of mosaic integration task is to project each data batch *b*_*i*_ into a consensus latent space 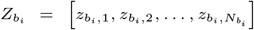, in which cells with similar biological signatures are clustered while cells with dissimilar biological signatures are separated.

ACE requires the existence of “bridge” batches in a mosaic dataset. More specifically, if each modality is seen as a node, then one batch that’s measured with two modalities, RNA and protein, connects nodes of RNA and protein. ACE requires all modalities in a dataset connected through several batches. These bridge batches lay the foundation for modality alignment. To remove within-modality batch effects across batches (if they exist), ACE applies harmony [16] to perform horizontal integration within each modality. Those low-dimensional representations are used as inputs of different modalities for each batch.

As mentioned in previous section, our framework generates two types of outputs: ACE-align and ACE-spec. ACE-align is derived from a modality alignment strategy which creates a shared latent space to align representations across modalities while preserving biological variations within each modality. Building on ACE-align, ACE-spec extends this framework by constructing modality-specific representations and imputing missing modalities through a cross-modality matching-based strategy. An overview of ACE is shown in Fig. 1.

**Fig. 1.**
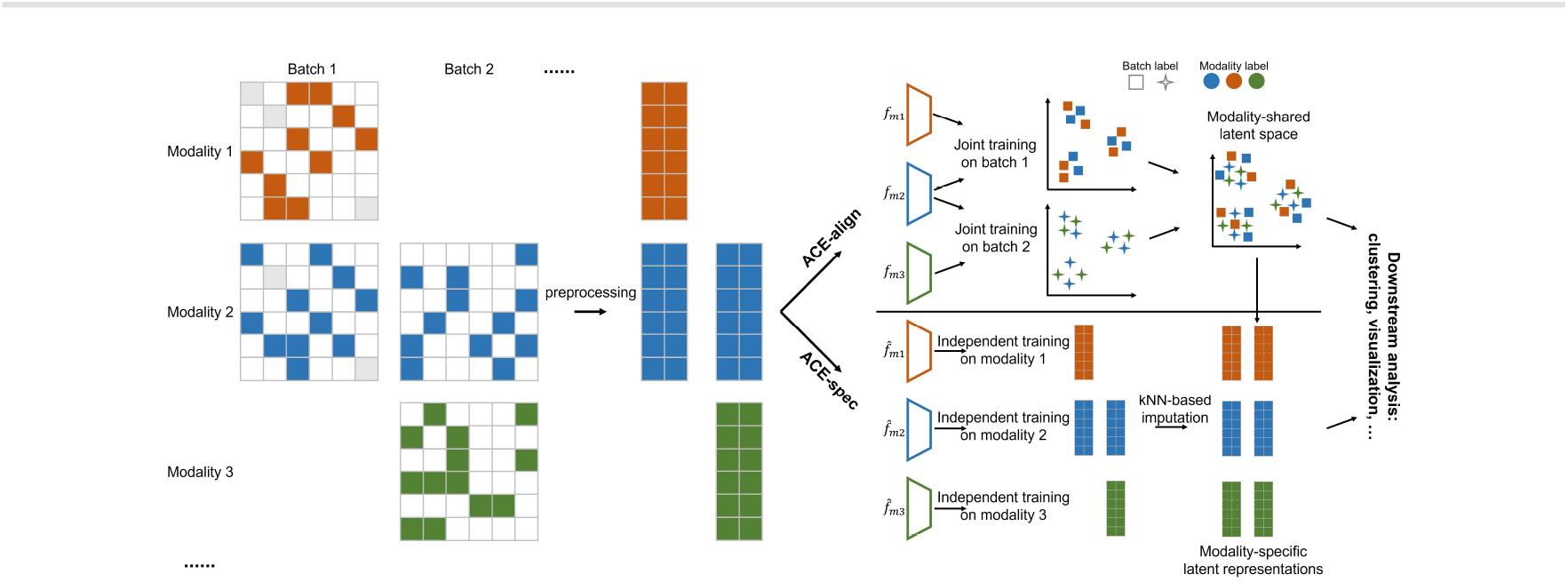
Architecture of ACE. ACE assembles two types of outputs: ACE-align and ACE-spec. Before model training, each modality profiles are preprocessed through dimension reduction and batch correction (harmony), respectively. ACE-align takes the low-dimensional representations as input and jointly trains the modality encoders on bridge batches to achieve modality-shared latent space. ACE-spec independently trains modality encoders on corresponding modality inputs, and then utilizes the modality-aligned latent space to impute the missing modality-specific representations. Finally, both outputs eliminate the inter-batch differences in horizontal batch effects and modality abundance.

### ACE-align

To build a modality-shared latent space, we utilize contrastive learning to perform modality alignment. Contrastive learning (CL) is a popular self-supervised learning framework which learns representations by concentrating predefined positive pairs while separating negative pairs [15]. CL loss has been validated as an effective objective for representation alignment because it can well align batches while preserving variations between cells [17], which motivates us to apply CL to perform modality alignment. Before us, NOVEL [18] and MatchClot [19] also used the common CL loss, InfoNCE [15], to learn aligned representations between two modalities of the same cells. However, NOVEL and MatchClot can only integrate datasets with two modalities and cannot be directly extended to datasets with three modalities. Additionally, the InfoNCE loss they adopted encounters the modality gap problem, which has been observed in the field of computer vision [14]. In other words, representations from different modalities are still clearly separable after alignment, indicating a poor quality of modality-shared latent space.

For ACE-align, embeddings of different modalities from the same cell form positive pairs. For instance, if a cell *i* is measured with three modalities *m*_1_, *m*_2_ and *m*_3_ (with embeddings 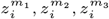), we define three positive pairs 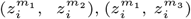 and 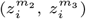. Negative pairs are constructed by randomly sampling embeddings from different cells, irrespective of modality (Supplementary Fig. S1a). For convenience, we first discuss two-modality situation. During training, one mini-batch of cells with size *n* is sampled from a bridge batch: 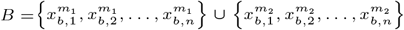 Each modality is projected by a separate encoder 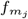 into a low-dimensional latent space 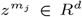. CL loss is used to train all encoders. InfoNCE is a commonly used CL loss function in cross-modal alignment research [20]. It’s defined as [15]:

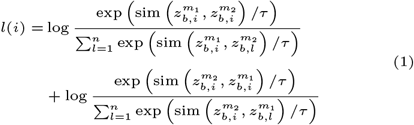

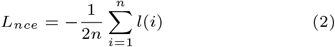

where sim 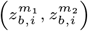 denotes the cosine similarity between two embeddings and *τ* can be a positive constant or a learnable parameter. By optimizing above objective, embeddings of different modalities from the same cell are pushed closer than embeddings of different modalities from different cells. However, one critical limitation of InfoNCE loss is that it ignores the relationships within intra-modality embeddings. Specifically, eq (1) only optimizes inter-modality embeddings of the same cell to be closer than inter-modality embeddings between different cells, indicating the intra-modality embeddings between different cells can be still closer than inter-modality embeddings of the same cell. As a result, the modality gap remains. Our solution is to add intra-modality embeddings between different cells as negative pairs in eq (1). For convenience, we define 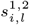 as a shorthand for sim 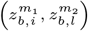 Then, our proposed CL loss is defined as:

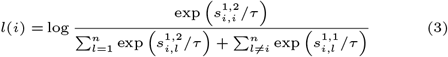

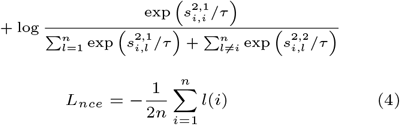

When existing *M* (*M >* 2) modalities to be aligned, we generalize above objective as:

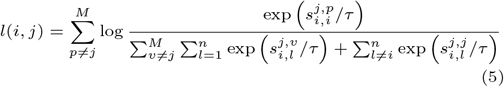

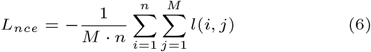

By optimizing this objective, all modalities’ embeddings of one cell are concentrated without modality gap, and embeddings from different cells are separated. After modality alignment, we can build the consensus representation across batches by combining embeddings from measured modalities. One common approach is weighted average:

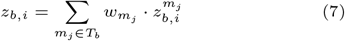

where *T*_*b*_ denotes the set of modalities that batch *b* is measured with and |*T*_*b*_| denotes the number of elements in 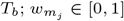 denotes the weight of modality *m*_*j*_ and 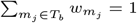.

### ACE-spec

We are aware of that modality alignment can bring loss of unique information inherent in each modality. Especially when the differences in biological variations are large between modalities, the modality with richer information will be biased to modality with poorer information [21]. To address this problem, we extend ACE-align to ACE-spec. Our approach is straightforward: learning modality-specific representations effectively preserves their original variations but fails to address inter-batch differences in modality abundance; thus, we design a strategy to impute the missing modalities for each batch based on the results of ACE-align.

Within each modality *m*_*j*_, we employ contrastive learning to train an encoder network, denoted as 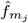 to project inputs into a modality-specific latent space. The training is conducted independently for each modality. Positive pairs are defined as each cell paired with itself, while negative pairs are constructed by randomly sampling embeddings from different cells (Supplementary Fig. S1b). For a mini batch of *n* cells sampled from modality *m*_j_, denoted as 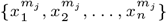, the loss function is defined as:

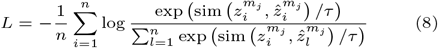

where 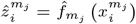 and sim 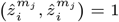, indicating that the objective is to maximize the separation between negative pairs. In other words, optimizing this function enables the model to learn discriminative features between different cells, which we believe helps to better preserve biological variations within each modality.

To impute embeddings for missing modalities, we combine the modality-shared embeddings *z* and modality-specific embeddings 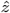 to infer the embeddings for missing modalities. In detail, let 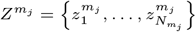 denotes aligned embeddings of modality *m*_*j*_ from all batches, where 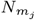 denotes the number of cells that are measured with modality 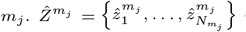 denotes the specific embeddings of modality *m*_*j*_ from all batches. Let 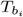 denote the set of modalities that batch *b*_*i*_ is measured with. 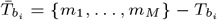 denotes the set of missing modalities of batch *b*_*i*_. For each modality 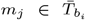, we first perform cross-modal matching between 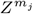 and 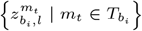. Specifically, for each 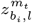, we find its *k* nearest neighbors *Q* = {*q*_1_, *q*_2_, …, *q*_*k*_} in 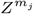. Then, embedding of modality *m*_*j*_ imputed from modality *m*_*t*_ is defined as:

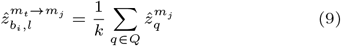

where *m*_*t*_ → *m*_*j*_ denotes the embedding is imputed based on modality *m*_*t*_. For each modality in 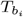, we repeat above imputation process and the final imputed embedding for missing modality *m*_*j*_ of batch *b*_*i*_ is:

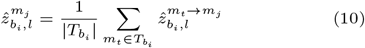

where 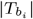 denotes the number of elements in 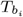. After the imputation, we average the representations from all modalities to get the consensus representation for each batch, which is similar to eq (7).

The workflow of ACE-spec is illustrated in Supplementary Fig. S2. Essentially, the computation process of ACE-spec is similar to regression-based methods, but our method completely decouples the modality-specific learning process and cross-modality learning process, ensuring better preservation of unique information in each modality. Notably, our proposed imputation strategy can also be applied to impute raw omics features. The modification is to change 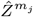 to the expression profiles of modality *m*_*j*_.

### Implementation detail

ACE-align employs modality-specific encoder networks to project inputs from each modality into a shared embedding space. Each encoder network consists of three fully connected layers. The RNA- and ATAC-specific encoders have output dimensions of 1024, 512, and 256 for the three layers, while the protein-specific encoder has output dimensions of 512, 2048, and 256 for the three layers. In each encoder, the first two fully connected layers are followed by an exponential linear unit (ELU) activation [22] and dropout regularization (*p* = 0.2) [23]. The contrastive learning loss is directly computed on the embeddings. The temperature parameter for contrastive learning is fixed at 0.1. ACE-align is trained using the Adam optimizer [24] with a learning rate of 2e-4, a batch size of 512, and 100 training epochs. ACE-spec shares the same network architecture and the temperature parameter with ACE-align. ACE-spec is trained using the Adam optimizer with a learning rate of 1.75e-4, a batch size of 512, and 10 training epochs. The models are implemented using PyTorch [25]. For embedding imputation, the number of nearest neighbors (*k*) is set to 2. When combining modality-specific embeddings to generate the final embeddings for ACE-spec, equal weights are assigned to all modalities. For ACE-align, the weights of RNA embeddings (or protein embeddings) are set to 1 for multi-modal batches. In single-modal batches, ACE-align assigns a weight of 1 to the respective modality.

### Datasets

We collected six datasets which cover different sequencing technologies, number of modalities and different scenarios of missing modalities. CITE and Multiome datasets are NeurIPS 2021 multi-modality competition [26] datasets sequenced from two types of technologies, CITE-seq and 10X genomics Multiome. CITE dataset contains two modalities, gene expression (RNA) and proteins (ADT) while Multiome dataset contains two modalities: gene expression (RNA) and chromatin accessibility (ATAC). Following data splitting scheme in the competition, we first divide all 12 batches in CITE datasets into two sets: training and testing. Training and testing sets contains 9 and 3 batches, respectively. Then, the RNA and ADT profiles of testing set are split into two parts: RNA-modal part and ADT-modal part, and they are treated as originating from different experiments. The task on CITE dataset is to integrate multi-modal (RNA+ADT) part (training set), RNA-modal part, and ADT-modal part. Similar to the splitting scheme on CITE, Multiome dataset is split into three parts: multi-modal part (10 batches), RNA-modal part (3 batches), and ATAC part (3 batches). BM-CITE dataset [27] is sequenced using CITE-seq, consisting of 30672 cells measured alongside a panel of 25 antibodies from bone marrow. BM-CITE consists of two batches. Following the split scheme of CITE and Multiome datasets, the two batches are split into three parts: multi-modal part (batch 1), RNA-modal part (from batch 2) and ADT-modal part (from batch 2). PBMC-Mult [28] is a dataset that is publicly available on the 10x website, where paired transcriptomes and ATAC-seq profiles are measured in 10412 peripheral blood mononuclear cells (PBMCs). PBMC-Mult consists of one batch, and we randomly split it into three parts: multi-modal part, RNA-modal part and ATAC-modal part. DOGMA dataset [5] contains two batches profiled by DOGMA-seq, which measures RNA, ATAC and ADT data simultaneously. Following the split scheme, these two batches are split into four parts: multi-modal part (from ‘control’ batch), RNA-modal batch (from ‘stim’ batch), ATAC-modal part (from ‘stim’ batch), and ADT-modal part (from ‘stim’ batch). CITE-ASAP dataset [5] contains four batches, in which two of them are profiled with CITE-seq and the other two are profiled with ASAP-seq. ASAP-seq measures RNA and ATAC. CITE-ASAP dataset is not split. Cell type labels of all datasets are from original studies which annotates each cell using all available modalities.

### Evaluation metrics

Following existing studies [7, 8], we evaluate mosaic integration methods from three aspects: biological preservation, batch correction, and modality alignment. In brief, biological preservation metrics compromise NMI and ARI. NMI and ARI both evaluate the overlap of two clustering. Score 0 corresponds to random clustering and score 1 corresponds to the perfect match for both NMI and ARI.

We used graph iLISI [29] as the batch correction metric. iLISI score is a diversity score to assess batch mixing degree. Following Stabmap [8], we not only use batch labels (iLISI_batch_) to compute iLISI but also used modality labels (iLISI_mod_) to compute iLISI. For example, if one batch is measured with RNA and ADT, then its modality label is RNA+ADT, whereas if one batch is measured with RNA, then its modality label is RNA. It can be regarded as evaluation at two levels of resolution, in which modality label is at coarse-grained resolution. Score 0 corresponds to separation of batches and score 1 corresponds to the perfect mixing of batches.

Modality alignment evaluation metrics compromise FOSCTTM (fraction of samples closer than the true match) [30] and matching score (MS) [26]. Both metrics evaluate the alignment between embeddings of different modalities for the same cells. Score 1 indicates perfect modality alignment for both metrics. The details of calculating each metric can be found in Supplementary Note A1.

Following scIB [29], all metrics are aggregated into three scores: bio-conservation score, batch-correction score, and modality-alignment score. For each method *q*_*i*_ on a dataset, the three scores are calculated via:

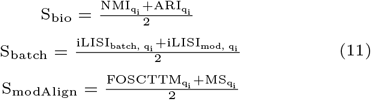

Following scIB, each metric is min-max scaled among compared methods before metrics aggregation so that all metrics have equal weights [29]. Finally, three scores are aggregated into the overall score as follows:

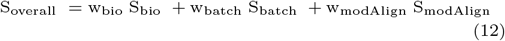

If the dataset doesn’t support evaluation of modality alignment metrics, we used w_bio_ = 0.6 and w_batch_ = 0.4, following the scIB benchmarking approach [29]. This weighting emphasizes the importance of preserving biological information while accounting for batch effects. When the dataset supports evaluation of modality alignment metrics, we used w_bio_ = 0.4, w_batch_ = 0.3, w_modAlign_ = 0.3, following the scMIB benchmarking proposed by MIDAS [31]. This setup similarly prioritizes biological conservation by giving it a higher weight.

## Results and discussion

### ACE shows superior performance in bi-modal mosaic integration

We organized four cases to evaluate mosaic integration methods across various batch effects and cell type composition scenarios. Case 1 includes BM-CITE and PBMC-Mult datasets, which lack intra-modality batch effects and have similar cell type compositions across batches. Case 2 includes CITE and Multiome datasets, which exhibit intra-modality batch effects but maintain similar cell type compositions. In case 3, we created multiple datasets by selecting different cell types from each part of the BM-CITE dataset and adjusted the proportion of shared cell types among batches (proportion= 0.1, 0.2, 0.4, 0.8) to simulate variations in cell type composition without batch effects. For case 4, we applied the same selection strategy to the CITE dataset to generate multiple datasets with both intra-modality batch effects and varying cell-type compositions. We compared ACE with five state-of-the-art bi-modal mosaic integration methods: Cobolt, CLUE, MatchClot, scMoMaT, and StabMap. Note that Cobolt, CLUE, scMoMaT, and StabMap cannot well handle intra-modality batch effects and thus we added harmony as their post-processing steps for comprehensive benchmarking, which led to four additional methods: Cobolt-harmony, CLUE-harmony, scMoMaT-harmony, and Stabmap-harmony. Detailed method settings can be found in Supplementary Note A2.

On both datasets of case 1, ACE-spec achieves the highest overall scores and followed by ACE-align (Fig. 2a and Supplementary Figs. S3a-b). ACE-spec has the highest bio-conservation scores and the highest modality alignment scores. Its batch correction scores are also comparable to the top performers. ACE-spec outperforms ACE-align by 6% with respect to ARI on average, indicating that using modal-specific representations can help preserve cellular heterogeneity. We visualized the latent embeddings of all methods on both datasets using UMAP [32] (Fig. 2b and Supplementary Fig. S4), and the visualization shows that in ACE-spec outputs, batches are well mixed and cell types are clearly separated. For example, CD4 Naïve and CD8 Naïve are well separated in ACE-spec’s outputs while for other methods, those two types are basically connected. Notably, MatchClot has the third highest modality alignment scores on BM-CITE dataset, but its embeddings show clear separation between batches that are measured with different modalities, which is the modality gap phenomenon and explains its 0 batch correction scores on this dataset. However, such phenomena are not observed in ACE-align’s embeddings, demonstrating that our proposed CL loss can better align embeddings between batches and are more suitable for mosaic integration tasks (refer to Supplementary Note B for further details). For the reason why MatchClot’s embeddings on PBMC-Mult dataset do not show the modality gap (Supplementary Fig. S4b), we think it’s probably because of greater information content disparity between protein and RNA, compared to that between ATAC and RNA.

**Fig. 2.**
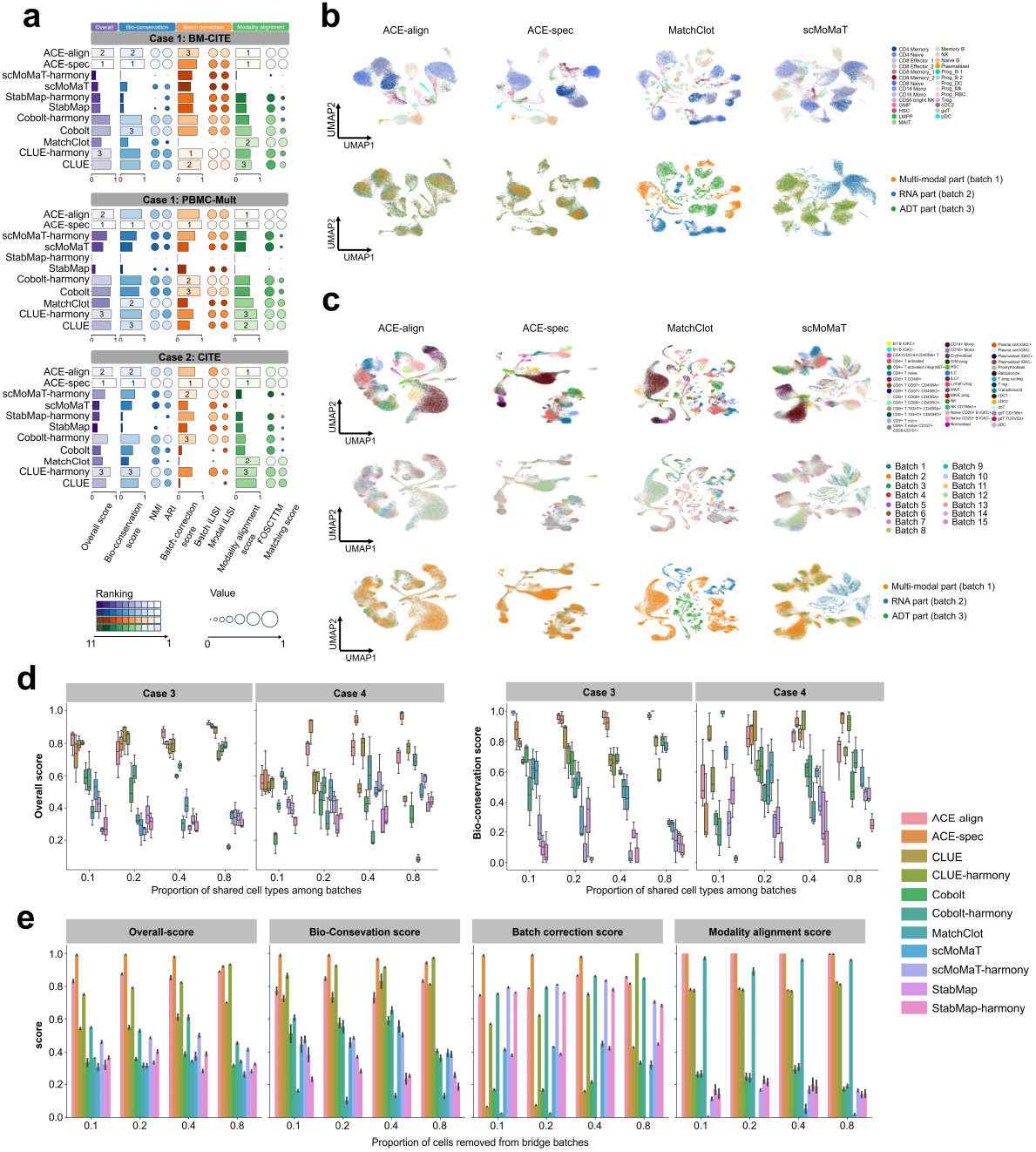
Bi-modal mosaic integration benchmark. (a) Overall scores for all compared methods in case 1 and case 2. We labeled the top three methods for each score. (b) UMAP plots for embeddings of ACE-align, ACE-spec, MatchClot and scMoMaT on BM-CITE dataset. Cells in the first row are colored by cell types and colored by modal labels (and batch labels) in the second row. (c) UMAP plots for embeddings of ACE-align, ACE-spec, MatchClot and scMoMaT on CITE dataset. Cells in the first row are colored by cell types, colored by batch labels in the second row, and colored by modal labels in the third row. (d) Overall scores and bio-conservation scores of all methods in case 3 and case 4. (e) Benchmarking of all methods’ robustness against the number of cells in bridge batches.

In case 2 where each dataset contains larger number of cells and more cell types than case 1, ACE-spec still achieves the highest overall scores, and followed by ACE-align (Fig. 2a, Supplementary Fig. S3c-d and S7). The bio-conservation scores of ACE-spec are also the highest and its modality alignment scores rank first and second on two datasets, respectively. Its batch-correction performance is the best on CITE dataset and competitive on Multiome dataset. Despite ACE-align not achieving the highest scores on these datasets, its performance remains comparable to the leading method. UMAP visualizations also show that ACE-spec produces well mixed batches and better separation of cell types (Fig. 2c and Supplementary Fig. S8). Similar to case 1, we observed modality gap phenomena in MatchClot’s results for CITE dataset while it does not appear in ACE-align’ results.

In case 3 and case 4, we did not evaluate the modality alignment score because the number of matched test cell pairs can be scarce. With different proportion of shared types, ACE-align and ACE-spec consistently attain the top overall scores (Fig. 2d and Supplementary Figs. S9a-b). When the proportion above certain thresholds, ACE-spec shows the superior performance over other methods. When the proportions are small, ACE-align and ACE-spec both rank within the top five among all methods, indicating their good generalization capabilities. Notably, when the proportions are small (≤ 0.1), ACE-align shows higher bio-conservation scores (and overall scores) than ACE-spec on both cases. The reason is that when the number of shared cell types among batches is small, it’s very likely to match cells with different cell types, eroding the original signal of cellular heterogeneity.

Finally, we evaluated the robustness of all methods against the number of cells in bridge batches. Specifically, we randomly removed certain proportion of cells from the bridge batches in CITE dataset (proportion=0.1, 0.2, 0.4, 0.8, remaining 60000, 53000, 40000, 13000 cells respectively) and reevaluated all the methods on these new datasets. Generally, with the proportion of removed cells increasing, all methods’ performance decreased with respect to specific metrics (Supplementary Fig. S9c). However, ACE-align and ACE-spec still attain the top three overall scores among all methods (Fig. 2e).

### ACE achieves competitive performance in tri-modal mosaic integration

We further applied ACE to tri-modal integration. DOGMA dataset and CITE-ASAP dataset were used in this experiment. They both exhibit intra-modality batch effect and following bi-modal settings, we organized tri-modal case 2 and tri-modal case 4 on them to evaluate integration methods. Specifically, case 2 includes DOGMA and CITE-ASAP datasets. In case 4, following bi-modal settings, we randomly sampled different cell types from each part in DOGMA dataset respectively and constructed multiple datasets with various proportions of shared types among batches. We used Cobolt, CLUE, scMoMaT, and StabMap for comparisons. MatchClot was not included because it cannot handle tri-modal integration situations. We also added harmony as the post-processing step for CLUE, Cobolt, scMoMaT, and Stabmap. Note that CITE-ASAP dataset does not contain test cell pairs, we didn’t evaluate modality alignment on it.

In tri-modal case 2, ACE-spec achieves the highest overall score on DOGMA dataset and followed by ACE-align (Fig. 3a and Supplementary Fig. S10a). Their bio-conservation scores rank in the top two and their batch-correction scores rank in the top three among all methods. UMAP plots also show that within ACE’s outputs, batches are well mixed and cell types are clearly separated (Fig. 3b and Supplementary Fig. S11a). Surprisingly, on CITE-ASAP dataset, StabMap reaches much higher bio-conservation scores than other methods, helping it achieve the highest overall score (Fig. 3c). UMAP plots show that StabMap indeed clearly separates four cell types in CITE-ASAP dataset whereas ACE-align and ACE-spec separate T cells and Myeloid cells into several subpopulations (Fig. 3d and Supplementary Fig. S11b). The superior performance of StabMap on the CITE-ASAP dataset can be attributed to the low resolution of annotations, which simplifies the task of preserving cellular heterogeneities.

**Fig. 3.**
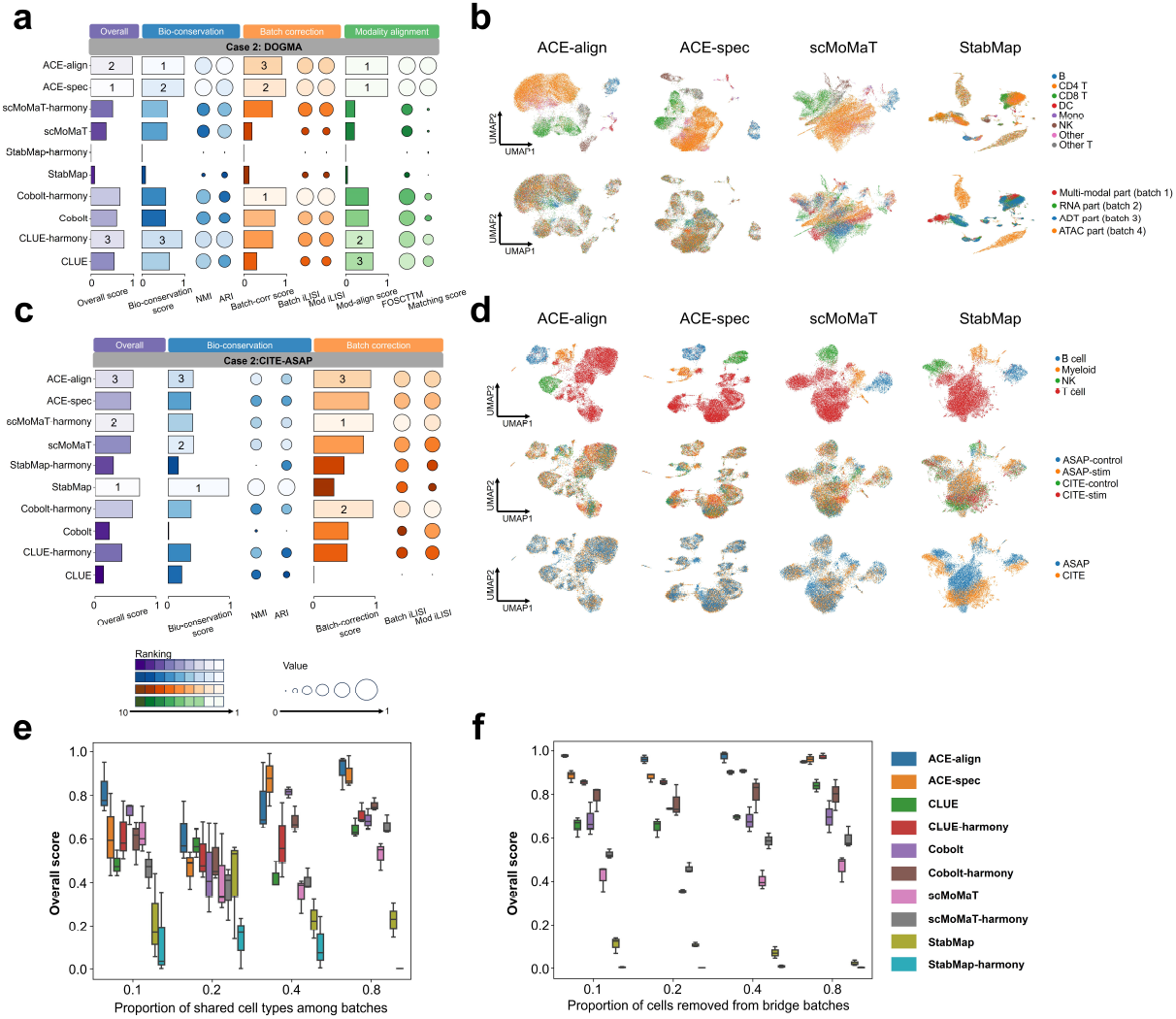
Tri-modal mosaic integration benchmark. (a) Overall scores of all methods on the DOGMA dataset in case 2. We labeled the top three methods for each score. (b) UMAP plots for embeddings of ACE-align, ACE-spec, scMoMaT and StabMap on DOGMA dataset. Cells in the first row are colored by cell types and colored by modal labels (batch labels) in the second row. (c) Overall scores of all methods on the CITE-ASAP dataset in case 2. (d) UMAP plots for embeddings of ACE-align, ACE-spec, scMoMaT and StabMap on CITE-ASAP dataset. Cells in the first row are colored by cell types, colored by batch labels in the second row, and colored by modal labels in the third row. (e) Overall scores of all methods in tri-modal case 4. (f) Benchmarking of all methods’ robustness against the number of cells in bridge batches.

In tri-modal case 4, ACE-align consistently ranks within the top two in terms of overall scores across various proportions of shared types. (Fig. 3e and Supplementary Fig. S12a-b). With the proportion increasing, ACE-spec’s overall scores gradually improve and reach similar or even exceed ACE-align, which is similar to the case 4 in bi-modal integration. The main reason is that low proportions of shared types among batches can easily result in incorrect matching of cells. We also evaluated the robustness of all methods against the number of cells in bridge batches. Specifically, we randomly removed certain proportion of cells from the bridge batches in DOGMA dataset (proportion=0.1, 0.2, 0.4, 0.8, remaining 6900, 6100, 4600, 1500 cells respectively) and reevaluated all the methods on these new datasets. Despite all methods’ performance decreasing with the removed cells increasing (Supplementary Fig. S12d), ACE-align and ACE-spec consistently achieves the top overall scores (Fig. 3f and Supplementary Fig. S12c).

### ACE-spec helps refine cell-type annotations of CITE-ASAP dataset

Note that in the integration task of CITE-ASAP dataset, Stabmap achieves top bio-conservation scores whereas on the rest of datasets, its scores are at a low level. This is because the annotation resolution of CITE-ASAP dataset is so low that preservation of biological variations is easily achieved. However, the embeddings of ACE-spec show stronger cellular heterogeneity than the original annotations, which inspires us to use ACE to refine the cell type annotations. We applied Louvain clustering algorithm on the embeddings of four batches using Scanpy [33] (resolution=1.0) and obtained 17 clusters (Fig. 4a). For a comparison, we also performed clustering on other methods’ embeddings with the same resolution.

**Fig. 4.**
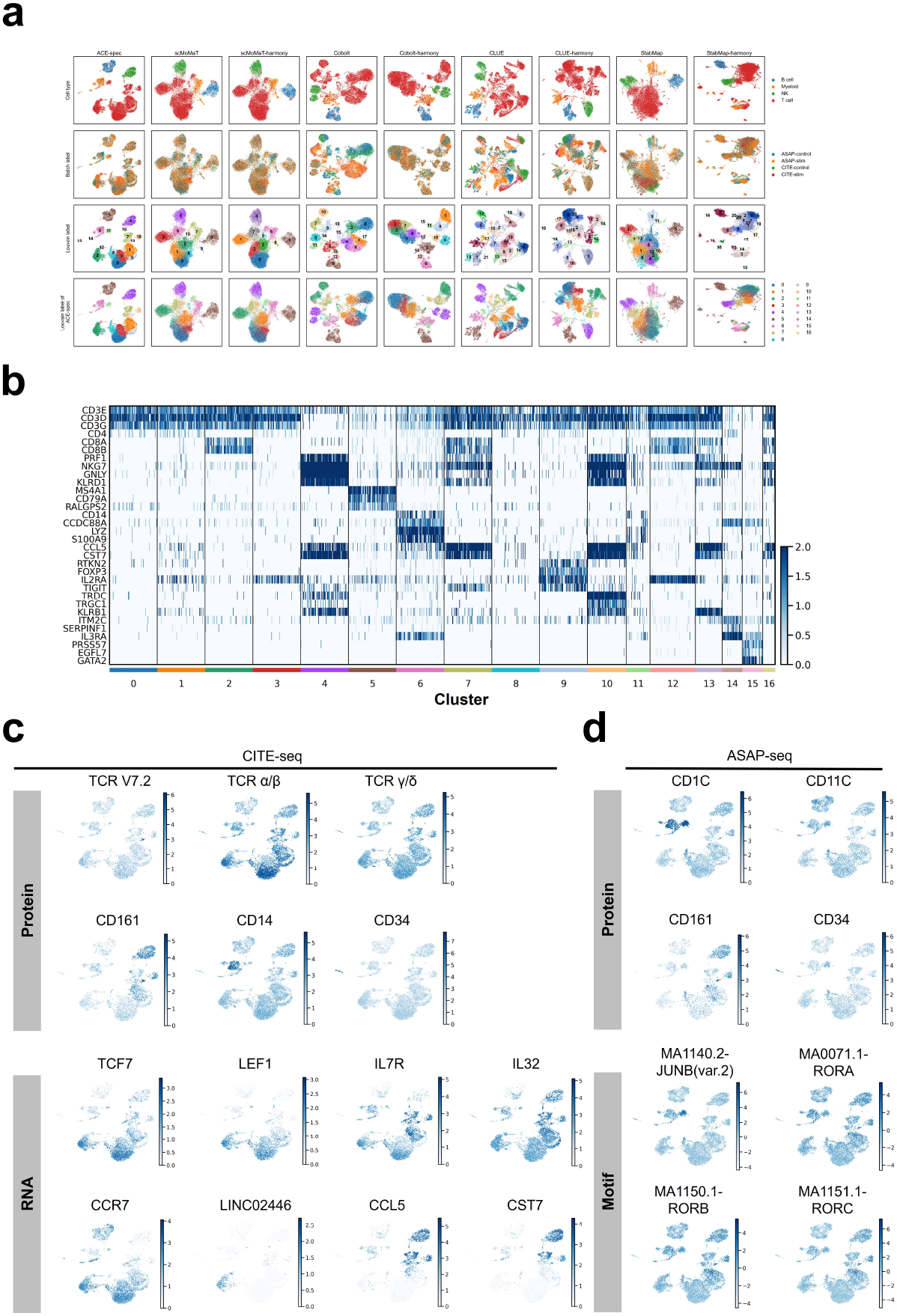
Analysis of ACE-spec’s results on the CITE-ASAP dataset. (a) UMAP plots of embeddings from different integration methods on this dataset. Cells are colored by original annotation (the first row), batch label (the second row), Louvain clustering label (the third row) and clustering labels from ACE-spec (the last row). (b) Expression heatmap of marker genes of known cell types in all clusters. (c) Expression heatmap of known marker genes and marker proteins in the batches measured with CITE-seq. (d) Expression heatmap of known marker genes and marker motifs in the batches measured with ASAP-seq.

As shown in Fig. 4a. All methods’ embeddings successfully separate the four annotated cell types. StabMap-harmony produces the largest number of clusters, but many appear to be outliers because there are significant discrepancies between the outputs of StabMap and StabMap-harmony. When comparing the clustering results of ACE-spec, scMoMaT, scMoMaT-harmony and Cobolt-harmony, we can observe that ACE-spec achieves the most compact clusters and clearest separation boundaries between clusters. The mapping of ACE-spec’s clustering labels onto the UMAP of other methods is largely consistent with their own clustering results, but ACE-spec provides a finer granularity in clustering resolution. We also applied weighted nearest neighbors (WNN) analysis [34] on the batches measured with CITE-seq and batches measured with ASAP-seq respectively. When projecting ACE’s cluster labels to the WNN UMAP plots, we observed the majority of clusters labels are consistent with the grouping in WNN analysis results (Supplementary Figs. S13a-b). The differences are as follows: in the batches measured with CITE-seq, WNN further separates clusters 1 and 2 into two groups, respectively, but mixes clusters 7 and 13, as well as clusters 2 and 12. In the batches measured with ASAP-seq, WNN mixes clusters 1 and 9, as well as clusters 6 and 14.

Next, we tried to annotate these clusters. Taking the two batches measured with CITE-seq as an example, cluster 4 corresponds to original Natural Killer (NK) cell and cluster 5 corresponds to original B cell. The upregulated genes *PRF1, NKG7, GNLY, KLRD1* [34] (*p <* 10^−10^ based on Wilcoxon test) and upregulated *MS4A1, CD79A, RALGPS2* [34] (*p <* 10^−10^ based on Wilcoxon test) validate the original annotations (Fig. 4b). The annotation refinement mainly happens within T cell and Myeloid cell populations. Clusters 0,1,2,3,7,8,9,10,12,13,16 have upregulated expression of T cell markers [34] (*CD3E, CD3D, CD3G, p <* 0.05), making them T cells (Fig. 4b). Clusters 0, 1, 3, 8, 9 have high expression of *CD4*, indicating they are CD4+ T cells (Fig. 4b). Clusters 2, 7, 12, 16 with high expression of *CD8A, CD8B* are CD8+ T cells (Fig. 4b). Clusters 10 and 13 belong to other T cells. Notably, cluster 11 consists of T cells in the original annotations, but any of *CD3D, CD3E, CD3G* is not highly expressed within them (*p >* 0.05). Instead, cluster 11 has upregulated expression of *CD11c* (*p <* 10^−10^) and *CD1c* [34, 35] *(p <* 0.05) and thus we annotated it as conventional dendritic cells (cDC).

Within CD4+ T cells, clusters 0, 3, 8 have high expression of *TCF7* and *LEF1* [34], making them naïve CD4+ T cells (Fig. 4c); cluster 1 has high expression of *IL7R* and *IL32* [34], making them CD4+ memory T cells (Fig. 4c); cluster 9 has high expression of *RTKN2, FOXP3, IL2RA, TIGIT* [34], making them T regulatory cells (Tregs) cells (Fig. 4b). Within CD8+ T cells, clusters 2 and 12 with high expression of *CCR7, LINC02446*, and *LEF1* [34] are naïve CD8+ T cells (Fig. 4c); clusters 7 and 16 with high expression of *IL7R, CCL5, CST7* are CD8+ memory T cells [34] (Fig. 4c). Within other T cells, cluster 13 has high expression of *KLRB1* [34] and surface protein TCR V*α*7.2, *CD161* [34, 35] *(Fig. 4b-c), making them mucosal-associated invariant T (MAIT) cells. Cluster 10 shows high expression of TRDC, TRGC1* [34] and surface protein TCR-*γδ* [36] (Figs. 4b-c), making them gamma delta T (gdT) cells.

Within the original myeloid cells, cluster 6 has high expression of *CD14, S100A9*, and *LYZ* [34] (Figs. 4b-c), making it CD14 monocytes. Cluster 14 is identified as plasmacytoid dendritic cell (pDC) due to its high expression of *ITM2C, SERPINF1, IL3RA* [34] (Fig. 4b). Cluster 15 is identified as hematopoietic stem and progenitor cell (HSPC) due its high expression of *PRSS57, EGFL7, GATA2* [34], and surface protein *CD34* [34] (Figs. 4b-c).

For each cluster, we performed KEGG pathway enrichment analysis on the top 200 differentially expressed genes (based on RNA data from CITE-seq) using clusterProfiler [37]. The results demonstrate a consistency between the enriched pathways and our cell type annotations (Supplementary Fig. S14). For instance, Natural Killer (NK) cell-mediated cytotoxicity is the most significantly enriched pathway in cluster 4, supporting our annotation of NK cells. In cluster 5, pathways such as B cell receptor signaling and Intestinal immune network for IgA production are enriched, further validating the cluster’s identification as B cells. Those clusters annotated as T cells display enrichment in pathways relevant to T cell function, including T cell receptor signaling, primary immunodeficiency, and cell adhesion molecules. Additionally, cluster 6, annotated as CD14+ monocytes, shows enrichment in pathways like phagosome and lysosome, which are critical for the phagocytic activity of monocytes [38]. For cluster 15, annotated as hematopoietic stem and progenitor cells (HSPCs), we observed enrichment in the hematopoietic cell lineage pathway.

Moreover, analysis on the other two batches measured with ASAP-seq can also validate our annotations. For example, surface protein *CD1c* and *CD11c* are highly expressed within cluster 11 (*p <* 0.01, Fig. 4d), consistent with above results. Cluster 13 has high expression of TCR V*α*7.2, *CD161* and cluster 15 has high expression of *CD34* (Fig. 4d and Supplementary Fig. S13c), confirming the annotations of MAIT cells and HSPC. We further used chromVar [39] to infer accessibility scores for known motifs. We found that peaks in cluster 11 were highly enriched for motifs for the transcription factor *JUNB* [40] (*p <* 10^−10^, Fig. 4d) that is essential for cDC identity [40], confirming our annotation for cluster 11. Peaks in our annotated MAIT cells are highly enriched for motifs for the pro-inflammatory transcription factor RORgammat (*p <* 10^−10^, Fig. 4d), consistent with existing studies [34]. Other motifs’ activity scores also support our annotations (Supplementary Fig. S13d).

### ACE-spec helps discriminate cellular heterogeneity in COVID-19 datasets

To further demonstrate ACE-spec’s ability in enhancing the representation of cellular heterogeneity, we consider a more complex task of mosaic integration for atlas-scale datasets from Coronavirus disease 2019 (COVID-19) patients. We collected a public scRNA-seq dataset [41] (referred as COVID19-RNA dataset) which analyzed the transcriptome of peripheral blood mononuclear cells (PBMCs) from 17 COVID-19 patients with moderate disease (n = 5), acute respiratory distress syndrome (ARDS) (severe, n = 6), or recovering from ARDS (recovering, n = 6). This dataset contains 69983 cells and 11 manually annotated cell types. A single cell resolution mass cytometry (CYTOF) dataset [42] spanning 160 patients and a total of 7.11 million cells was collected, which was generated from granulocyte depleted whole blood of COVID-19 patients, sepsis patients and healthy volunteers. We used the CITE-seq dataset (referred as CITE2 dataset) of 161764 PBMCs from healthy donors [34] as the bridge dataset. Following [43], we removed cells from individuals with sepsis in the CYTOF dataset, resulting in 116 samples and 5.17 million cells. The CYTOF dataset was then down sampled to 1000 cells per sample and finally it contains 116000 cells in total. The CITE2 dataset shares 31 surface protein features with the CYTOF dataset and shares 19668 gene features with COVID19-RNA dataset. UMAP visualizations for each dataset and their embeddings from ACE-spec are shown in Fig. 5a and Supplementary Fig. S15.

**Fig. 5.**
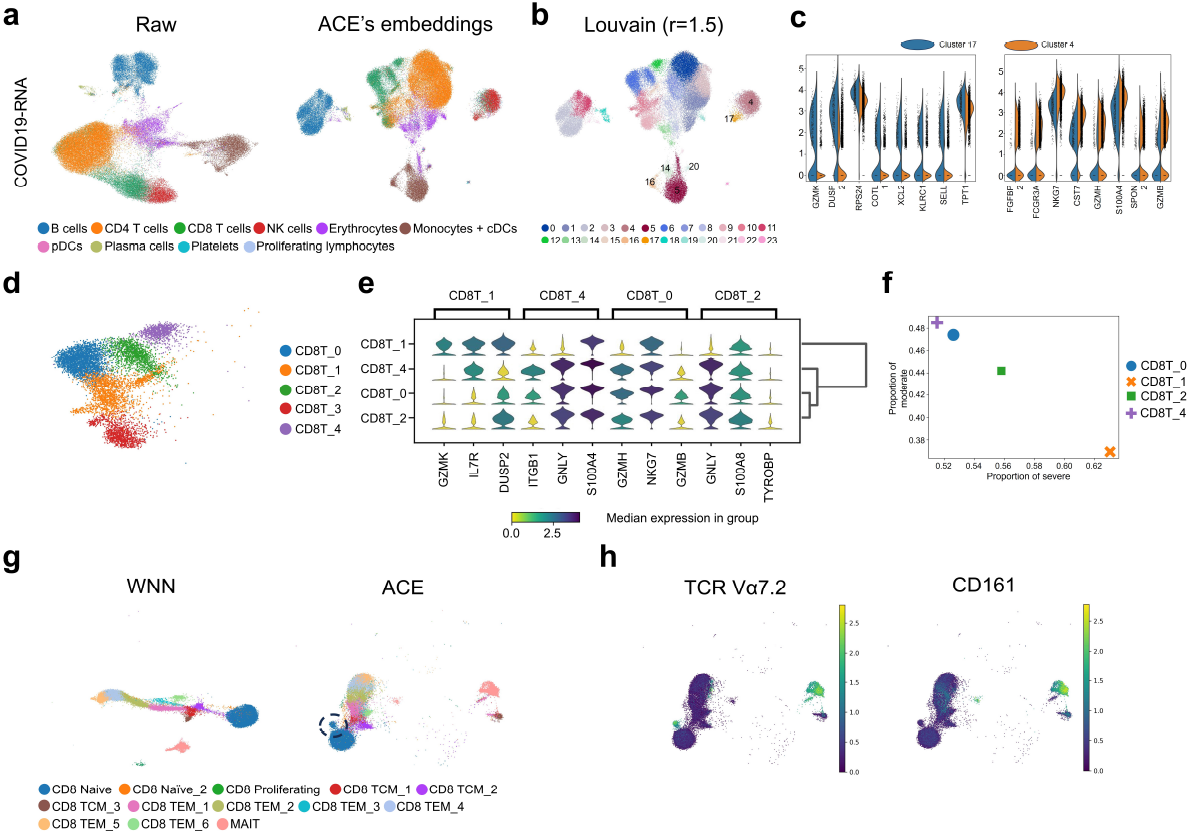
Analysis of ACE-spec’s results on the COVID-19 related datasets. (a) UMAP plots for raw expression profiles of COVID19-RNA dataset and its embeddings from ACE-spec. Cells are colored by original annotations. (b) UMAP plot of ACE’s embeddings for COVID19-RNA dataset. Cells are colored by clustering labels through performing Louvain on the embeddings. (c) The most significant DEGs between cluster 17 and 4. (d) Clustering labels within original CD8 T populations by performing Louvain on the embeddings of CD8 T populations. (e) Stacked violin plot displaying the top DEGs among four clusters. (f) Scatter plot showing the proportion of cells from different severity groups within four clusters. (g) UMAP plots of CD8 T populations and MAIT cells in the bridge dataset using embeddings from WNN analysis and ACE. The black dotted circle highlights a separated cluster from the CD8 Naïve cluster. Cells are colored by the level-3 annotations in the bridge dataset. (h) Heatmap of expression of TCR V*α*7.2 and CD161 for cells in (g).

ACE-spec enhanced the representation of cellular heterogeneity for COVID19-RNA dataset (Fig. 5b, clustering is performed on the embeddings using scanpy with 1.5 resolution). For example, within originally annotated NK cells, ACE-spec separates them into two clusters, 4 and 17. DEG analysis between these two clusters shows that *GZMK* is the most significantly upregulated gene in cluster 17 (*p <* 10^−10^) and *GZMB* is one of most significantly upregulated genes in cluster 4 (*p <* 10^−10^) (Fig. 5c). The expression difference between two clusters may be correlated with their proportion of cells from different patients. Specifically, within cluster 17, cells from moderate patients occupy 37% and cells from severe (severe and recovering) patients occupy 63% while within cluster 4, they occupy 44% and 56% respectively. Increasing number of cells from moderate patients within cluster 4 may lead to higher expression of *GZMB* and lower expression of *GZMK*, which is consistent with the finding that *GZMB* has increased expression in NK cells of mild patients and *GZMK* has predominantly increased expression in NK cells of severe patients [44].

Within the originally annotated Monocytes/cDC, ACE-spec separates them into clusters 5, 14, 16, 20. It’s noteworthy that ACE-spec groups a part of original erythrocytes into cluster 5. We observed that this small part of cells has high expression of *S100A8, S100A9, LYZ* and *CD14* [34], indicating it’s CD14 monocytes (Supplementary Fig. S16). Cluster 14 also highly expresses these genes, making it CD14 monocytes. Cluster 16 highly expresses *CDKN1C, FCGR3A, MS4A7*, and *HES4* [34], making it CD16 monocytes (Supplementary Fig. S16). Cluster 20 has high expression of *FCER1A, HLA-DQA1, CLEC10A*, and *CD1C* [34], making it cDC (Supplementary Fig. S16).

Within the original CD8 T cells, we observed that ACE-spec separates them into several groups, and we performed Louvain algorithm among them using scanpy (resolution=0.3), resulting in 5 groups (Fig. 5d). Cluster CD8T 3 highly expresses *CCR7, LINC02446, LEF1* and *OXNAD1* [34], making it Naïve CD8 T cells (Supplementary Fig. S16). Clusters CD8T 0, CD8T 1, CD8T 2, CD8T 4 highly express *CD8A, CCL5, GZMH*, and *KLRD1* [34], making them CD8 T Effector Memory (CD8 TEM) cells (Supplementary Fig. S16). Within these four clusters, CD8T 1 has notable expression differences to the other three (Fig. 5e), which may be correlated with its much higher proportion of cells from severe patients (Fig. 5f).

Moreover, for the bridge dataset, it also obtained enhanced representations. For example, within its level-3 annotations, ACE-spec isolates a group of cells (n=742) from CD8 Naïve cluster (Fig. 5g). We found this group of cells has high of expression of TCR V*α*7.2 but low expression *CD161*, marking them TCR V*α*7.2 + CD161-T cells (a subtype differing from MAIT cells and TCR V*α*7.2− conventional T cells, Fig. 5h) [45]. Together, ACE-spec enhances the representation of cellular heterogeneity, not only helping to define fine-grained cell type annotations but also revealing changes in phenotype.

### Parameter sensitivity and scalability

To test how ACE’s performance is affected by different parameter settings, we evaluated ACE on the CITE dataset as in bi-modal case 2 and on multiple downsampled CITE datasets as in bi-modal case 4. The hyper-parameters in ACE mainly include the temperature parameter *τ*, the latent dimension *d* and the number of nearest neighbors *k* for embedding imputation in ACE-spec. The learning rate and training epochs are fixed across experiments. Supplementary Figs. S17a-b show that ACE-align is generally insensitive to the choice of latent dimension, and it performs better when *τ* is not greater than 0.1. When *τ* is greater than 0.1, modality alignment scores of ACE-align will decrease. Also, ACE-spec gets higher modality alignment scores when *τ* is no greater than 0.1. The latent dimension has a notable impact on ACE-spec’s bio-conservation scores and batch correction scores. Specifically, (batch) iLISI score gets higher when *d* decreasing whereas bio-conservation scores get higher when *d* increasing. However, the influence of *d* on the batch correction scores is comparatively minor compared to its impact on the bio-conservation scores. So, we recommend using higher *d* (e.g., 256) for ACE-spec. As for the reason why bio-conservation scores favor large latent dimension, we believe it’s because better discrimination of cellular heterogeneity requires more feature dimensions. Varying k has little impact on the NMI and ARI scores of ACE-spec, while smaller values of *k* led to better iLISI scores of ACE-spec (Supplementary Fig. S17c). This is likely because the imputed embeddings that are averaged over more neighbors’ embeddings will exhibit reduced fidelity to the real ones, introducing batch effects within intra-modality embeddings. We recommend setting *k* to 2, as this value consistently attains superior batch correction and robust bio-conservation performance for ACE-spec.

We compared the scalability of ACE (align + spec) against other methods on the CITE2 dataset which simultaneously measures RNA and protein expressions in 161764 PBMCs from healthy donors [34]. We randomly selected cells from them with various proportions (1%, 10%, 20%, 40%, 80%, 100%) and then randomly partitioned them into three groups: multi-modal, RNA-modal, and protein-modal part. We evaluated the running time of all methods on a Linux server equipped with Intel(R) Core(TM) i9-10980XE CPU, 128 GB memory and a GeForce RTX 3090 GPU. Supplementary Fig. S17d demonstrates that ACE is the most time-efficient method and completes its run on 160000 cells within 3 minutes, suggesting its scalability for larger-scale datasets.

## Conclusion

In this study, we presented ACE, a mosaic integration framework for single-cell multi-omics data analysis. ACE assembled two strategies, ACE-align and ACE-spec, to handle the disparity in modality abundance across datasets. ACE-align applied contrastive learning for explicit modality alignment to construct shared latent space across modalities, thereby bridging the gap between modalities. We proposed a novel contrastive learning loss, which addressed the modality gap problem and achieved better modality alignment. ACE-spec utilized the results of ACE-align to impute the missing modality-specific representations, which helped preserve and better represent cellular heterogeneities.

We evaluated ACE-align and ACE-spec under various data integration scenarios using comprehensive metrics, and experimental results showed that both strategies achieved superior performance compared to other state-of-the-art methods. Each strategy has its own suitable applications. Generally, ACE-align fits scenarios with a small proportion of shared cell types across batches, whereas ACE-spec is more appropriate for scenarios with a large proportion of shared cell types across batches. Particularly, ACE-spec demonstrated outstanding performance in capturing cell-to-cell variations, rendering it an advantageous integration method for enhancing cellular representations of existing datasets.

We conducted extensive ablation studies to investigate the robustness of our proposed framework. First, we validated that the observed cellular heterogeneity was robust regardless of the choice of UMAP parameters (Supplementary Note C). Second, we validated that our proposed loss function significantly outperformed the InfoNCE loss in terms of batch correction and bio-conservation performance (Supplementary Note D). Third, we found that ACE was generally robust to the choice of batch correction methods applied prior to model training (Supplementary Note E). Fourth, we observed that ACE achieved higher NMI and ARI scores when using shared nearest neighbor (SNN)-based clustering algorithms [46, 47] compared to KMeans, while ACE-spec showed lower unsupervised metric scores under the same conditions (Supplementary Note F).

In most scenarios, the goal of data integration is to extract biological insights from the datasets, which is why the bio-conservation score is given a larger weight [29, 31]. However, in some cases, the objective may shift towards aligning batches or modalities. For these scenarios, a larger weight should be assigned to batch correction or modality alignment scores to better reflect the integration target. Based on our experiment results, ACE-align and ACE-spec demonstrated superior performance in biological conservation and modality alignment. Therefore, assigning larger weights to bio-conservation or modality alignment scores maximizes the overall performance of ACE’s framework, whereas placing a higher weight on batch correction reduces ACE’s comprehensive performance.

ACE can be used not only for data representation, but also for reconstruction of raw omics features of missing modalities. We performed detailed comparison between our method and other reconstruction methods (Supplementary Note G), including scVAEIT [48], TotalVI [49], and MultiVI [50]. Experimental results showed that our simple reconstruction strategy even achieved comparable performance to the state-of-the-art method, scVAEIT, and outperformed TotalVI and MultiVI with respect to overall scores and robustness. Moreover, UMAP plots confirmed that our reconstructed features showed better separation of cell types compared to scVAEIT.

Overall, ACE is a valuable tool for understanding of single-cell multi-modal data. We envisage that ACE will serve as a promising tool for the community of single-cell multi-omics data analysis.

## Supporting information

Supplementary information for the manuscript

## Code and data availability

All data, code, and materials used in the analyses is available at https://zenodo.org/records/10851161.

## CRediT author statement

Xuhua Yan: Methodology, Software, Analysis, Writing. Jinmiao Chen: Formal analysis, Writing. Ruiqing Zheng: Methodology, Data analysis, Writing. Min Li: Conceptualization, Supervision, Validation, Writing. All authors have read and approved the final manuscript.

## Acknowledgments

This work was supported in part by the National Natural Science Foundation of China under Grant (No. 62225209), and the Hunan Provincial Science and Technology Program (2019CB1007 and 2021RC4008).

## Supplementary material

Supplementary Data are available at Online.

