## Supplementary information for the manuscript for "ACE: a versatile contrastive learning framework for single-cell mosaic integration"

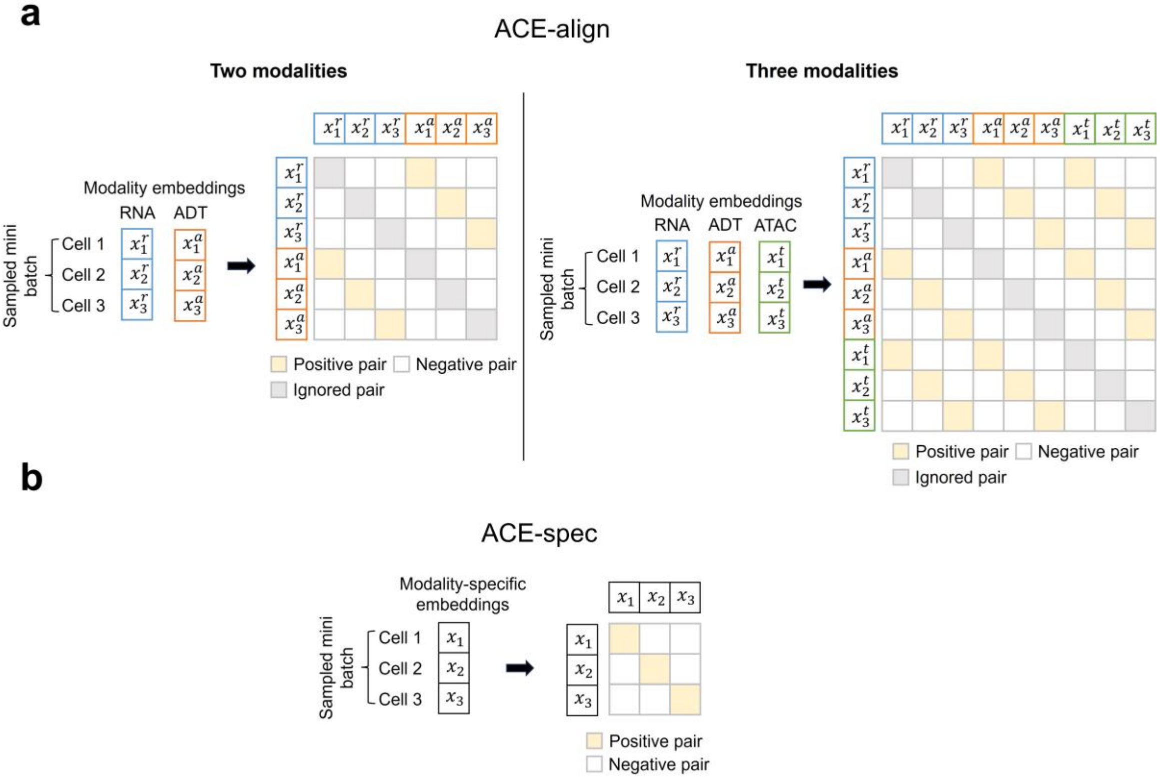


Supplementary Fig. S1. Examples of constructing positive and negative pairs for contrastive learning. (a) Construction examples of ACE-align in two-modality and three-modality cases, respectively. (b) Construction example of ACE-spec.

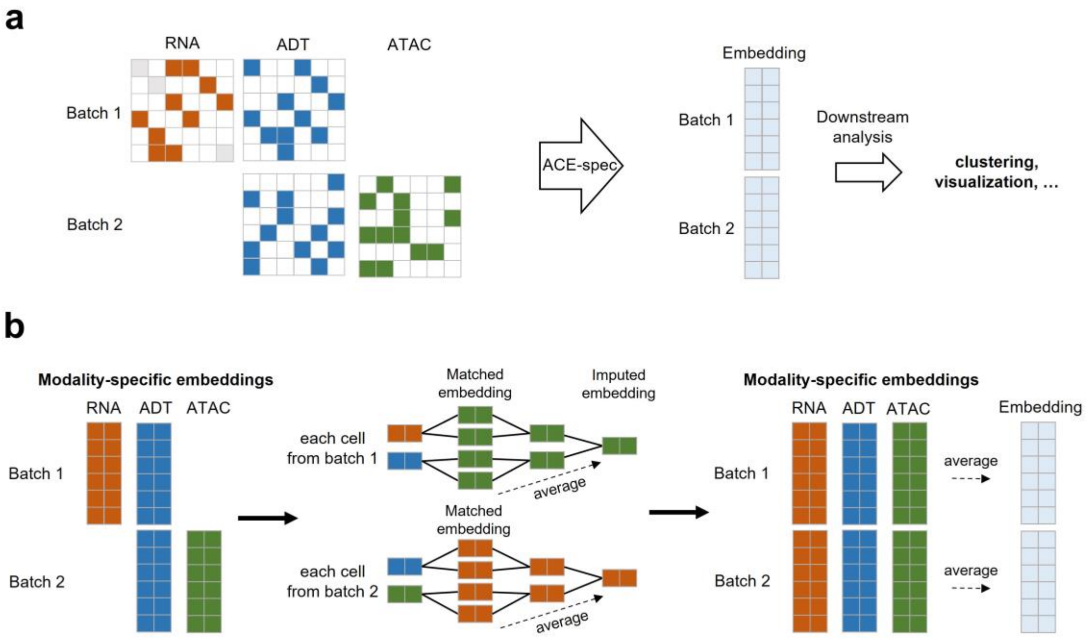


Supplementary Fig. S2. Illustration of ACE-spec's workflow. (a) Input and output of ACE-spec. (b) The embedding imputation process of ACE-spec. ACE-spec imputes the missing modality-specific embeddings in each batch and then averages the embeddings from all modalities to get the final embedding for each batch.

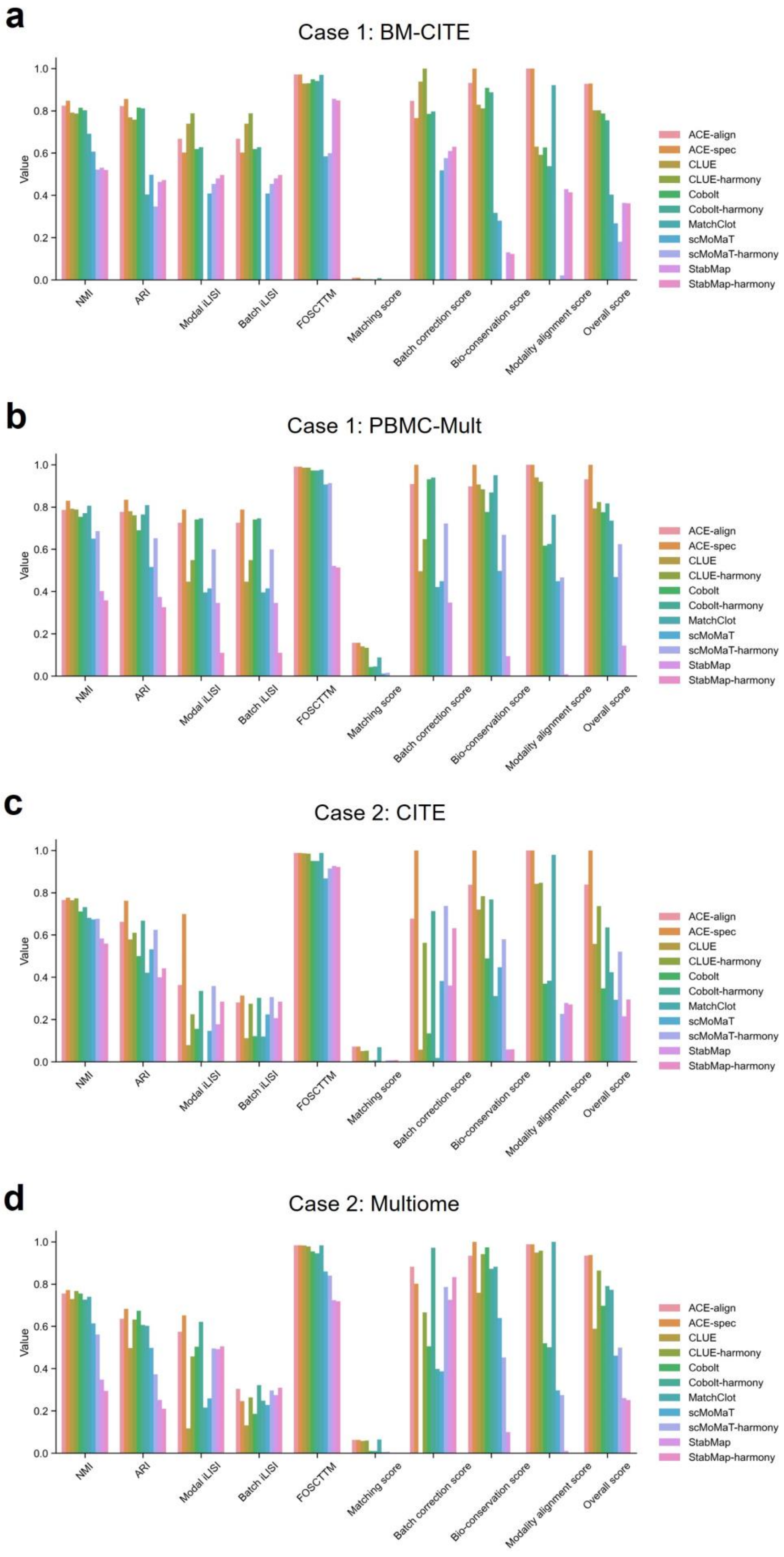


Supplementary Fig. S3. Bar plots of bi-modal benchmarking results. (a), (b), (c), (d) show the scores on the BM-CITE, PBMC-Mult, CITE and Multiome datasets, respectively.

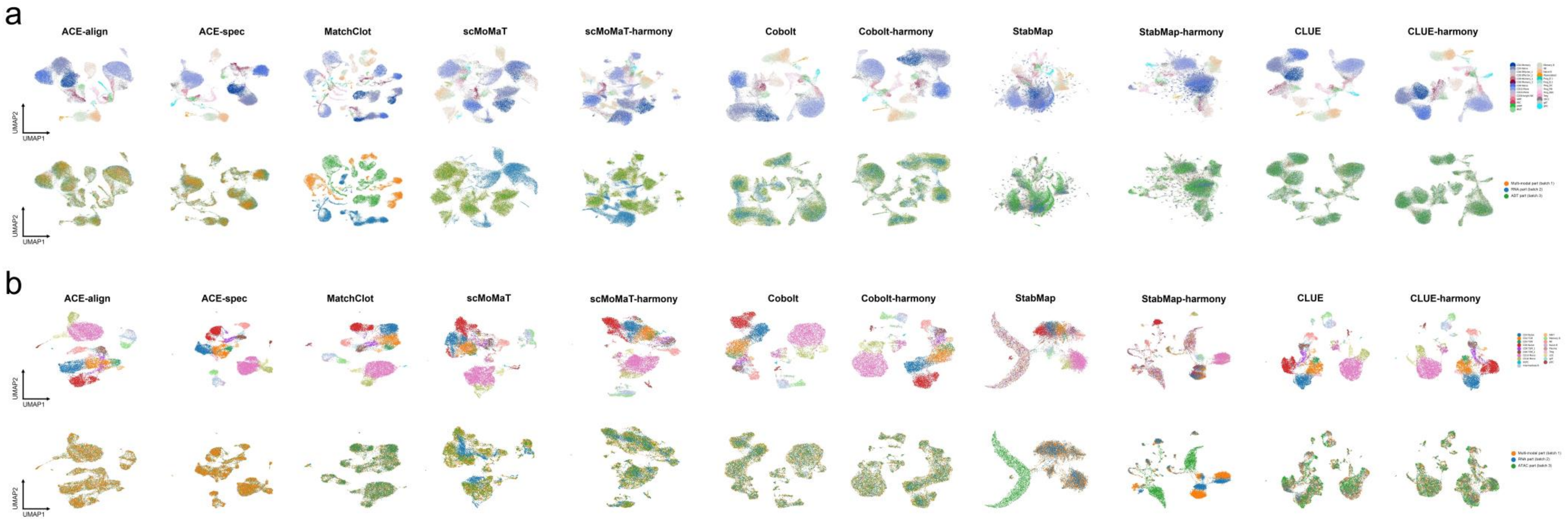


Supplementary Fig. S4. UMAP plots of embeddings in bi-modal case 1 from all compared methods. In each panel, cells in the first row are colored by cell types and colored by modal labels (batch labels) in the second row. (a) UMAP plots on the BM-CITE datasets. (b) UMAP plots on the PBMC-Mult dataset.

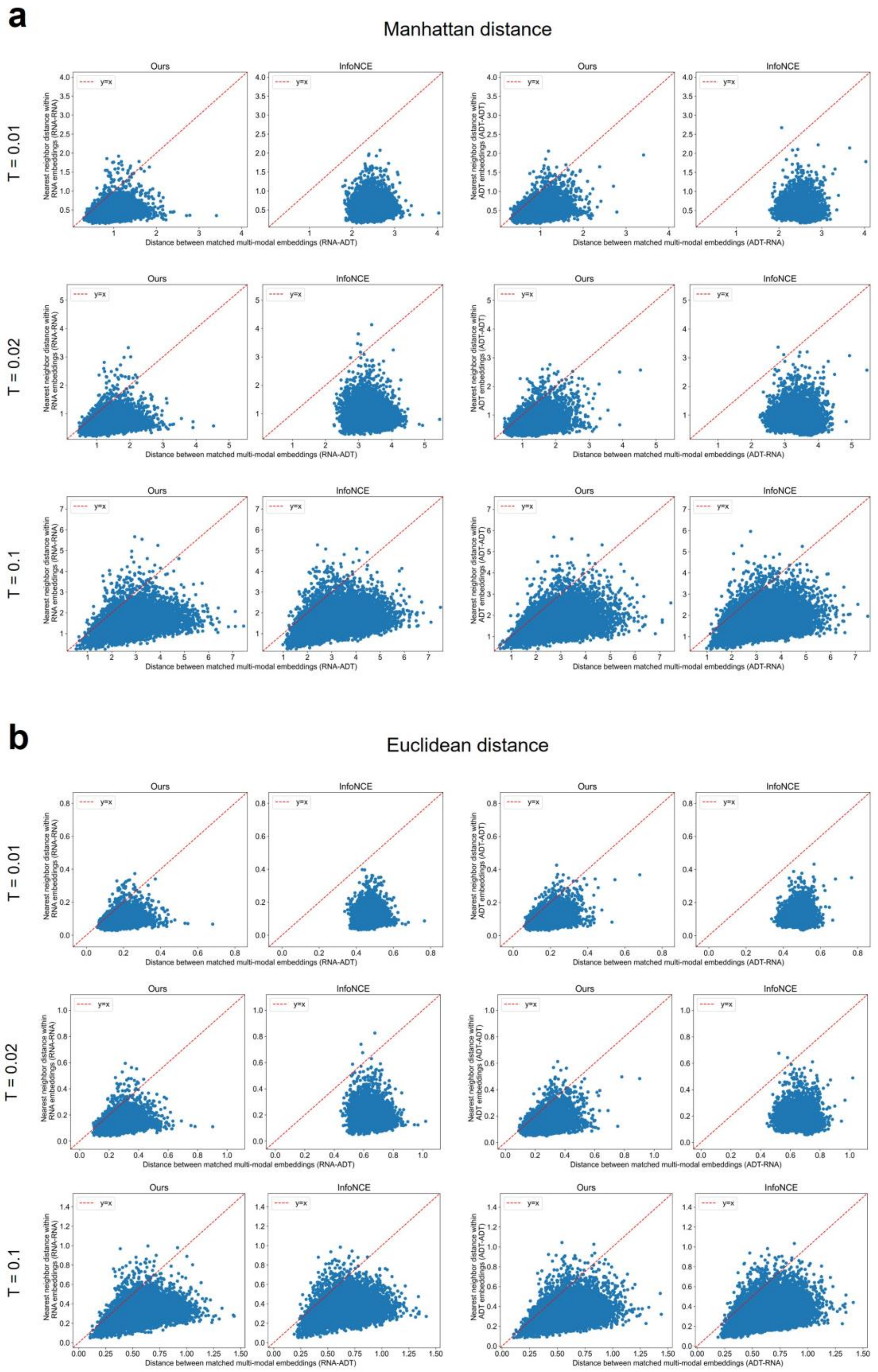


Supplementary Fig. S5. Analysis of modality gap phenomena. (a) For each cell, Manhattan distances were calculated to the nearest neighbor within the same modality (intra-modality) and to its paired embedding in the other modality (inter-modality), using embeddings derived from InfoNCE and our proposed loss functions. (b) For each cell, Euclidean distances were calculated to the nearest neighbor within the same modality (intra-modality) and to its paired embedding in the other modality (inter-modality), using embeddings derived from InfoNCE and our proposed loss functions.

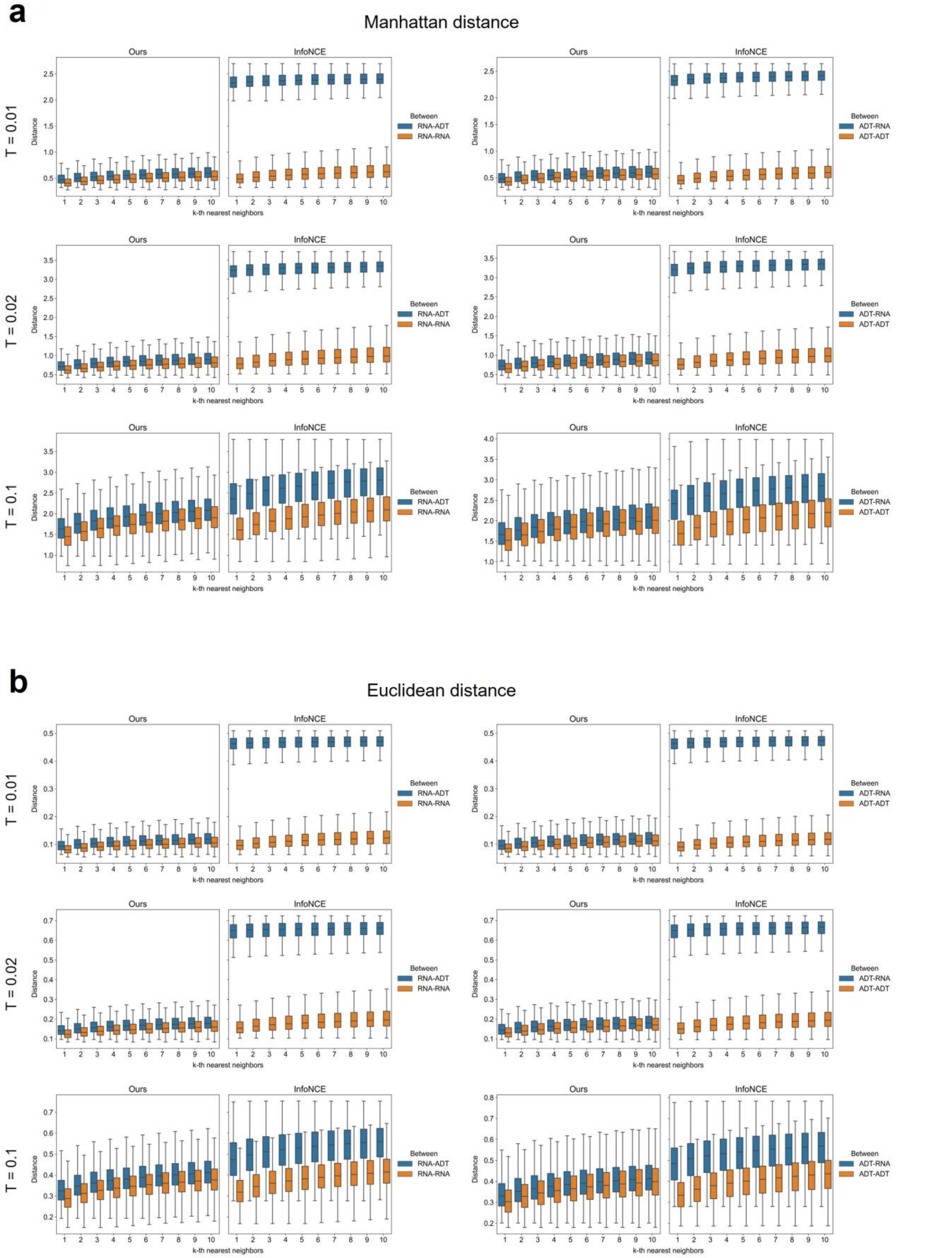


Supplementary Fig. S6. Analysis of modality gap phenomena within a larger area. (a) For each cell, Manhattan distances to the top 10 nearest neighbors within intra- and inter-modality were calculated based on embeddings derived from InfoNCE and our loss function. (b) For each cell, Euclidean distances to the top 10 nearest neighbors within intra- and inter-modality were calculated based on embeddings derived from InfoNCE and our loss function.

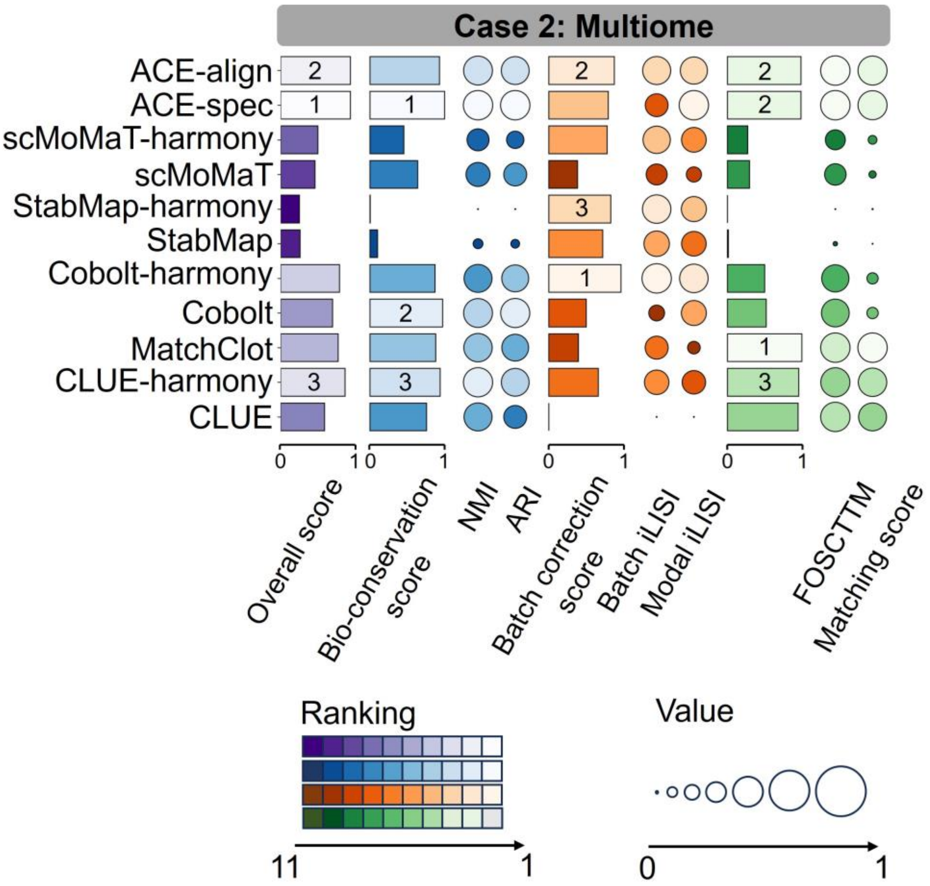


Supplementary Fig. S7. Mosaic integration benchmark on Multiome dataset of bi-modal case 2. We labeled the top three methods for each score.

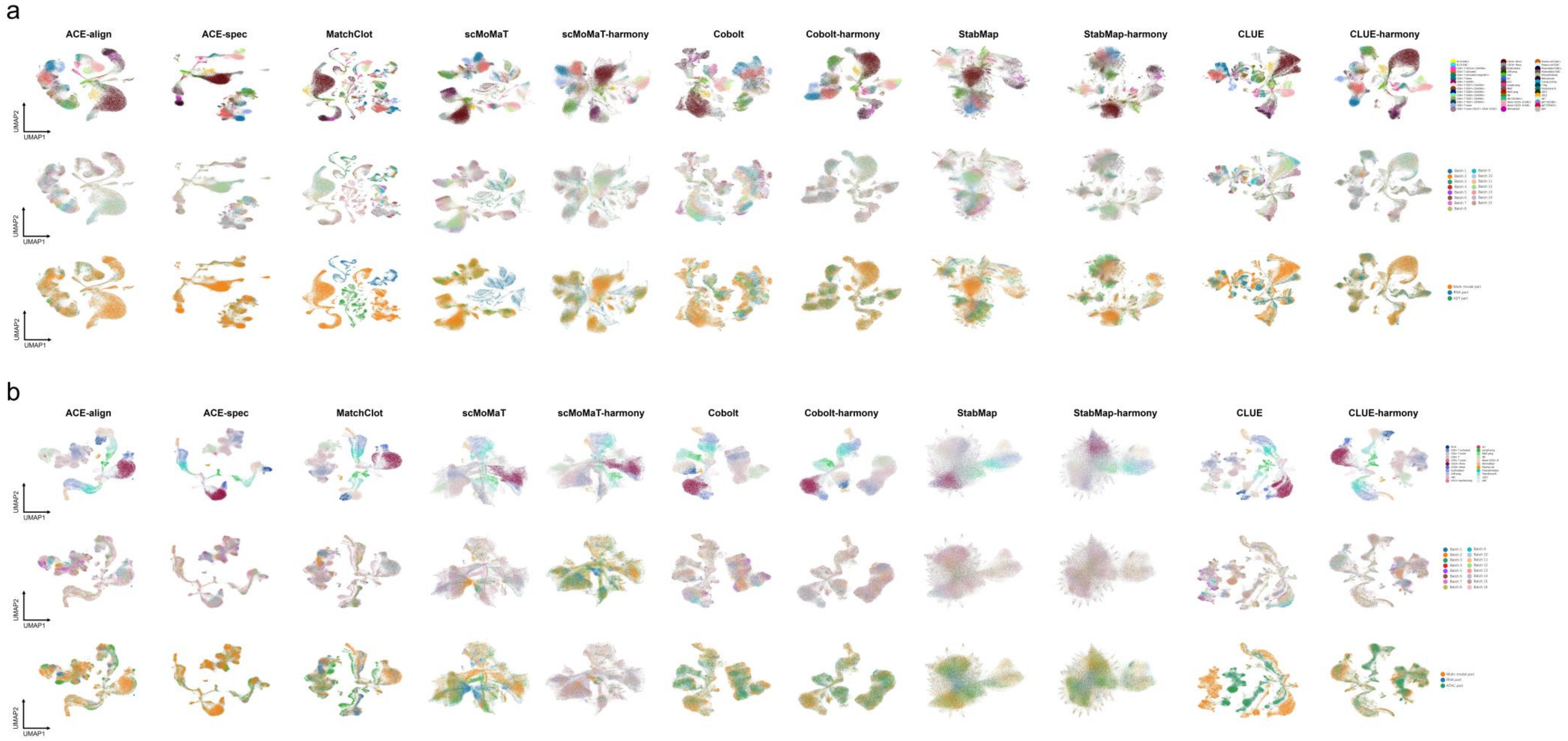


Supplementary Fig. S8. UMAP plots of embeddings in bi-modal case 2 from all compared methods. In each panel, cells in the first row are colored by cell types, colored by batch labels in the second row, and colored by modal labels in the third row. (a) UMAP plots on the CITE dataset. (b) UMAP plots on the Multiome dataset.

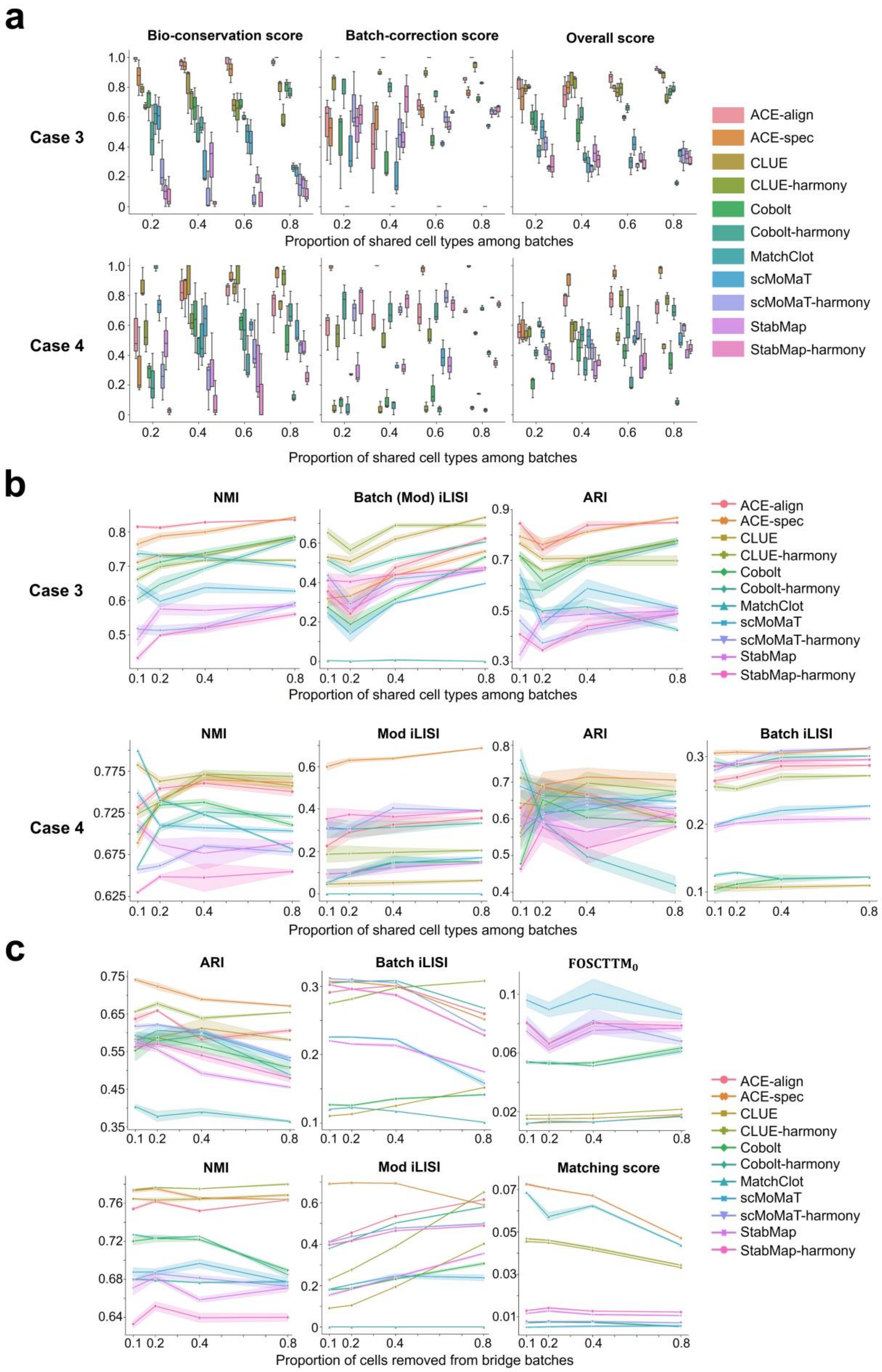


Supplementary Fig. S9. Specific metrics and scores in bi-modal mosaic integration benchmark. (a) Bio-conservation scores, batch-correction scores, and overall scores in case 3 and case 4. (b) Specific metrics in case 3 and case 4. They are not min-max scaled. (c) Specific metrics in experiment of removing cells from bridge batches. They are not min-max scaled.

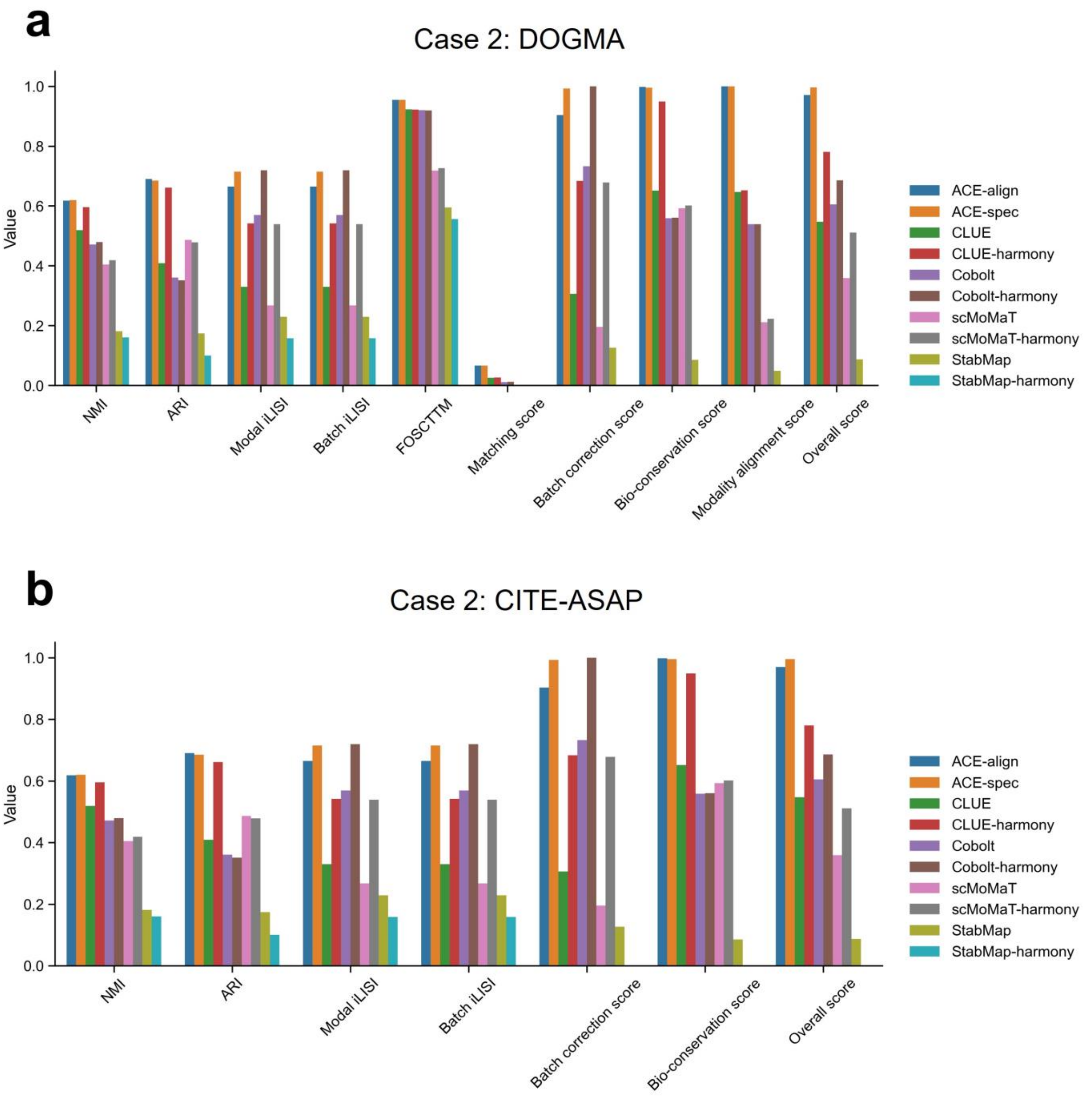


Supplementary Fig. S10. Bar plots of tri-modal benchmarking results. (a), (b) show the scores on the DOGMA and CITE-ASAP datasets, respectively.

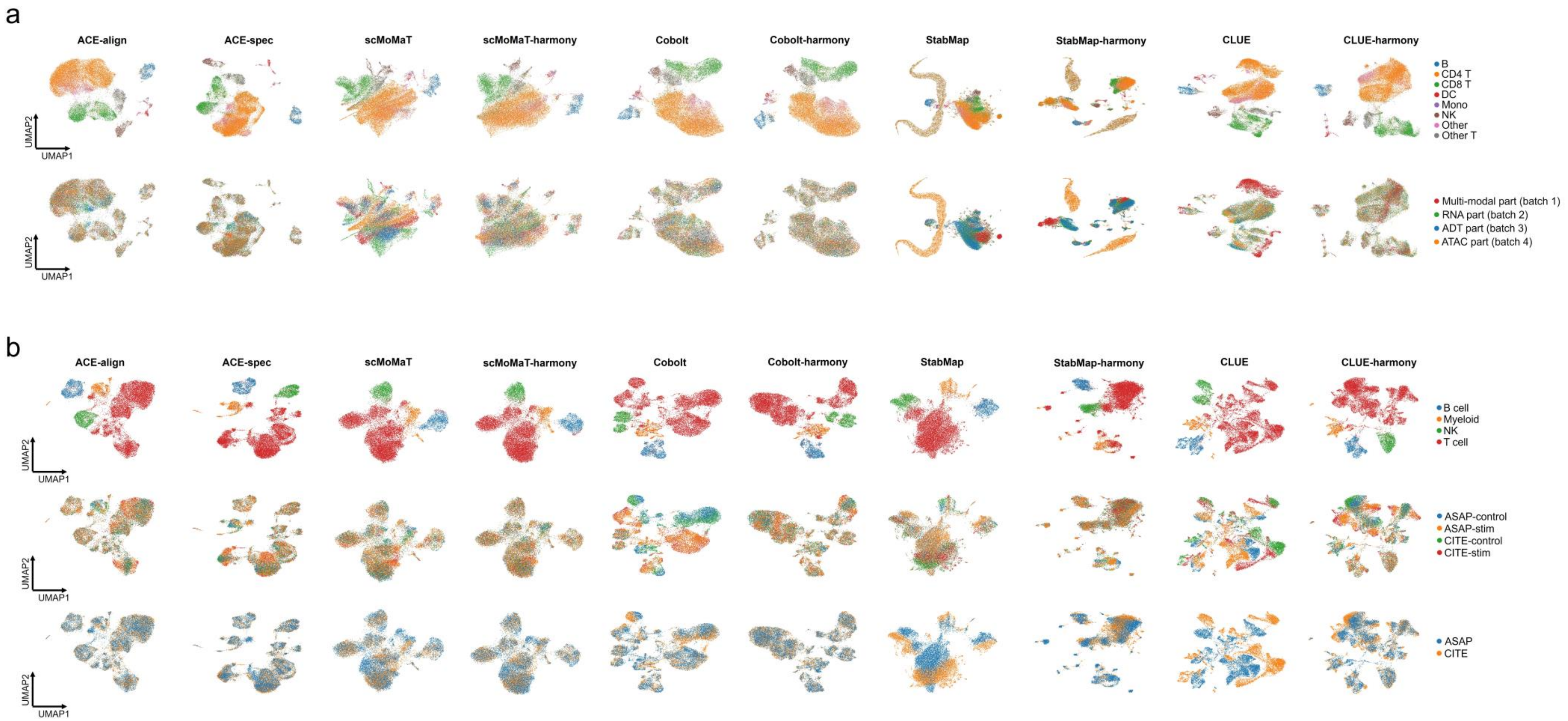


Supplementary Fig. S11. UMAP plots of embeddings in tri-modal integration tasks from all compared methods. (a) UMAP plots on the DOGMA dataset. Cells in the first row are colored by cell types, colored by modal (batch) labels in the second row. (b) UMAP plots on the Multiome dataset. Cells in the first row are colored by cell types, colored by batch labels in the second row, and colored by modal labels in the third row.

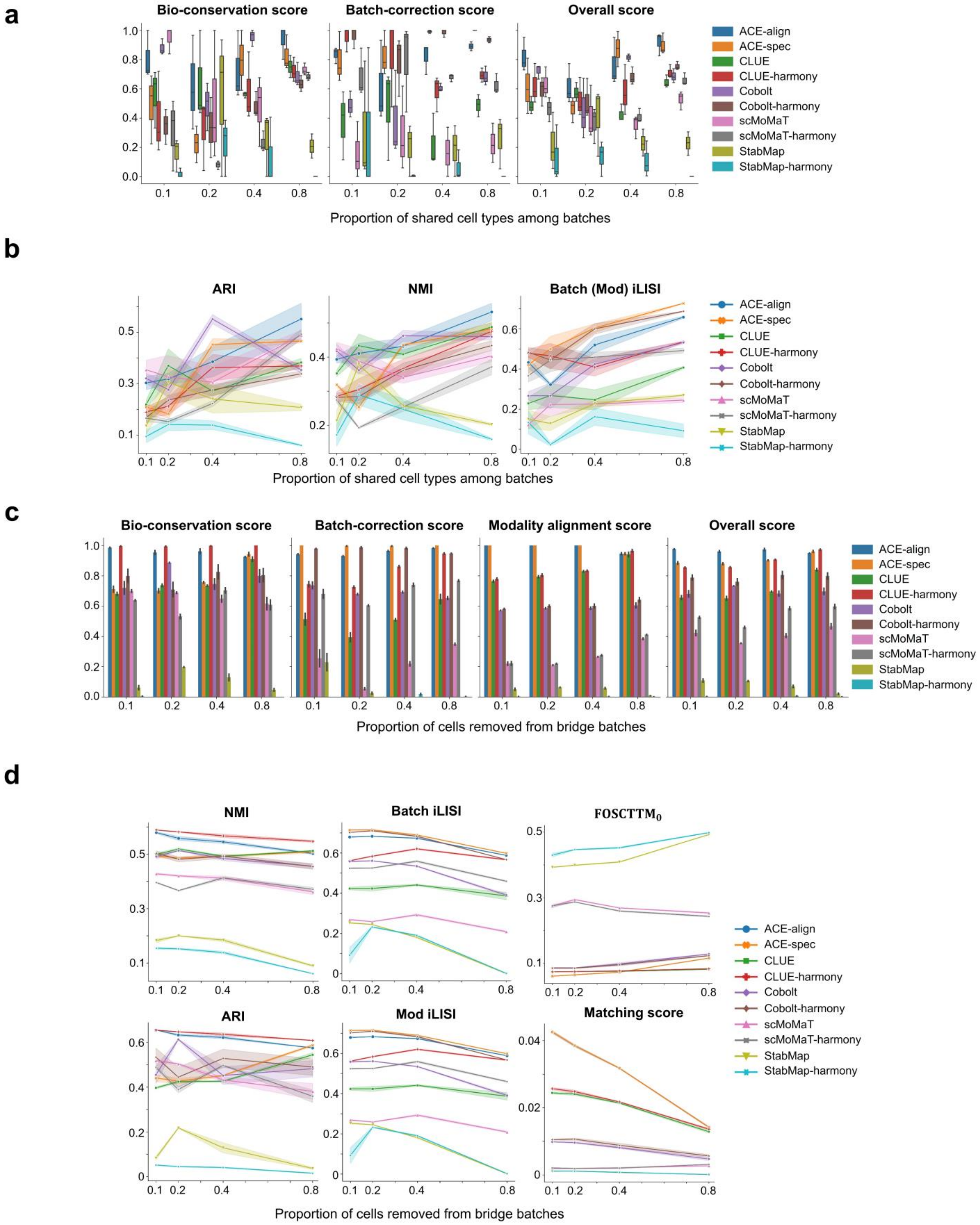


Supplementary Fig. S12. Specific metrics and scores in tri-modal mosaic integration benchmark. (a) Bio-conservation scores, batch-correction scores, and overall scores in tri-modal case 4. (b) Specific metrics in tri-modal case 4. They are not min-max scaled. (c) All scores in experiment of removing cells from bridge batches. (d) Specific metrics in experiment of removing cells from bridge batches. They are not min-max scaled.

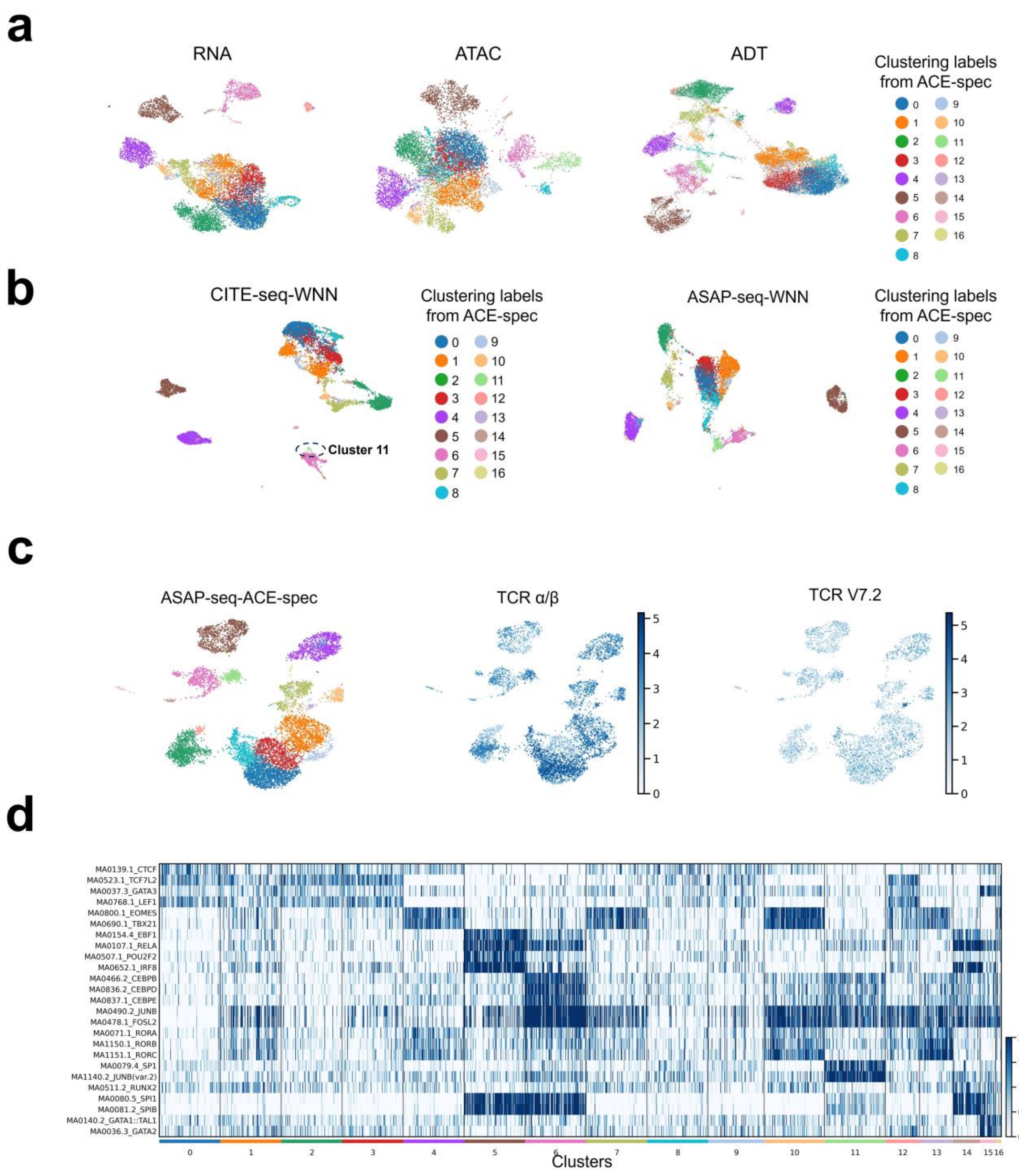


Supplementary Fig. S13. Analysis of ACE-spec's results on the CITE-ASAP dataset. (a) UMAP plots based on raw modality profiles. Cells are colored by clustering labels from ACE-spec. (b) UMAP plots of WNN results on the batches measured with CITE-seq and ASAP-seq. Cells are colored by clustering labels from ACE-spec. (c) UMAP plots of ACE-spec's embedding on the batches measured with ASAP-seq. In the left, cells are colored by the clustering labels; heatmap displaying the expression level of surface protein TCR Vα7.2 in these cells. (d) Activity heatmap of known marker motifs: NK: EOMES, TBX21 [1,2]; B: SPI1, EBF1, IRF8 [1,3]; Naïve CD4 T: CTCF, TCF7L2 [1,4]; CD4+ Memory T: GATA3 [5]; T regulatory (Treg): JUNB, FOSL2 [1,6]; Naïve CD8 T: CTCF, TCF7L2, LEF1 [1]; CD8+ memory T: TBX21, EOMES [7]; MAIT: RORA, RORB, RORC [8]; gdT: RORC, RORA, RORC, TBX21, EOMES [9]; CD14 Monocytes: CEBPB, CEBPD, CEBPE, CEBPG [10]; cDC: SP1, JUNB [11]; pDC: RUNX2, SPI1, SPIB [12]; HSPC: GATA1::TAL1, GATA2, SPI1 [13].

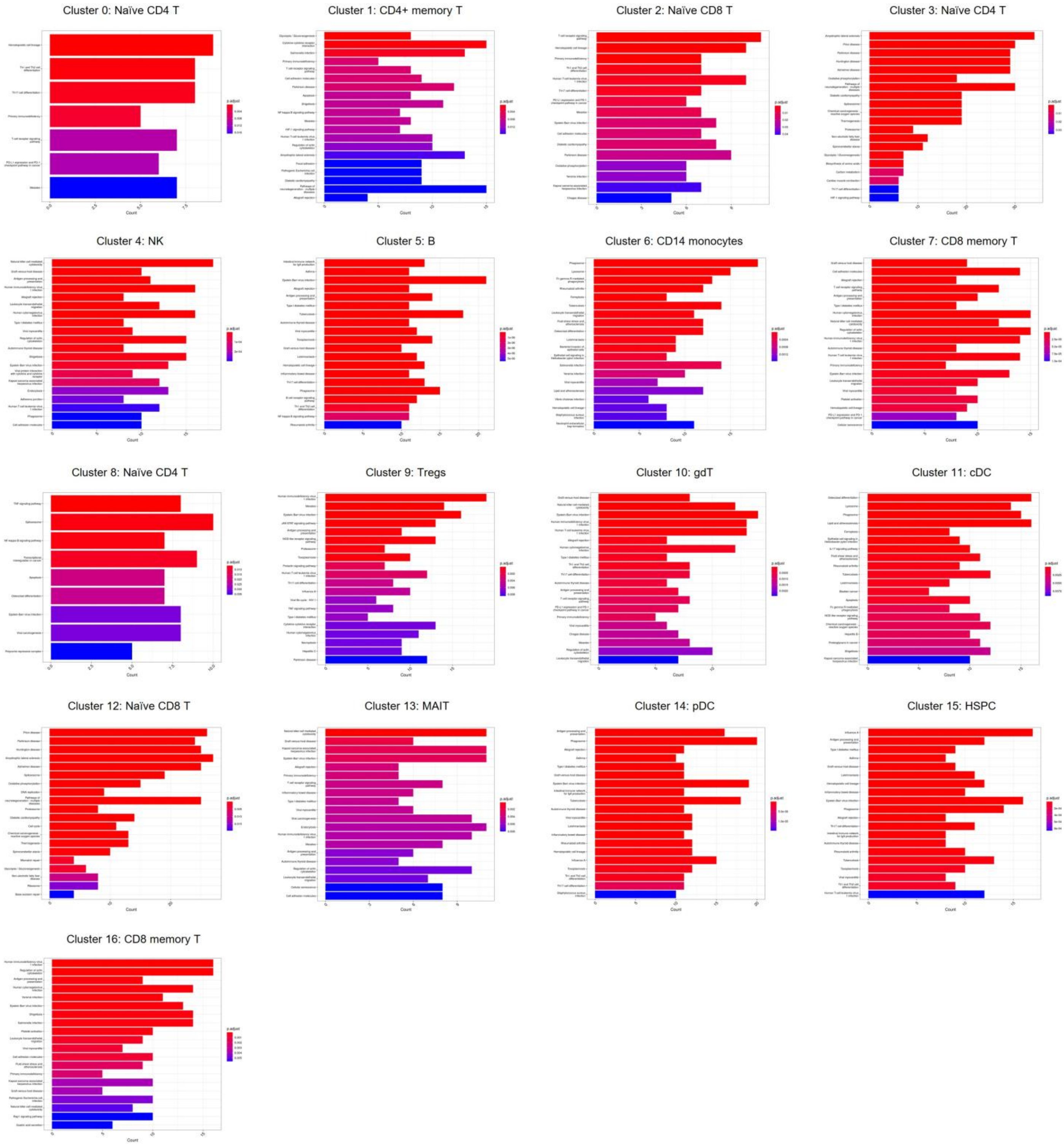


Supplementary Fig. S14. KEGG pathway enrichment analysis on the top 200 differentially expressed genes for each ACE-spec identified cluster (with our annotations).

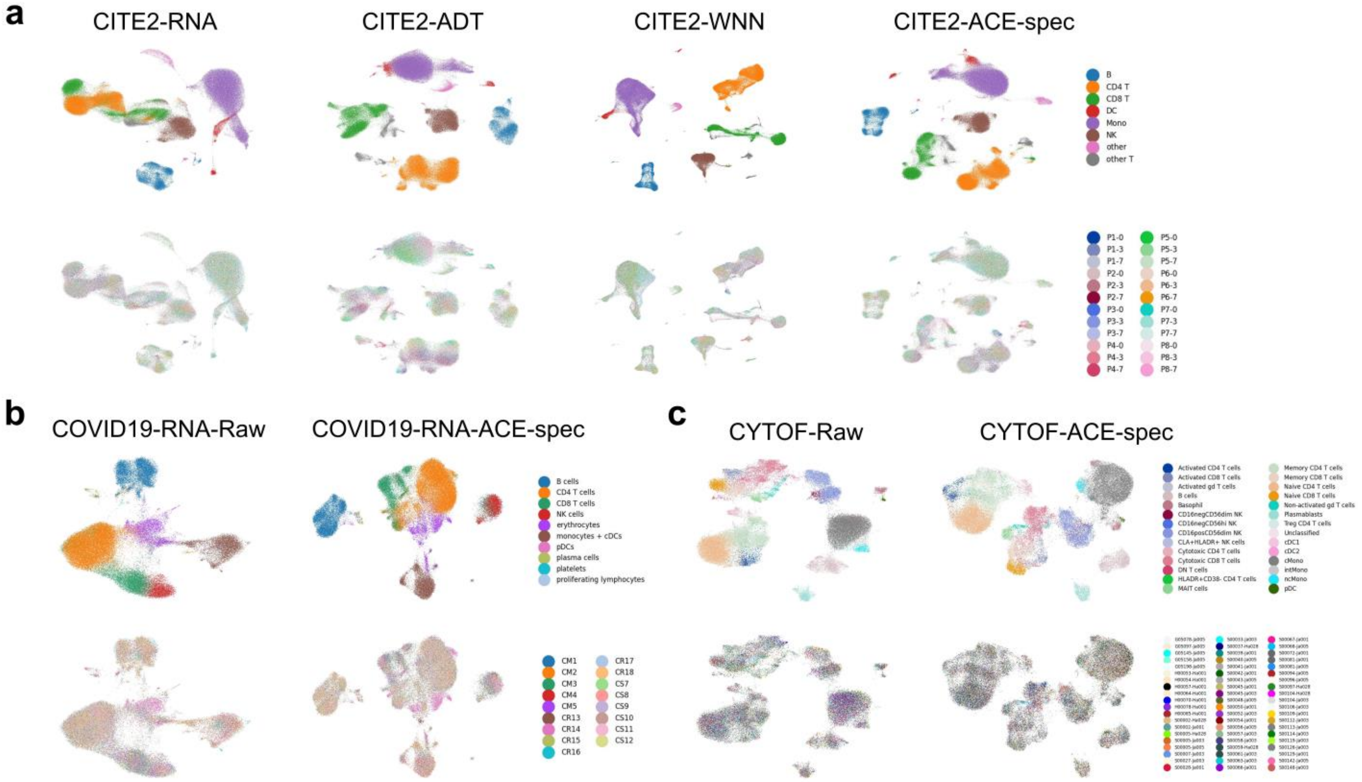


Supplementary Fig. S15. UMAP plots of raw CITE2, COVID19-RNA, CYTOF datasets and embeddings from ACE-spec. In each panel, cells are colored by cell type annotations in the first row and colored by batch labels in the second row. (a) UMAP visualizations of raw RNA, protein data of CITE2 dataset, WNN results, and embeddings from ACE-spec. (b) UMAP visualizations of raw COVID19-RNA dataset and embeddings from ACE-spec. (c) UMAP visualizations of raw CYTOF dataset and embeddings from ACE-spec.

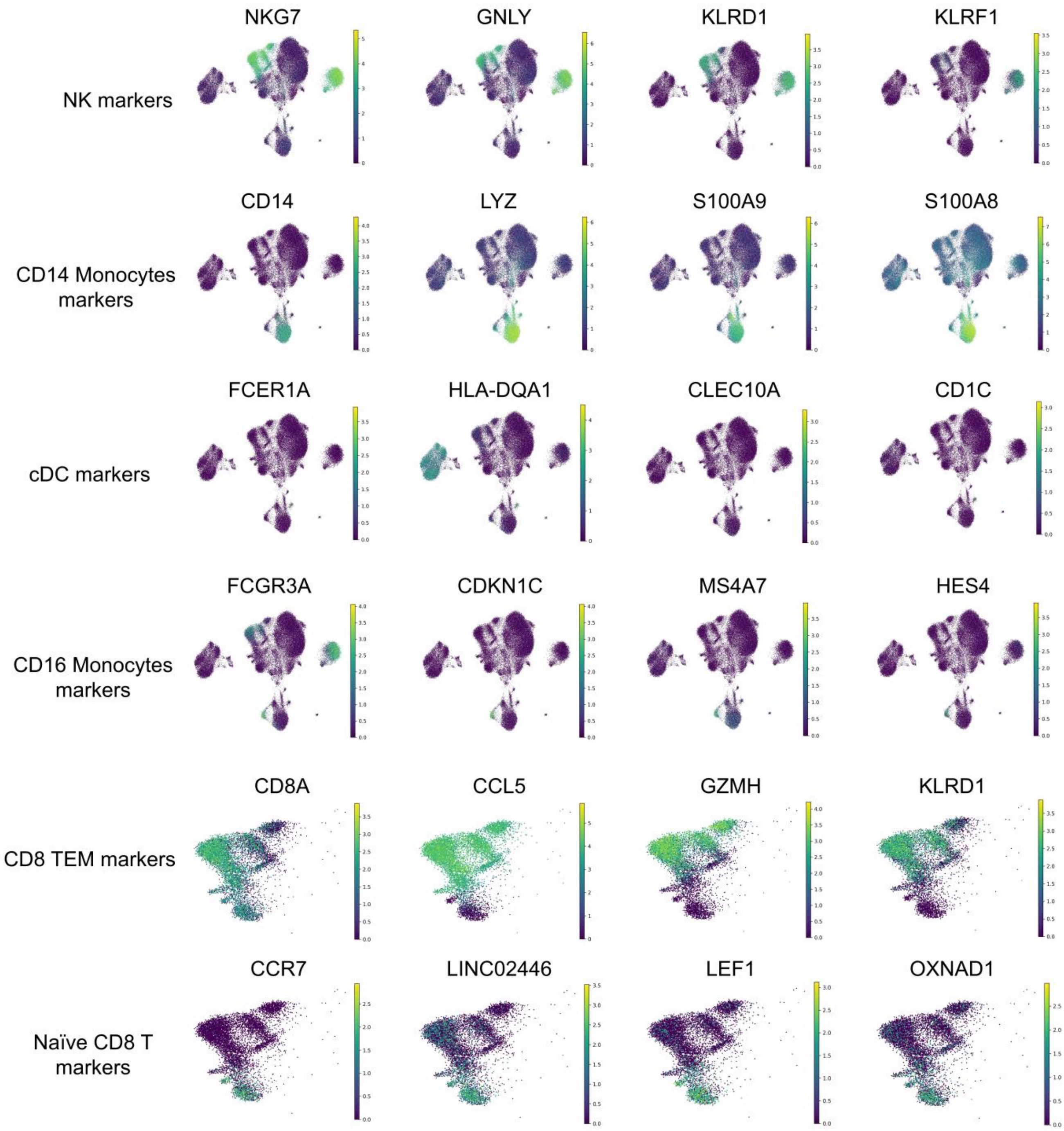


Supplementary Fig. S16. Expression heatmap of known markers genes in COVID19-RNA dataset.

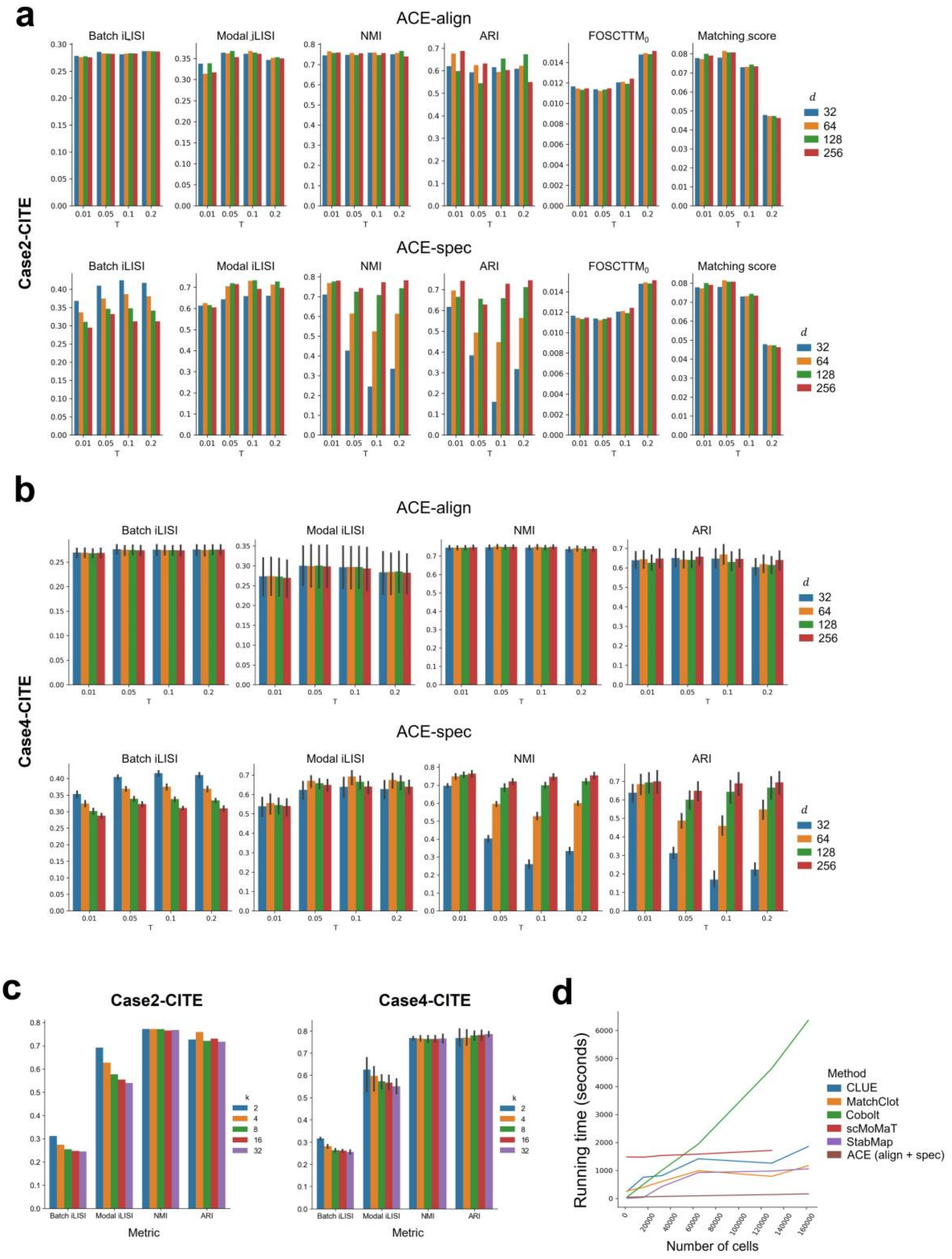


Supplementary Fig. S17. Parameter sensitivity experiment of ACE and comparison of time consumption among all methods. (a) ACE-align and ACE-spec's sensitivity to parameters $d$ and $\tau$ with respect to iLISI, NMI, ARI, $\mathrm{FOSCTTM}_0$, and matching scores in bi-modal case 2. (b) ACE-align and ACE-spec's sensitivity to parameters $d$ and $\tau$ with respect to iLISI, NMI, ARI in bi-modal case 4. (c) ACE-spec's sensitivity to parameter $k$. (d) Comparison of time consumption (seconds) of all methods on multiple subsampled CITE2 dataset. Note that the time consumed by data loading and preprocessing/post-processing steps such as normalization, dimension reduction and batch correction is not recorded.

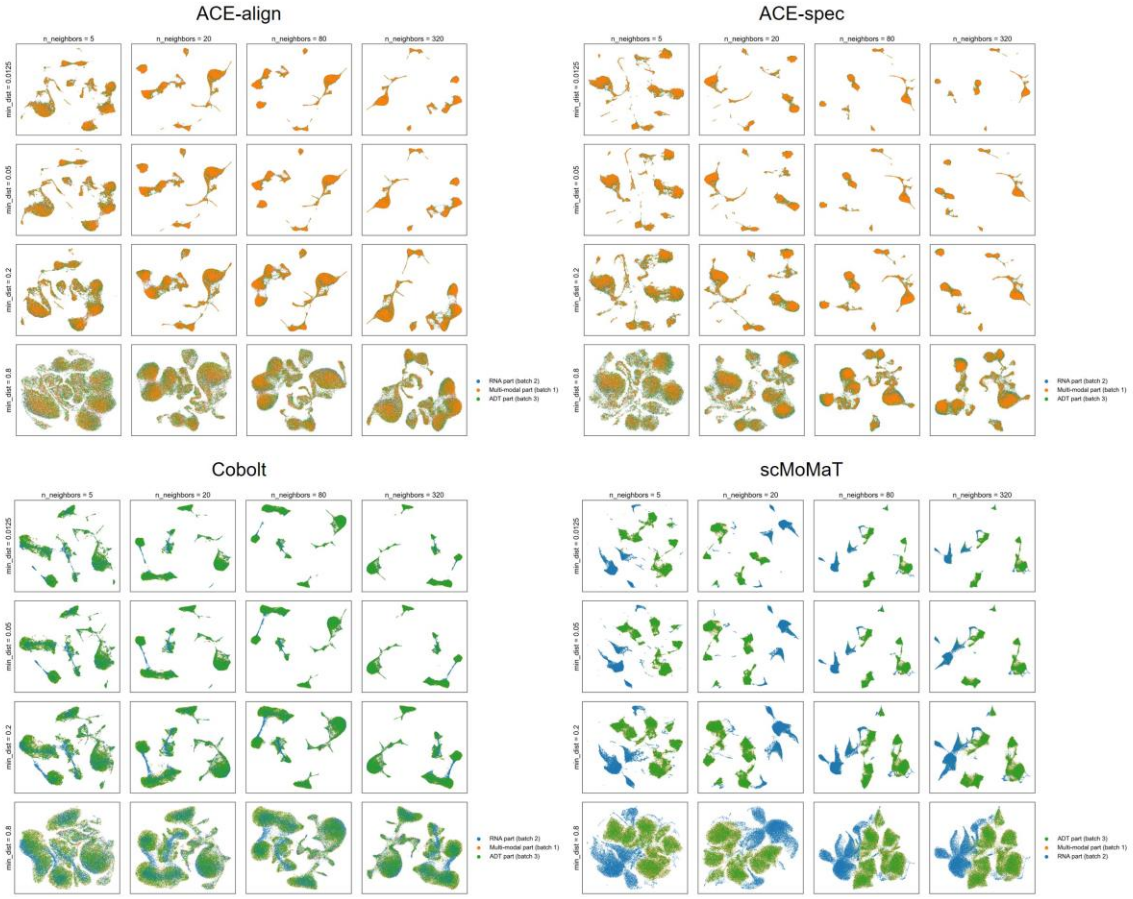


Supplementary Fig. S18. Varying UMAP hyperparameters, number of nearest neighbors and minimum distance, in visualizing ACE-align, ACE-spec, Cobolt and scMoMaT's embeddings on BM-CITE dataset. Cells are colored by batch labels.

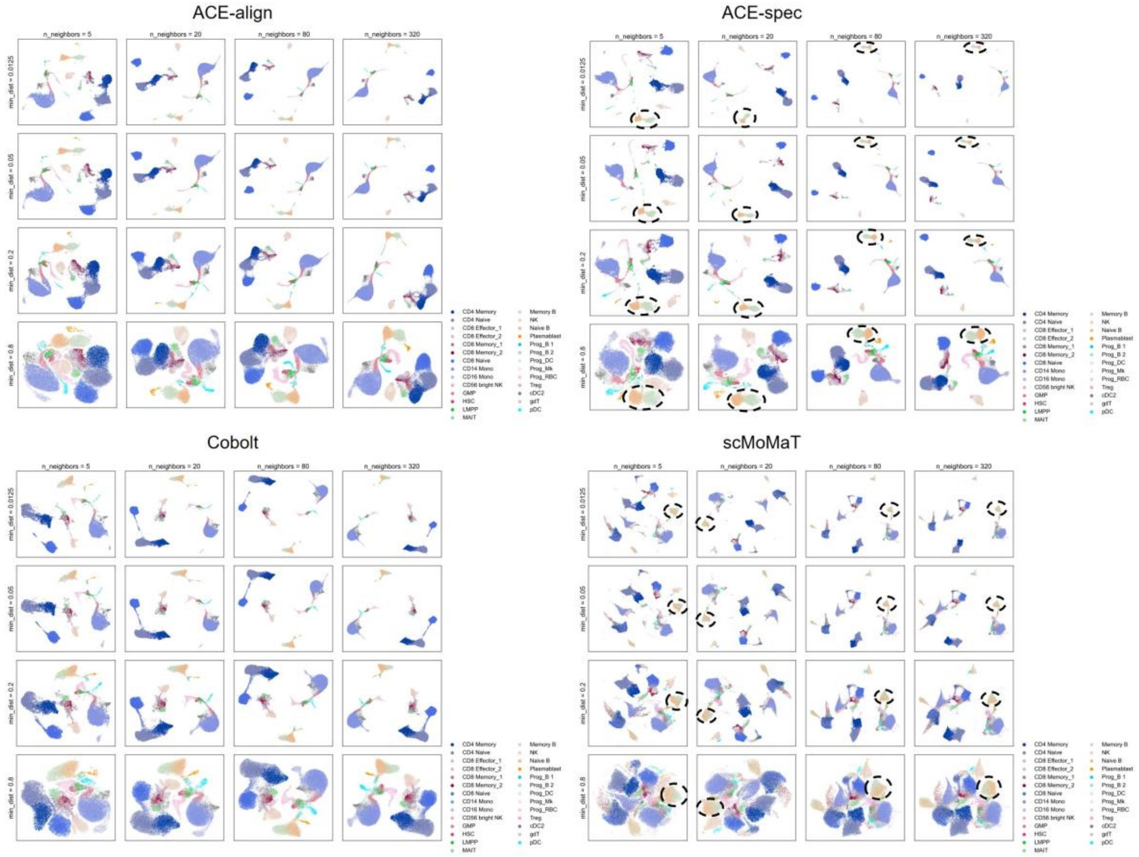


Supplementary Fig. S19. Varying UMAP hyperparameters, number of nearest neighbors and minimum distance, in visualizing ACE-align, ACE-spec, Cobolt and scMoMaT's embeddings on BM-CITE dataset. Cells are colored by cell type labels.

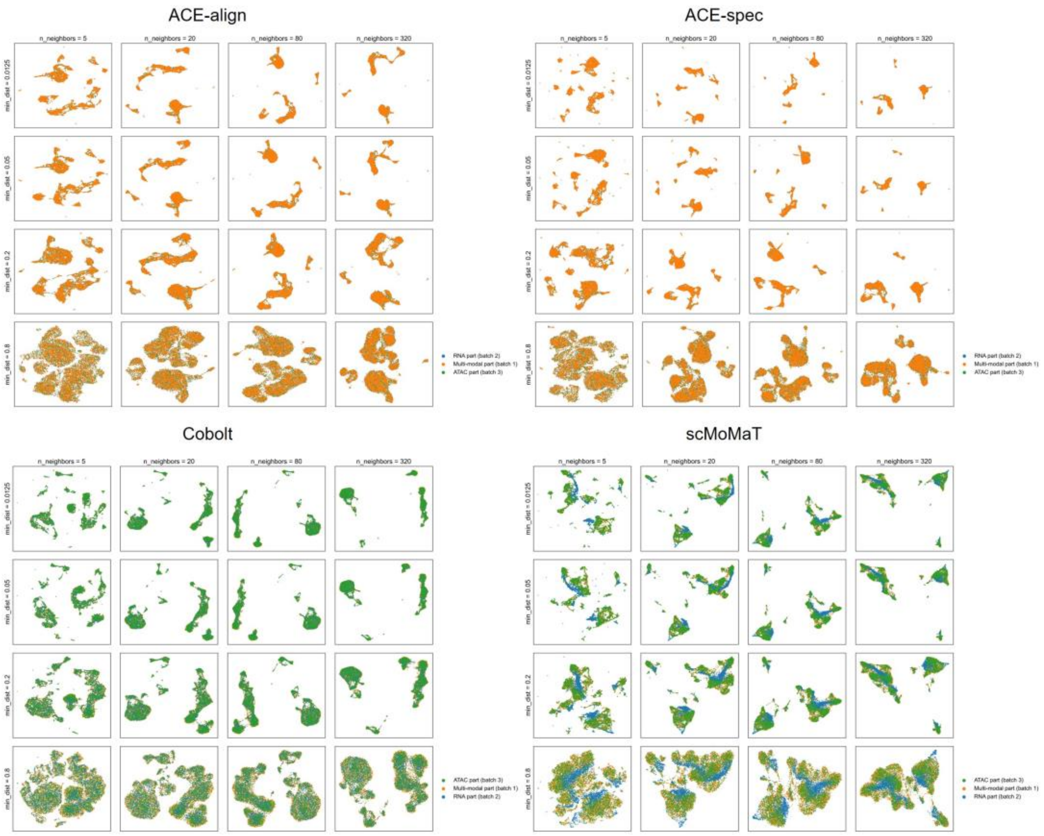


Supplementary Fig. S20. Varying UMAP hyperparameters, number of nearest neighbors and minimum distance, in visualizing ACE-align, ACE-spec, Cobolt and scMoMaT's embeddings on PBMC-Mult dataset. Cells are colored by batch labels.

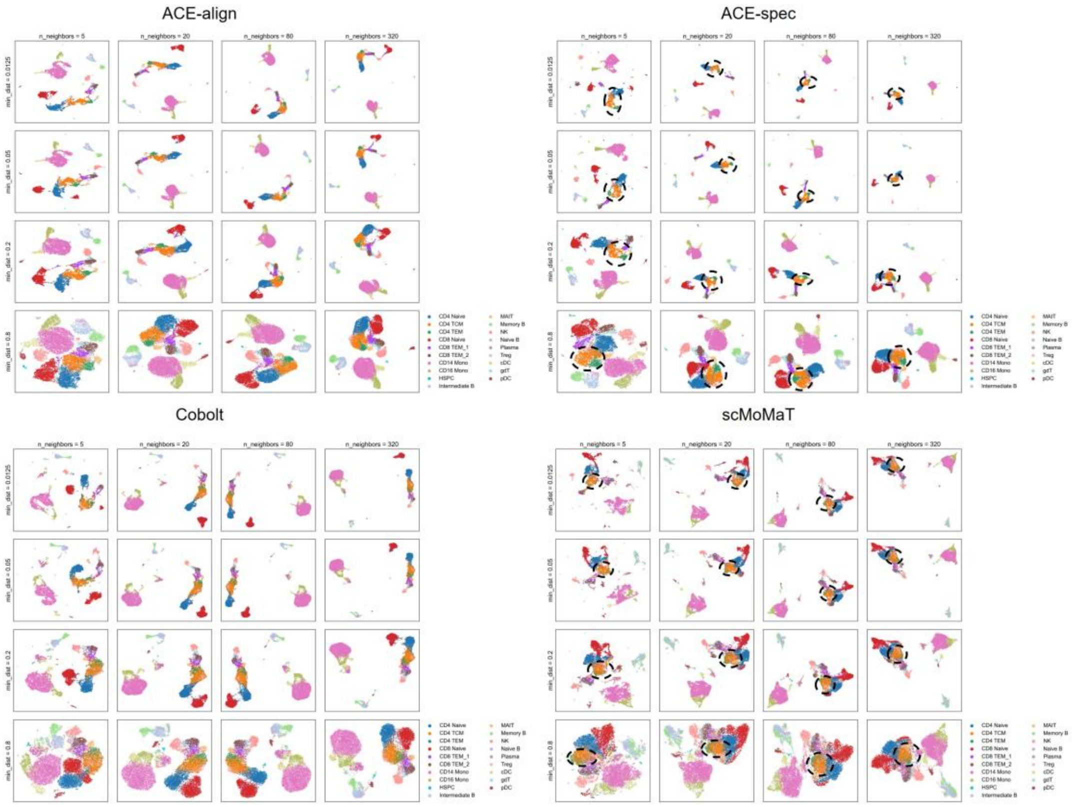


Supplementary Fig. S21. Varying UMAP hyperparameters, number of nearest neighbors and minimum distance, in visualizing ACE-align, ACE-spec, Cobolt and scMoMaT's embeddings on PBMC-Mult dataset. Cells are colored by cell type labels.

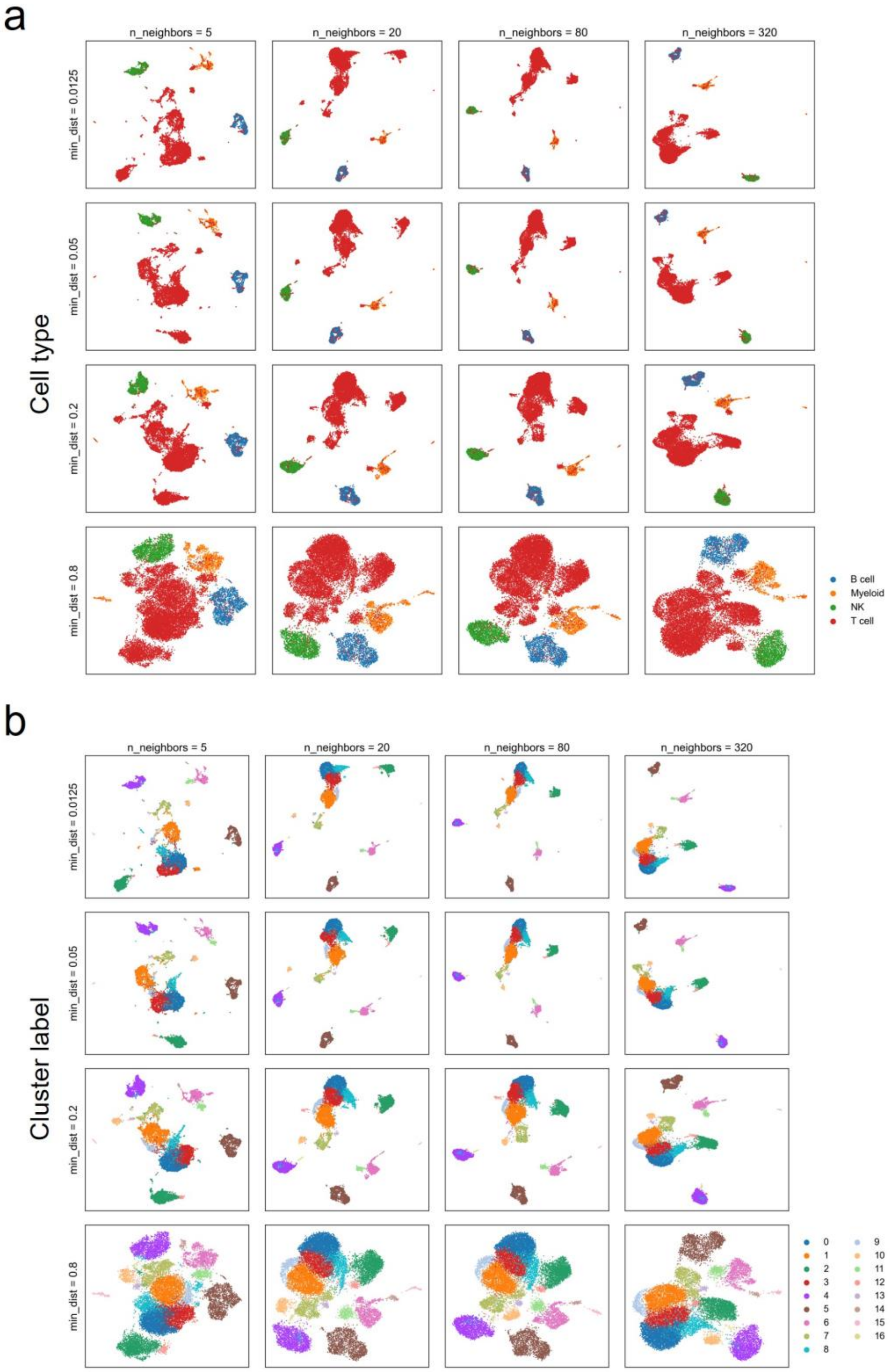


Supplementary Fig. S22. Varying UMAP hyperparameters, number of nearest neighbors and minimum distance, in visualizing ACE-spec's embeddings on CITE-ASAP dataset. Cells are colored by cell type and cluster labels in (a) and (b) respectively.

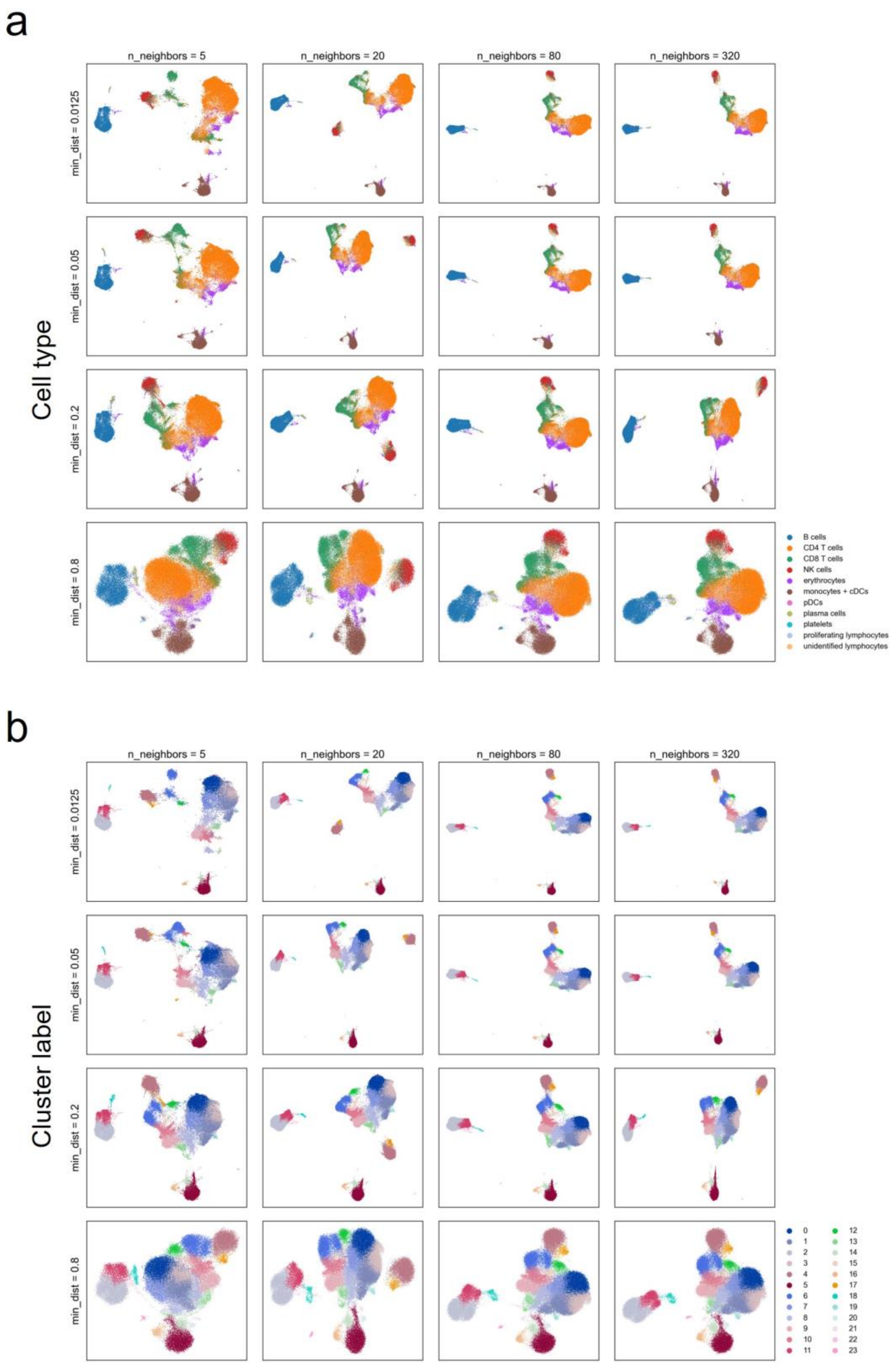


Supplementary Fig. S23. Varying UMAP hyperparameters, number of nearest neighbors and minimum distance, in visualizing ACE-spec's embeddings on COVID-19 dataset. Cells are colored by cell type and cluster labels in (a) and (b) respectively.

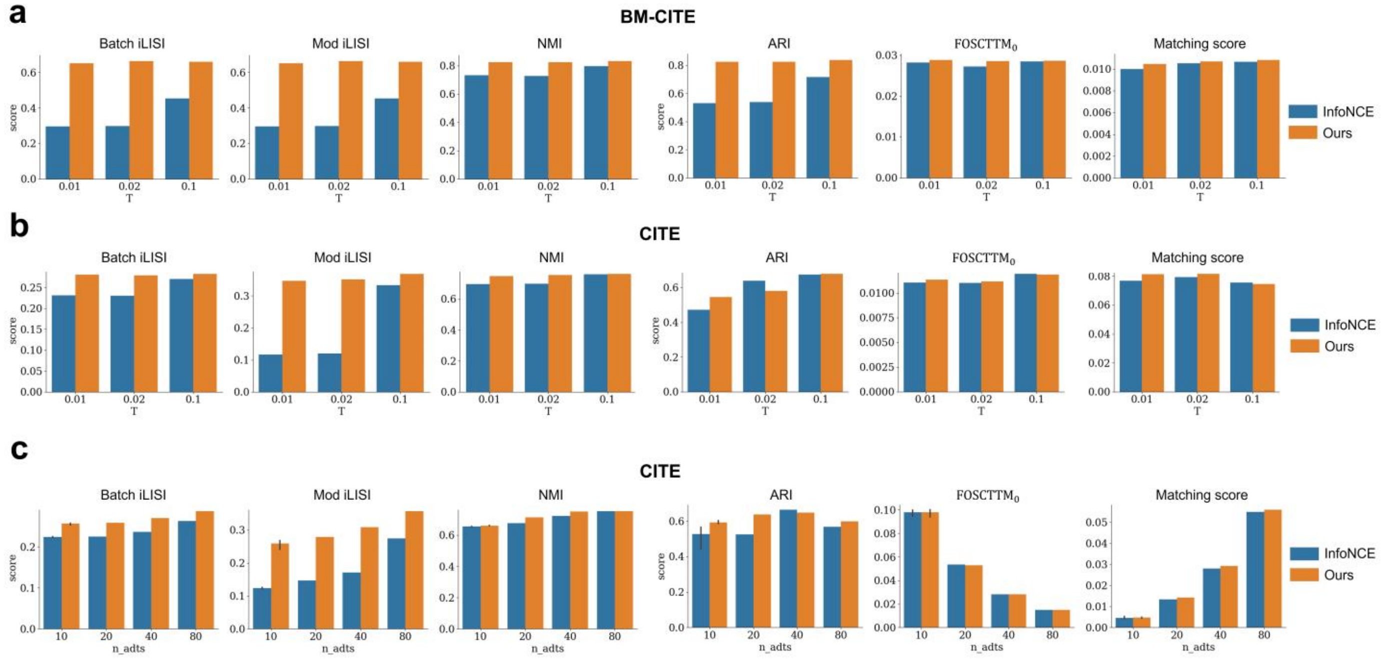


Supplementary Fig. S24. Ablation study on our proposed contrastive learning loss. (a) Comparison between our loss function and InfoNCE on BM-CITE dataset for different temperature value settings. (b) Comparison between our loss function and InfoNCE on CITE dataset for different temperature value settings. (c) Comparison between our loss function and InfoNCE on CITE dataset for different input numbers of protein features.

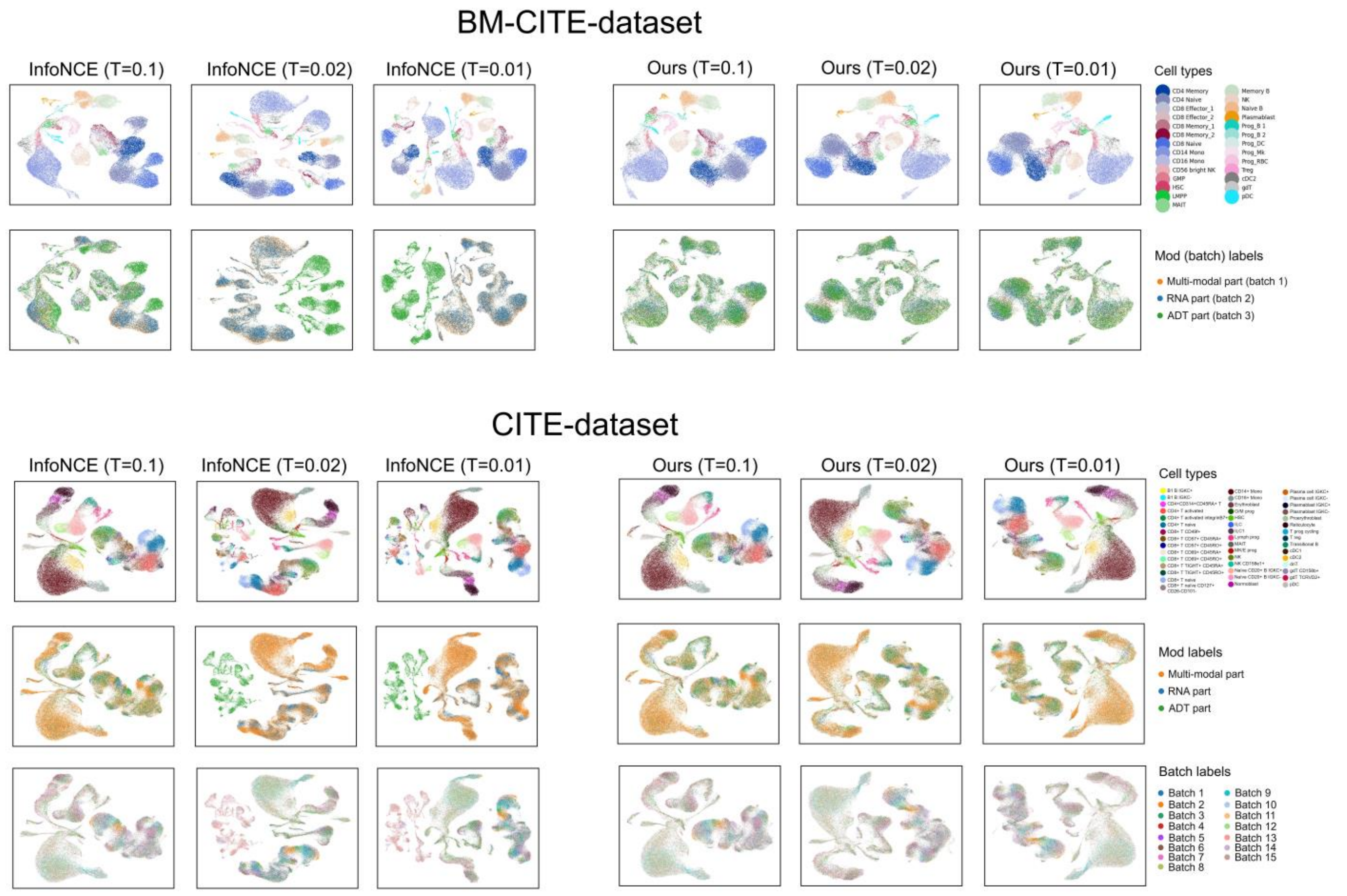


Supplementary Fig. S25. UMAP visualizations of embedding on BM-CITE and CITE datasets, derived from our proposed loss function and InfoNCE with different temperature value settings. For BM-CITE dataset, cells are colored by cell types in the first row and colored by modal (batch) labels in the second row. For CITE dataset, cells are colored by cell types in the first row, colored by modal labels in the second row, and colored by batch labels in the third row.

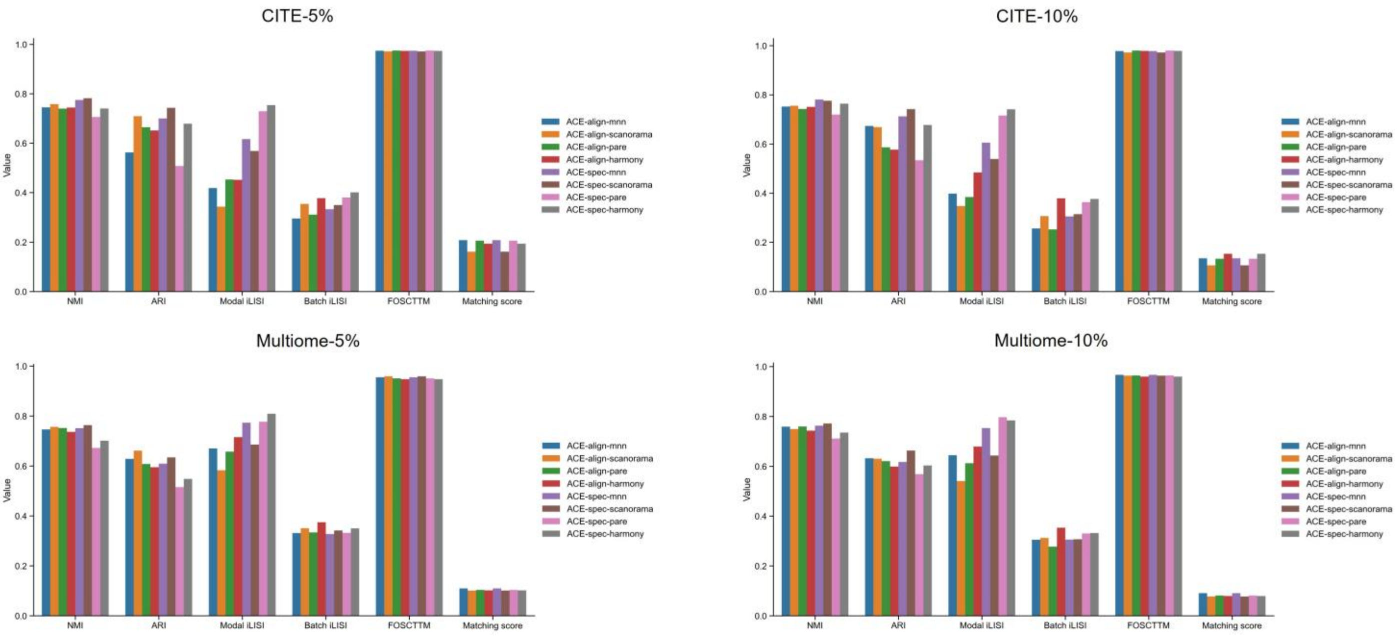


Supplementary Fig. S26. Evaluation of the impact of four batch correction methods on the performance of ACE-align and ACE-spec in CITE-5%/10% and Multiome-5%/10% datasets. The terms “-5%” and “-10%” denote subsampled datasets with sampling ratios of 5% and 10%, respectively.

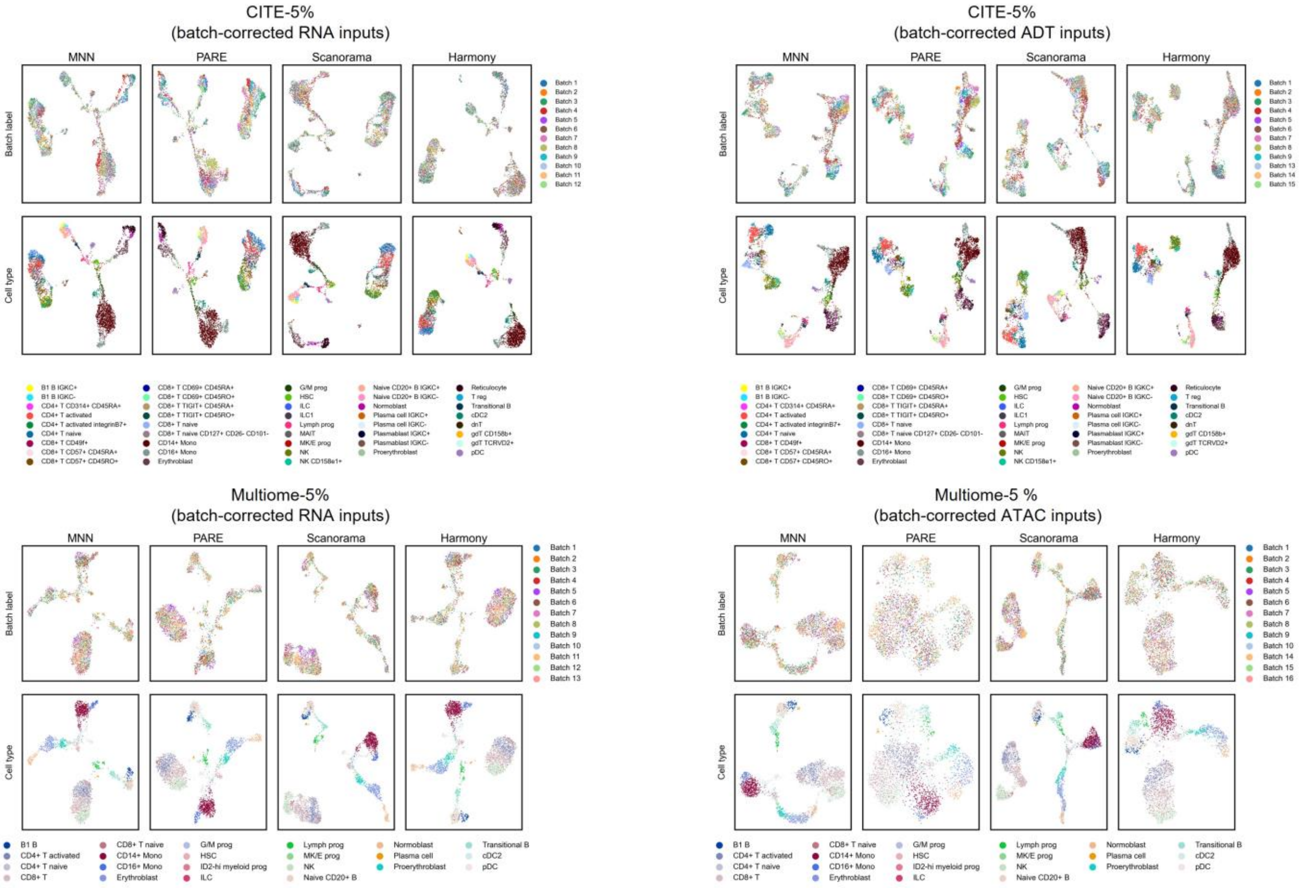


Supplementary Fig. S27. UMAP visualizations of batch corrected modality inputs processed from four batch correction methods. The terms “-5%” and “-10%” denote subsampled datasets with sampling ratios of 5% and 10%, respectively.

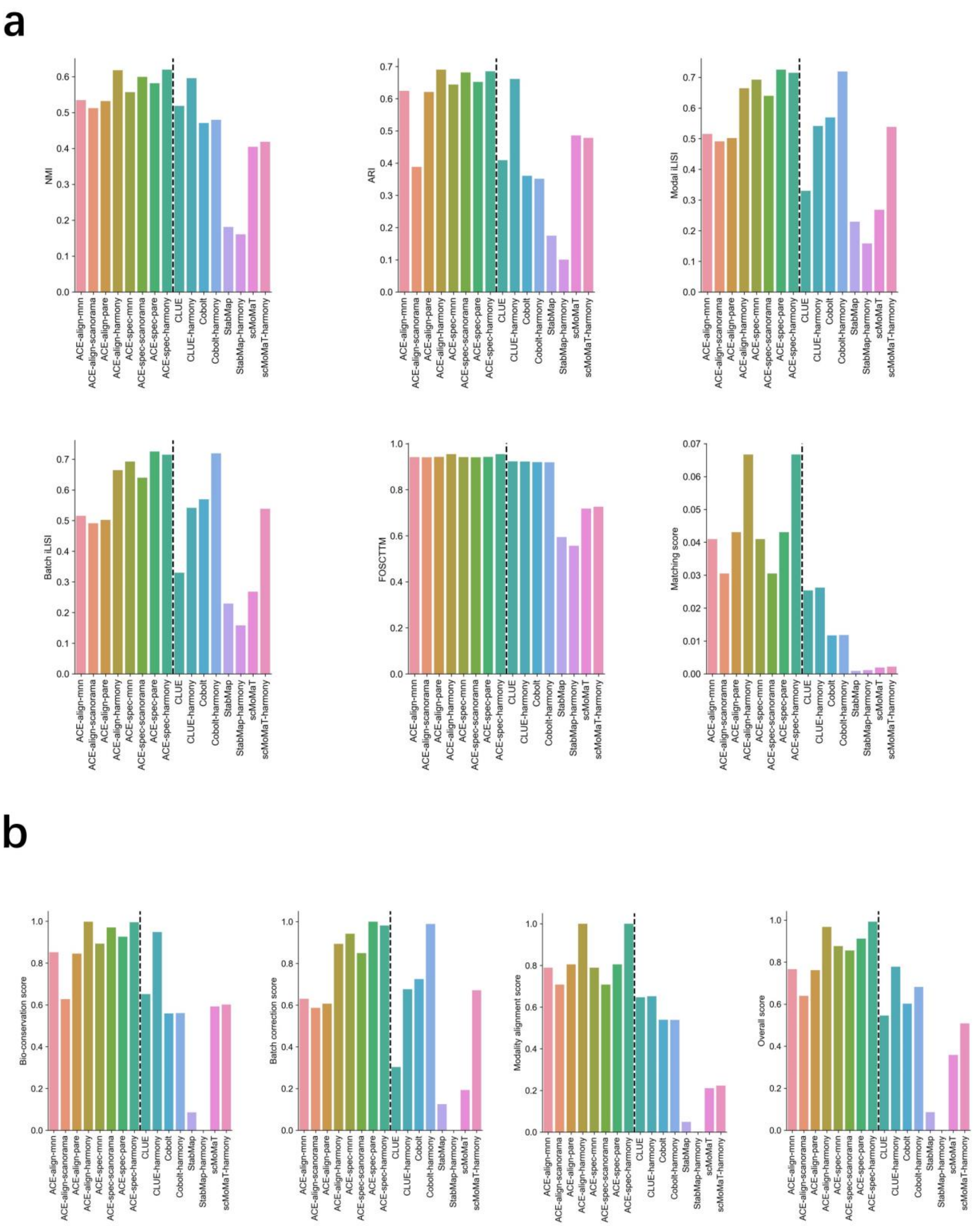


Supplementary Fig. S28. Evaluation of the impact of four batch correction methods on the performance of ACE-align and ACE-spec in DOGMA dataset. (a) Values of specific evaluation metrics, including NMI, ARI, modal iLISI, batch iLISI, FOSCTTM and matching score. (b) Bio-conservation score, batch-correction score, modality-alignment score and overall score.

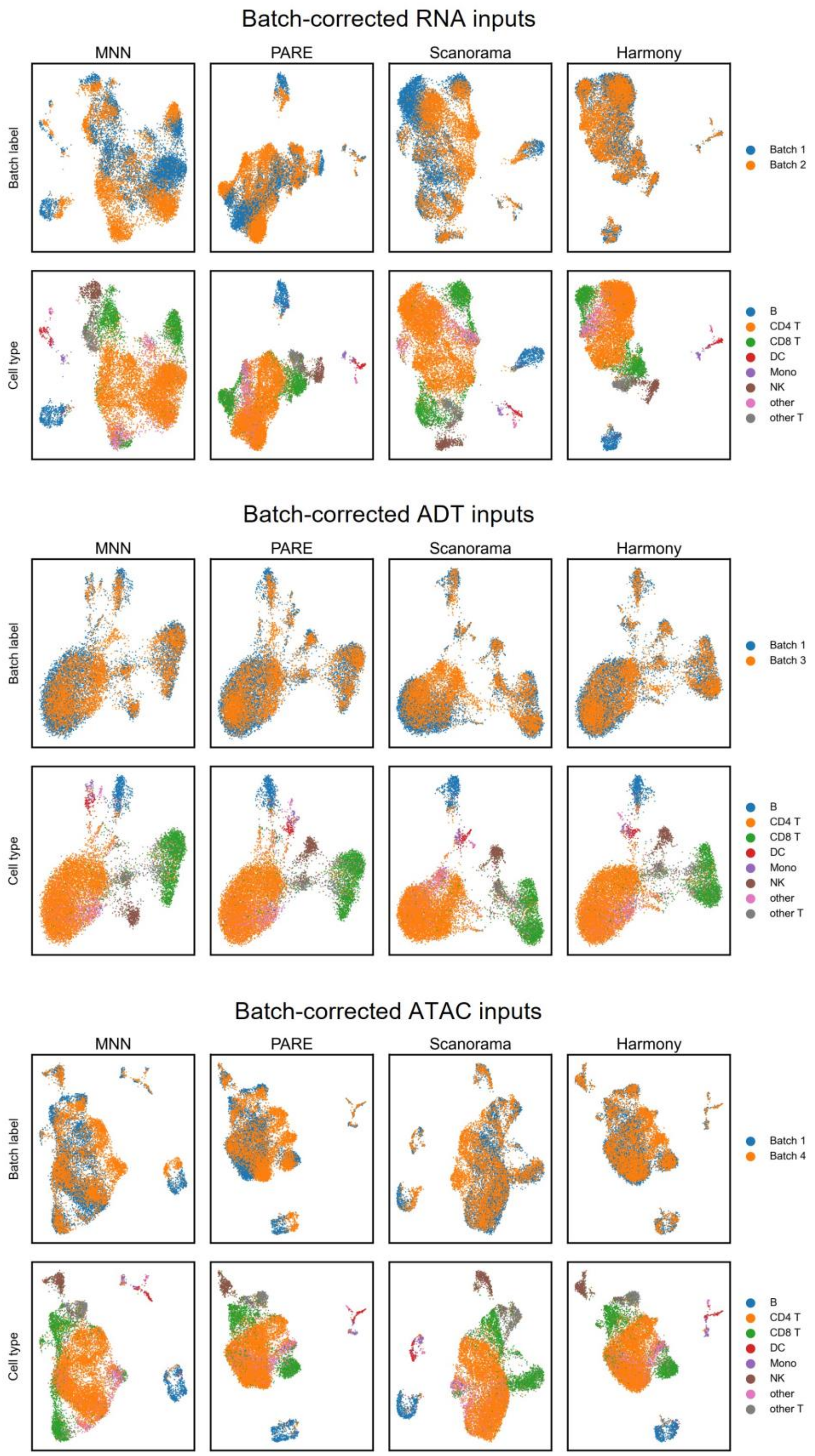


Supplementary Fig. S29. UMAP visualizations of batch corrected modality inputs processed from four batch correction methods.

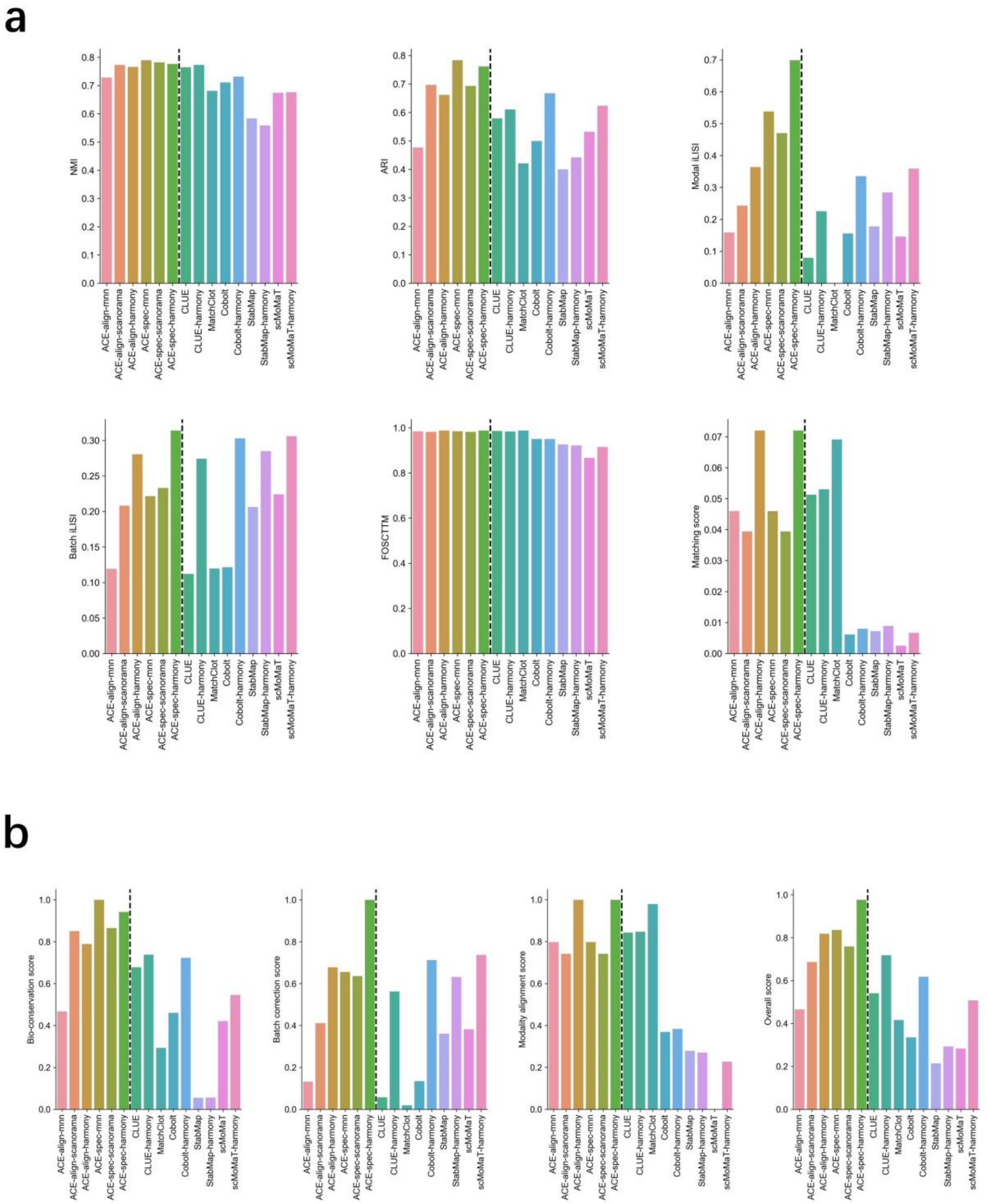


Supplementary Fig. S30. Evaluation of the impact of three batch correction methods on the performance of ACE-align and ACE-spec in CITE dataset. (a) Values of specific evaluation metrics, including NMI, ARI, modal iLISI, batch iLISI, FOSCTTM and matching score. (b) Bio-conservation score, batch-correction score, modality-alignment score and overall score.

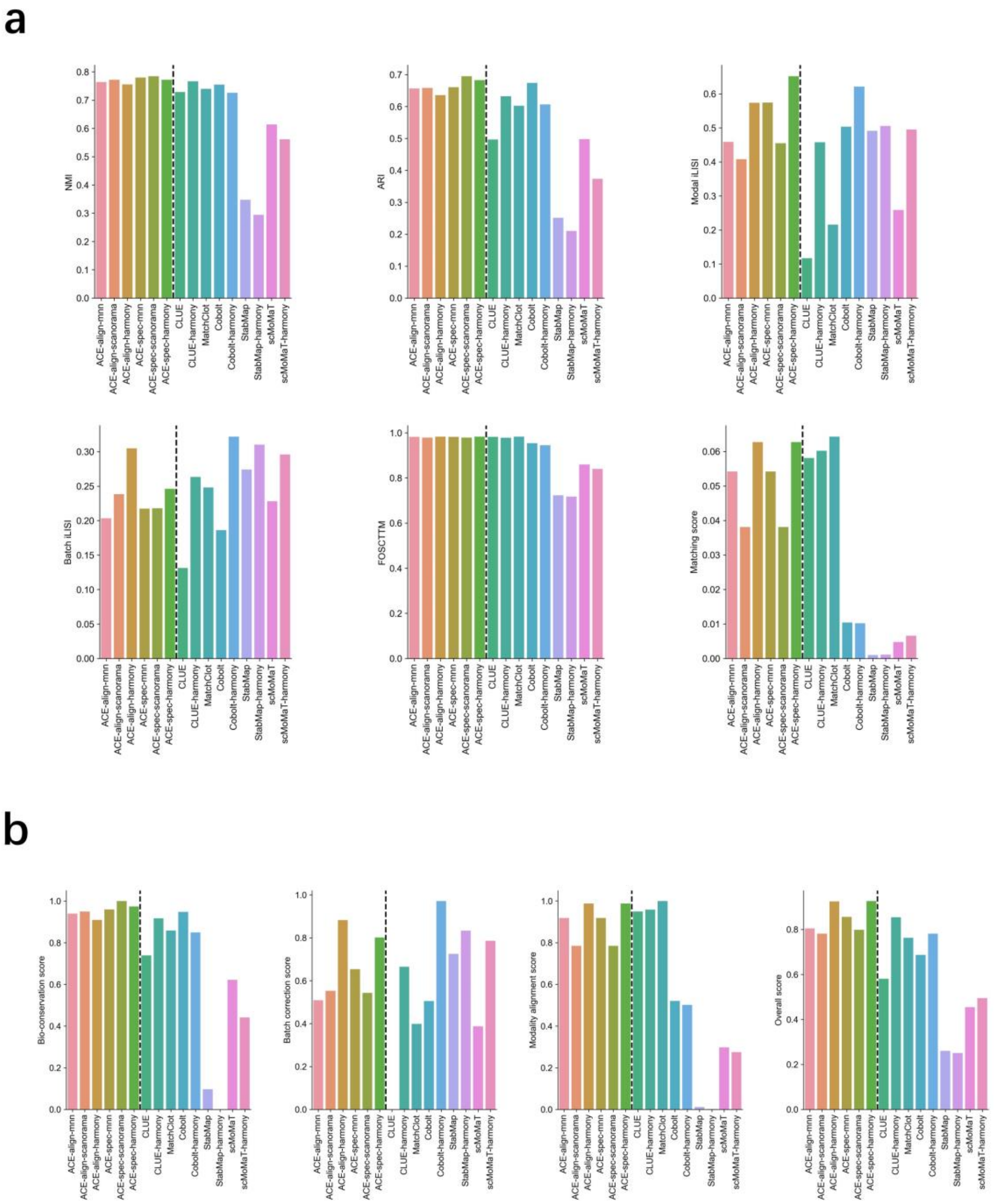


Supplementary Fig. S31. Evaluation of the impact of three batch correction methods on the performance of ACE-align and ACE-spec in Muliome dataset. (a) Values of specific evaluation metrics, including NMI, ARI, modal iLISI, batch iLISI, FOSCTTM and matching score. (b) Bio-conservation score, batch-correction score, modality-alignment score and overall score.

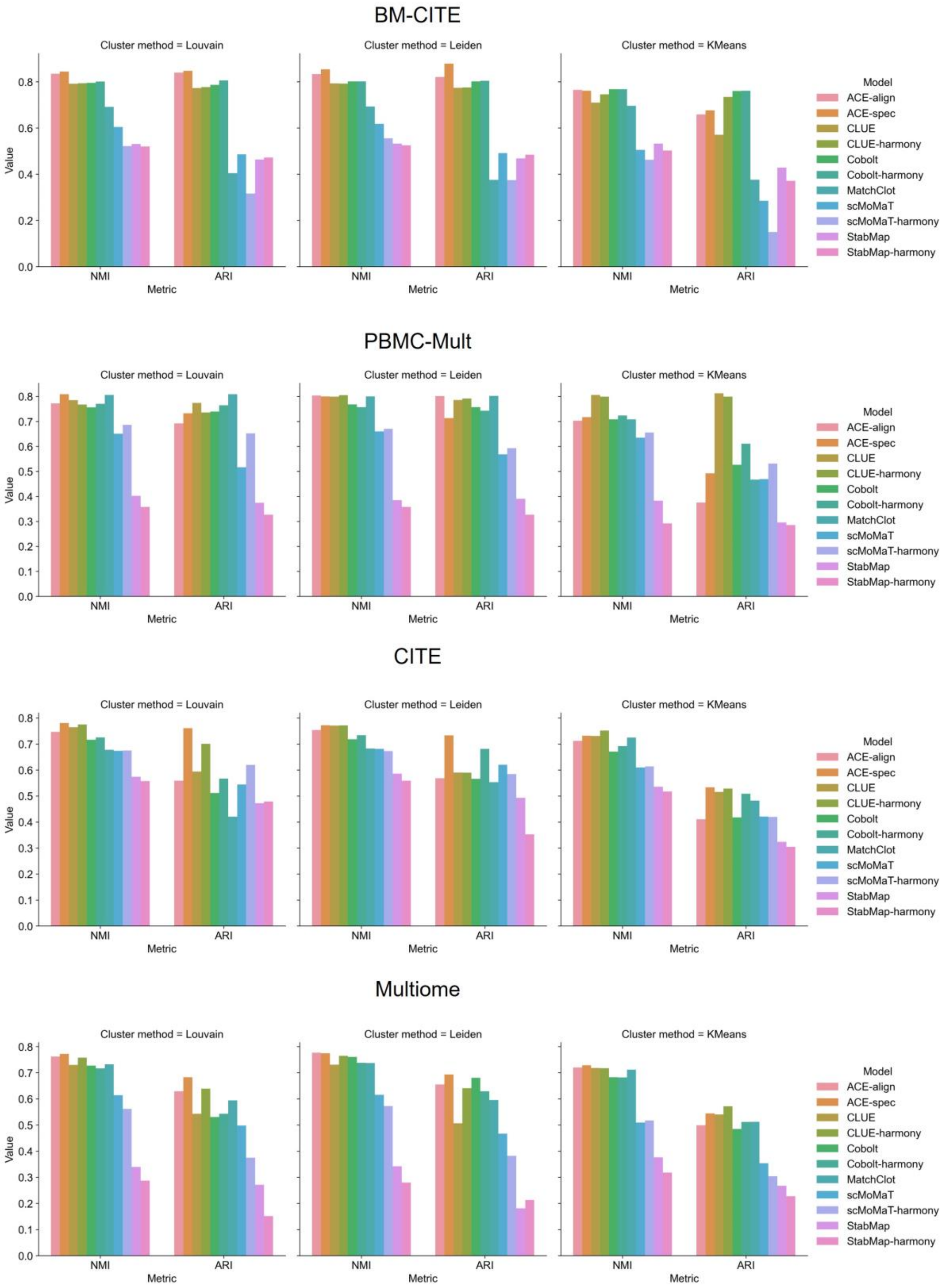


Supplementary Fig. S32. Evaluation of the impact of three clustering algorithms on supervised metrics (NMI and ARI) for various integration methods across four datasets.

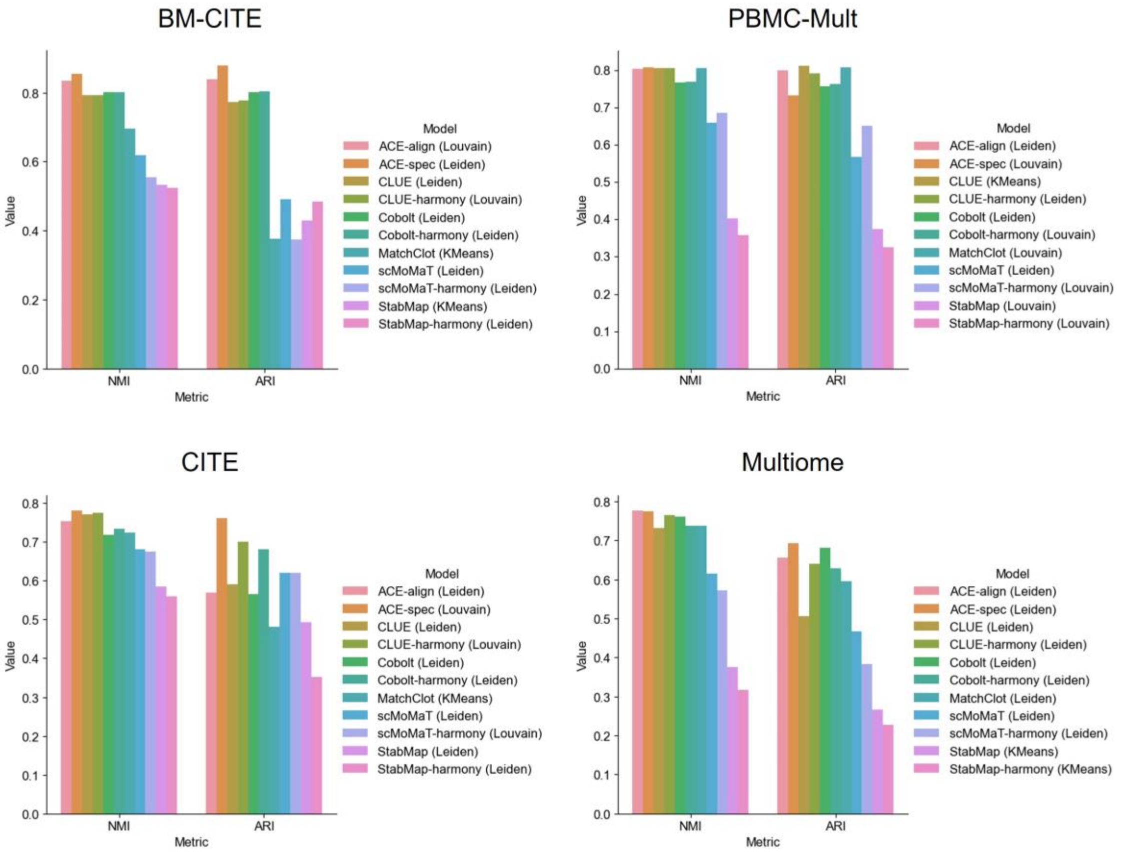


Supplementary Fig. S33. Comparison of various integration methods combined with their optimal clustering algorithm, selected based on the highest NMI scores.

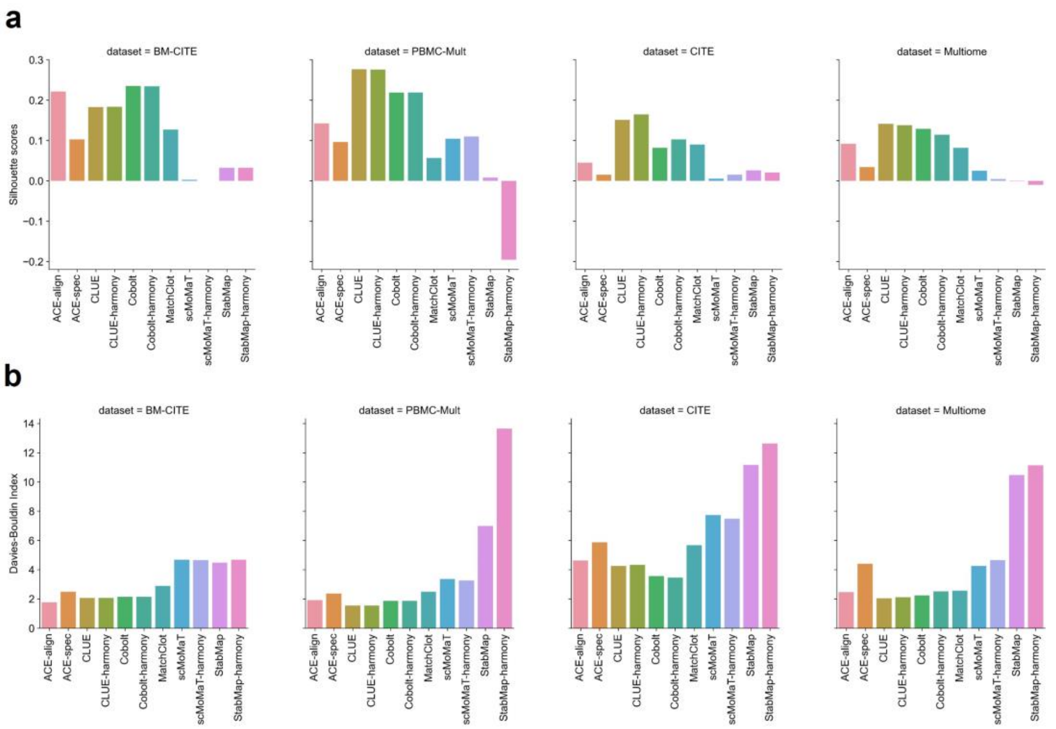


Supplementary Fig. S34. Comparison of various integration methods using unsupervised metrics: silhouette score in (a) and Davies-Bouldin index in (b).

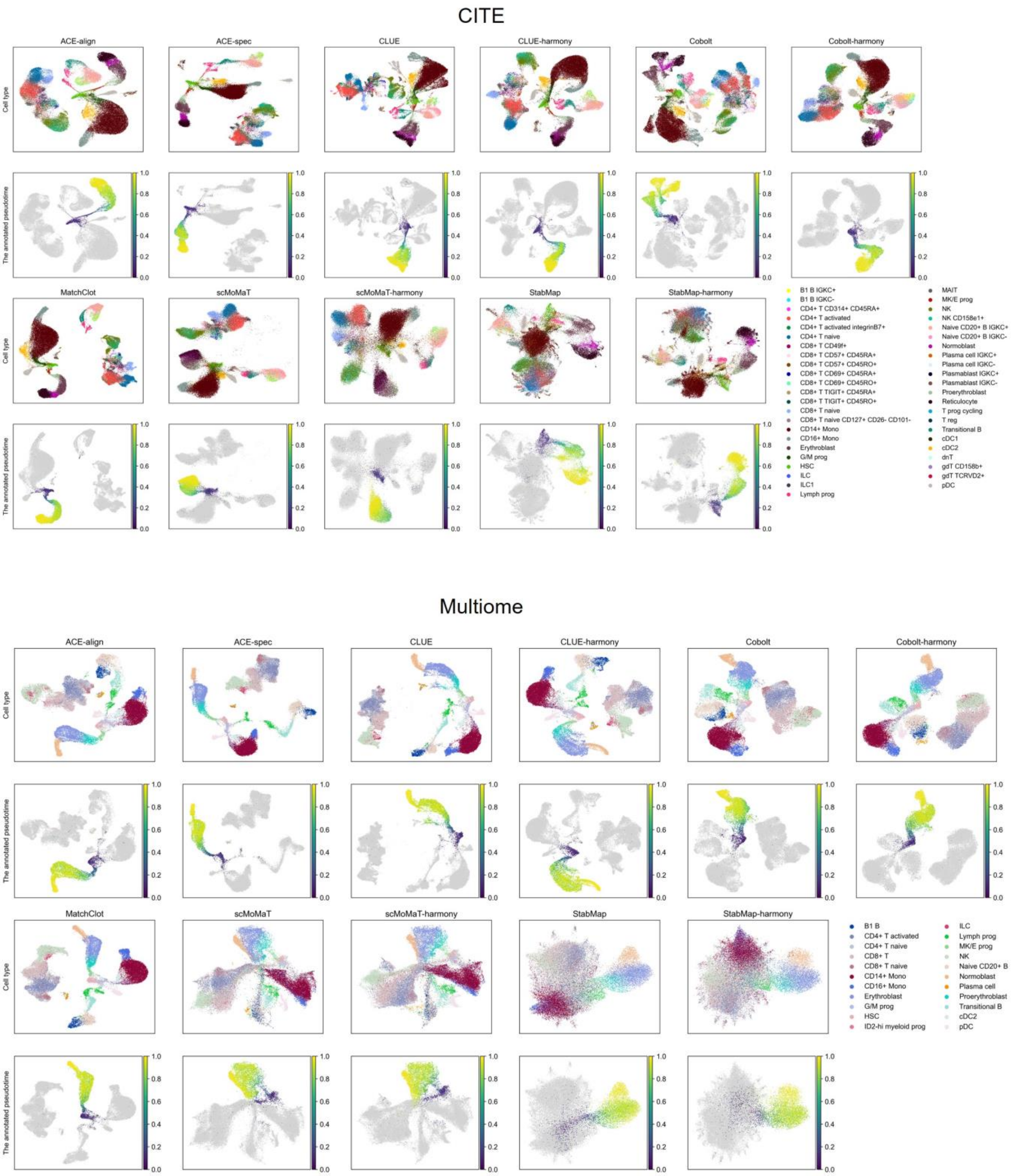


Supplementary Fig. S35. UMAP visualizations of embeddings generated by various integration methods in CITE and Multiome datasets, with cells colored by annotated cell type and pseudotime.

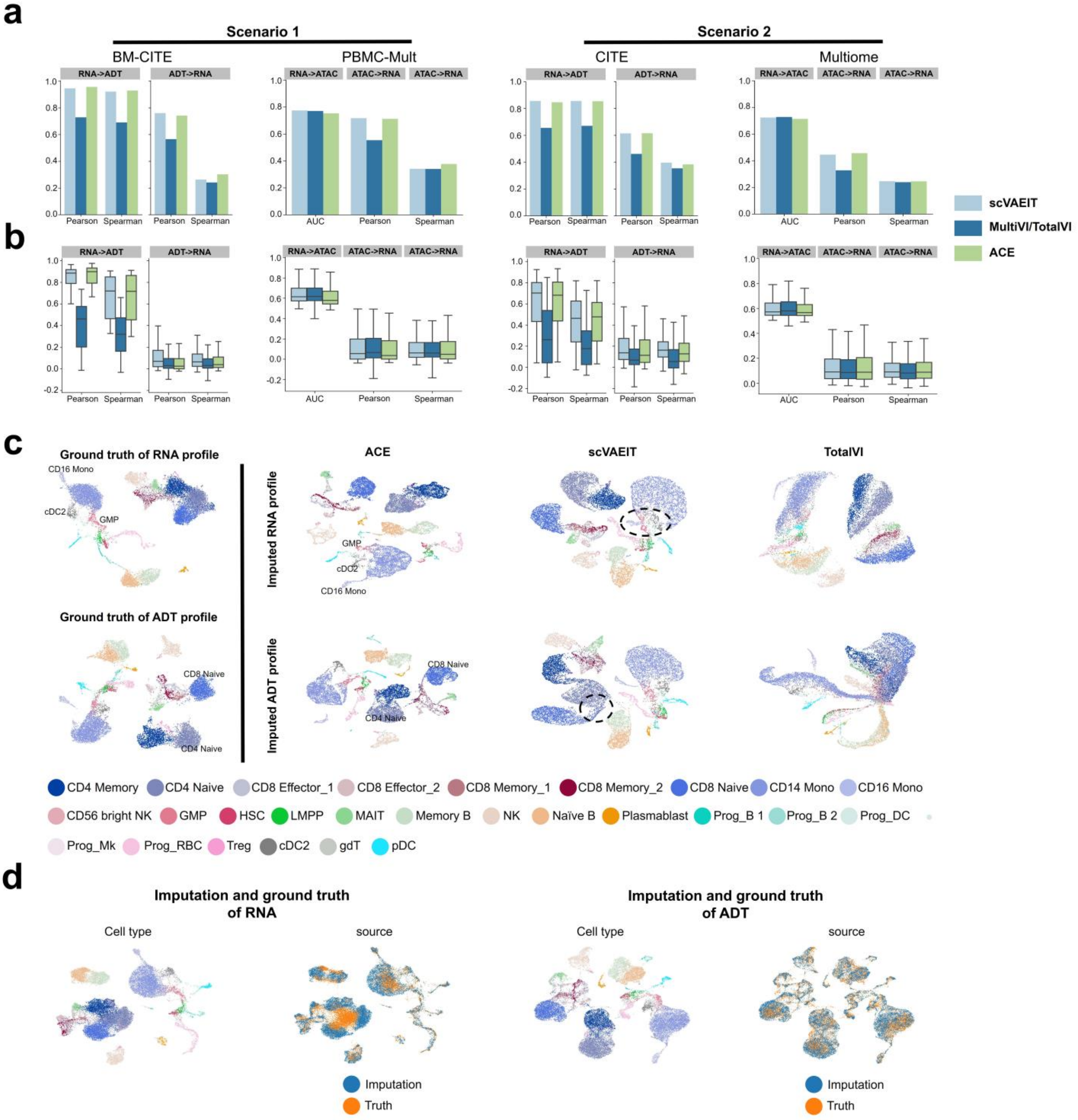


Supplementary Fig. S36. Evaluation of performance in imputing missing modality profiles. (a) Bar plots displaying overall scores of scVAEIT, MultiVI/TotalVI, and ACE in different subtasks of different imputation scenarios. (b) Box plots displaying metric scores of imputation for all individual features. (c) UMAP plots for cells with imputed features (RNA and protein) on the BM-CITE dataset from ACE, scVAEIT, and TotalVI. Cells are colored by cell types. (d) UMAP plots for cells with imputed features and cells with corresponding ground truth of profiles. For each feature, cells are colored by cell types (the same as (c)) in the left and colored by their source in the right.

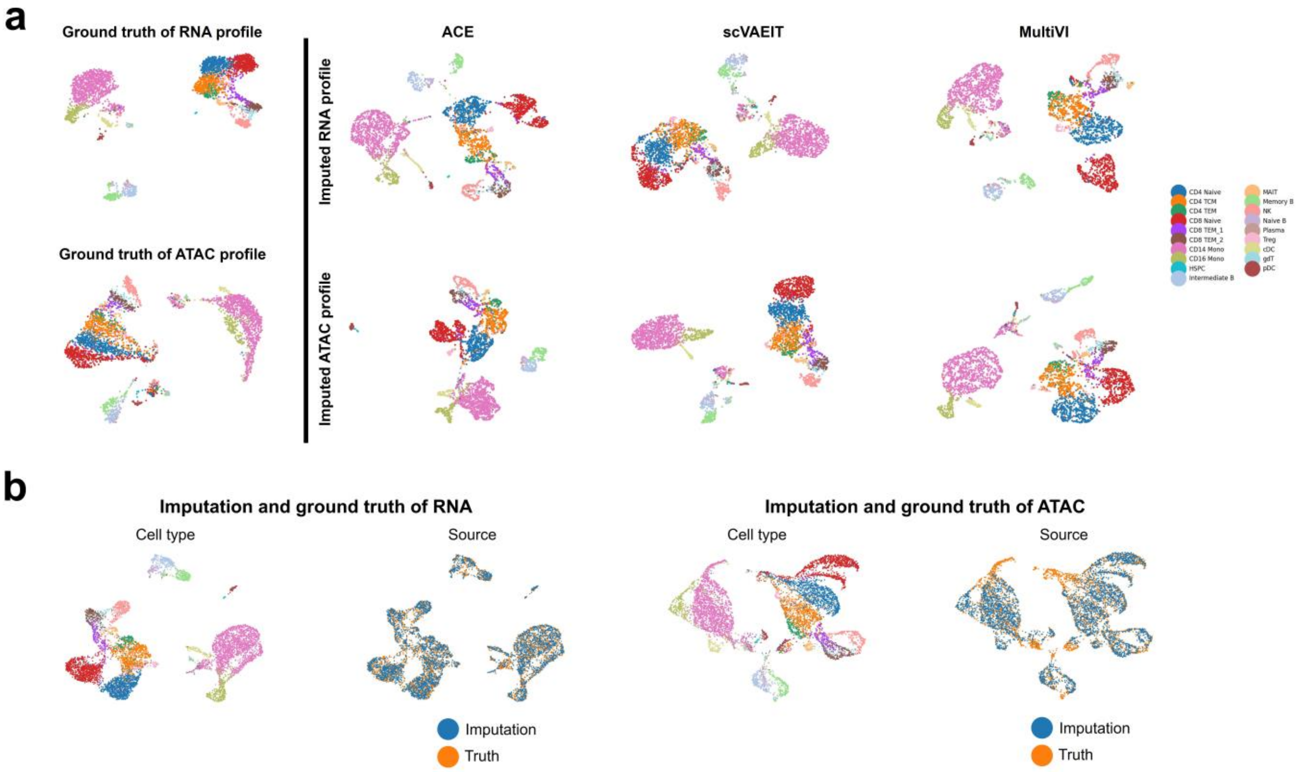


Supplementary Fig. S37. UMAP plots of imputed features on PBMC-Multiome dataset. (a) UMAP plots for cells with imputed features (RNA and ATAC) on this dataset from ACE, scVAEIT, and MultiVI. Cells are colored by cell types. (b) UMAP plots for cells with imputed features and cells with corresponding ground truth of profiles. Within each feature, cells are colored by cell types (the same as (a)) in the left and colored by their source in the right.

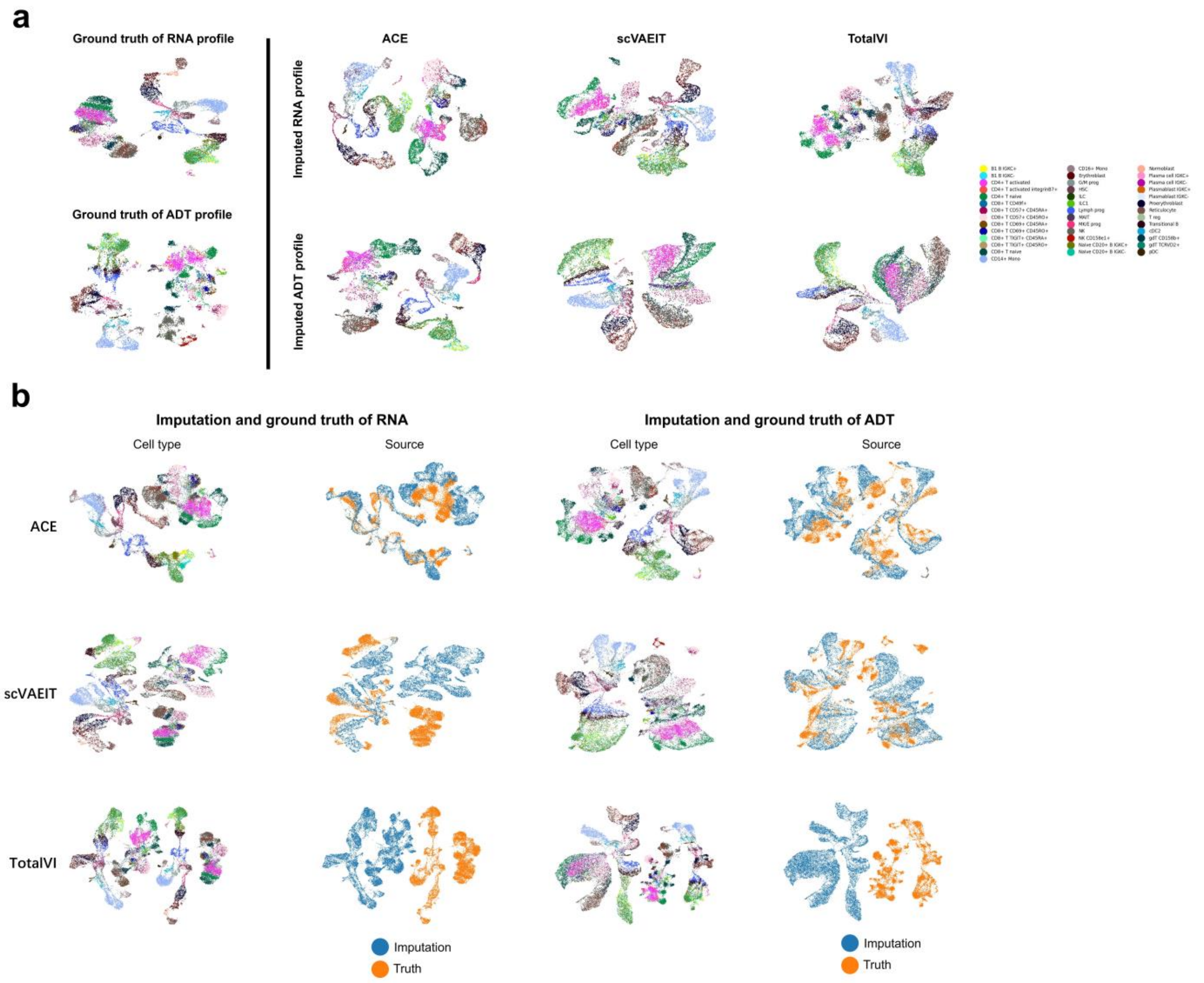


Supplementary Fig. S38. UMAP plots of imputed features on the CITE dataset. (a) UMAP plots for cells with imputed features (RNA and protein) on this dataset from ACE, scVAEIT, and TotalVI. Cells are colored by cell types. (b) UMAP plots for cells with imputed features from different methods and cells with corresponding ground truth of profiles. Within each feature, cells are colored by cell types (the same as (a)) in the left and colored by their source in the right.

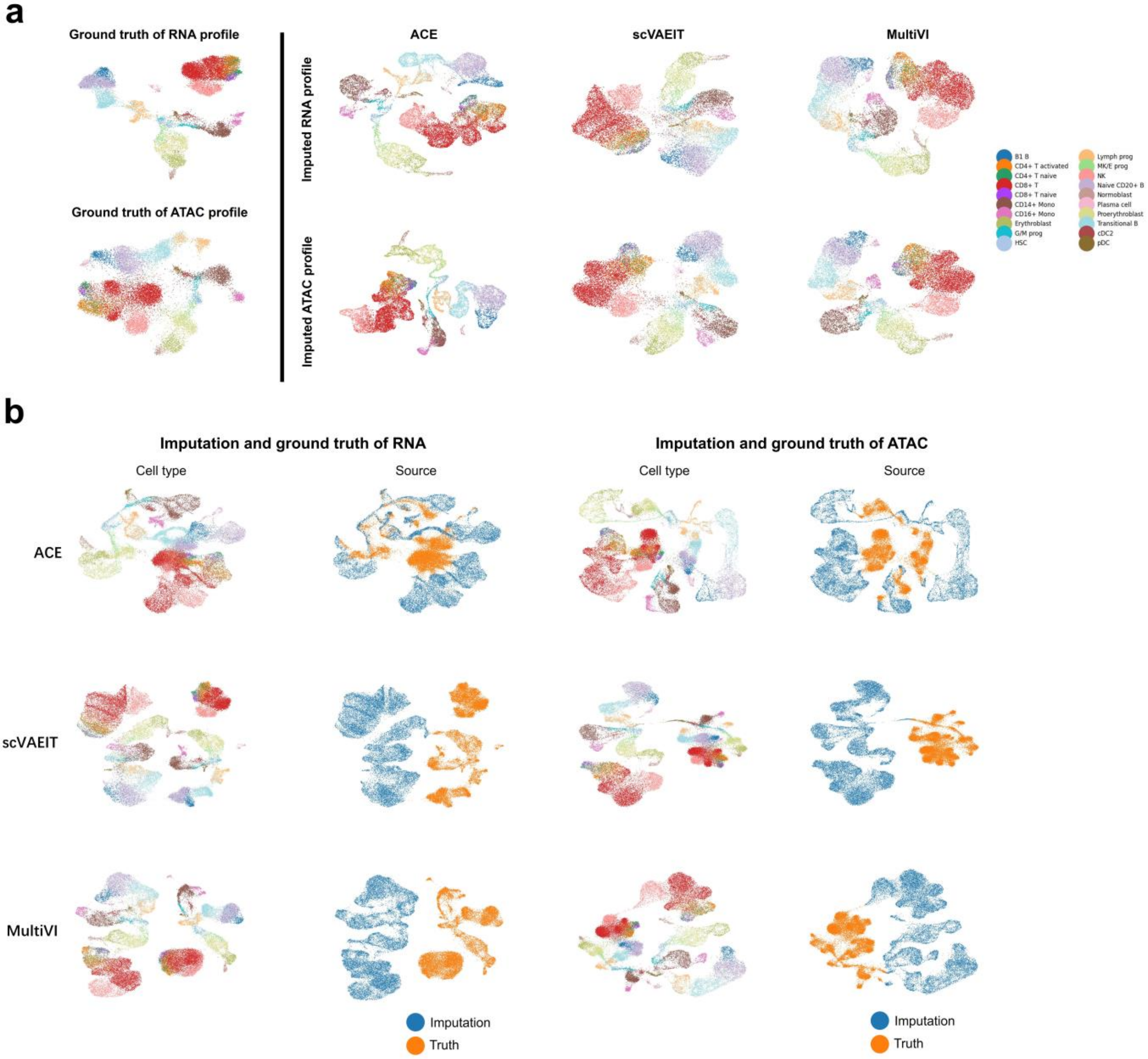


Supplementary Fig. S39. UMAP plots of imputed features on the Multiome dataset. (a) UMAP plots for cells with imputed features (RNA and ATAC) on this dataset from ACE, scVAEIT, and MultiVI. Cells are colored by cell types. (b) UMAP plots for cells with imputed features from different methods and cells with corresponding ground truth of profiles. Within each feature, cells are colored by cell types (the same as (a)) in the left and colored by their source in the right.

#### Supplementary Note A1

Detail of evaluation metrics:

Biological preservation metrics compromise NMI and ARI. NMI and ARI both evaluate the overlap of two clustering. Score 0 corresponds to random clustering and score 1 corresponds to the perfect match for both NMI and ARI. We performed Louvain clustering to obtain the best match between clusters and cell type labels. Louvain clustering was performed at a resolution range of 0.1 to 2 in steps of 0.1, and the clustering output with the highest NMI with the labels was used. We calculated ARI and NMI by the Python functions adjusted_rand_score() and normalized_mutual_info_score() from the scikit-learn library, respectively.

We used graph iLISI as the batch correction metric. Graph iLISI score is a diversity score to assess batch mixing degree, which are computed from neighborhood lists per node from integrated kNN graphs. Following Stabmap, we not only use batch labels ($\text{iLISI}_{\text{batch}}$) to compute iLISI but also used modality labels ($\text{iLISI}_{\text{mod}}$) to compute iLISI. For example, if one batch is measured with RNA and ADT, then its modality label is RNA+ADT, whereas if one batch is measured with RNA, then its modality label is RNA. It can be regarded as evaluation at two levels of resolution, in which modality label is at coarse-grained resolution. Score 0 corresponds to separation of batches and score 1 corresponds to the perfect mixing of batches. We calculated graph iLISI scores by the Python function scib.metrics.lisi.ilisi_graph() from the scib package.

Modality alignment evaluation metrics compromise FOSCTTM (fraction of samples closer than the true match) and matching score (MS). FOSCTTM measures the alignment degree between modalities embeddings. For each modality pair $(m_1,\ m_2)$, it is defined as [14]:

$$\text{FOSCTTM}_0(m_1,m_2) = \frac{1}{2N}\left(\sum_i \frac{N_i^{m_1}}{N} + \sum_i \frac{N_i^{m_2}}{N}\right)$$

$$N_i^{m_1} = |\{l \mid \|z_l^{m_1} - z_i^{m_2}\|_2 < \|z_i^{m_1} - z_i^{m_2}\|_2\}|$$

$$N_i^{m_2} = |\{l \mid \|z_l^{m_2} - z_i^{m_1}\|_2 < \|z_i^{m_2} - z_i^{m_1}\|_2\}|$$

where $N$ is the number of cells that's measured with modalities $m_1$ and $m_2$; $N_i^{m_1}$ denotes the number of cells in modality $m_1$ that are closer to $z_i^{m_2}$ than $z_i^{m_1}$ to $z_i^{m_2}$, and $N_i^{m_2}$ is the same; If existing three modalities in one dataset, we calculated $\text{FOSCTTM}_0$ for every modality pair and took an average. Finally, we rescaled FOSCTTM as 1-$\text{FOSCTTM}_0$, where score of 1 indicates perfect modality alignment. To calculate matching score between two modalities $m_1, m_2$, a cross-modality matching matrix $P$ is first constructed by computing a Jaccard index of cross-modality nearest neighbors in the aligned embedding space [14]:

$$P_{m_1,m_2}(i,j) = \frac{|(NN^{m_2}(x_i^{m_1}) \cap NN^{m_2}(x_j^{m_2})) \cup (NN^{m_1}(x_j^{m_2}) \cap NN^{m_1}(x_i^{m_1}))|}{|(NN^{m_2}(x_i^{m_1}) \cup NN^{m_2}(x_j^{m_2})) \cup (NN^{m_1}(x_j^{m_2}) \cup NN^{m_1}(x_i^{m_1}))|}$$

where $NN^{m_2}(x_i^{m_1})$ denotes the set of cell $x_i^{m_1}$'s nearest neighbors in modality $m_2$; $NN^{m_2}(x_j^{m_2})$ denotes the set of cell $x_j^{m_2}$'s nearest neighbors in modality $m_2$; the other is the same. $|\cdot|$ denotes the number of elements in a set. Following CLUE, the number of nearest neighbors is set to 1. The matching score is computed as follows:

$$\text{MS} = \frac{1}{N}\sum_i \sum_j \tilde{P}_{m_1,m_2}(i,j) \cdot \delta_{i,j}$$

where $\tilde{P}$ denotes row-normalized $P$. $\delta_{i,j}$ is 1 if cell $i$ and $j$ were measured in the same cell and 0 otherwise. $N$ is the number of cells. If existing three or more modalities in one dataset, we calculated matching score for every modality pair and took an average. Matching score has a range of [0, 1], where a score of 1 indicates a perfect alignment.

#### Supplementary Note A2

Detail of compared methods settings:

Cobolt: We run Cobolt following its tutorial (https://github.com/epurdom/cobolt/blob/master/docs/tutorial.ipynb). We first selected highly variable genes or peaks and input the count data matrix into the model for training. Most of training hyperparameters are set to default values: training iteration=100, latent dimension=10, batch size=128. The default learning rate is 0.005 but sometimes it will cause numeric issues. If so, we set the learning rate to 0.001. When executing 'calc_all_latent' function on the CITE-ASAP dataset, we set the parameter 'target' as '[True, True, False]', otherwise it cannot execute normally. Since Cobolt cannot well deal with the batch effect within the same modality, we added harmony as the post-processing step to correct its final embedding across all experiments.

scMoMaT: We run scMoMaT following its tutorial (https://github.com/PeterZZQ/scMoMaT/blob/main/demo_scmomat.ipynb). We first selected highly variable genes or peaks and input the count data matrix into the model for training. Note scMoMaT generates pseudo-count RNA matrix using scATAC-seq data, but we find this step had negative impact in most of cases. Thus, we ignored this step across all experiments. The reason is possibly that the pseudo-count RNA matrix is noisy which will introduce much noise and reduce model performance. We used the default parameter settings in all datasets except the BM-CITE dataset. For the BM-CITE dataset, we found that setting training epochs to 4000 caused overfitting so we set the training epochs to 2000.

StabMap: We run StabMap following its tutorial (https://github.com/MarioniLab/StabMap/tree/main/vignettes). We first selected highly variable genes or peaks and input the log-normalized data matrix into the method for training. All the parameters are set to default values except the 'maxFeatures' parameter which we tuned it according to the total number of features across all modalities. Since StabMap cannot well deal with the batch effect within the same modality, we added harmony as the post-processing step to correct its final embedding across all experiments.

MatchClot: We run MatchClot following its pipeline (https://github.com/AI4SCR/MatchClot/tree/main/tutorials). We first trained MatchClot on all bi-modal datasets using following preprocessing steps: TF-IDF transformation (except the protein data), log-normalization, principal component analysis (PCA, except protein data), batch correction for each modality respectively using harmony. The batch corrected low-dimensional representations or protein expression profiles are input for training. The number of max training epoch is set to 1000 and other training parameters are set to default values across all experiments. Then, we perform model inference using the same inputs as the training step and saved the final cell embeddings. Note that MatchClot proposes to use optimal transport (OT) method to refine the neighborhood graph among cells, which can be seen as an advanced post-processing step for model outputs (and this step can be applied to other methods as well). Considering the fairness of comparison, we did not perform this step for MatchClot.

CLUE: We run CLUE following its pipeline (https://github.com/openproblems-bio/neurips2021_multimodal_topmethods/tree/main/src/match_modality/methods/clue). We used the data count matrix as input and performed CLUE's required preprocessing pipeline for different modality inputs. The preprocessing parameters followed their settings for different sequencing techniques. We first trained CLUE on the batches measured with multiple modalities. CLUE provided two groups of reference training parameters. One group is for CITE dataset and the other one is for Multiome dataset. The hyper parameters for training on all datasets except the Multiome dataset followed the setting on the CITE dataset. We found that even on the PBMC-Mult dataset, the setting on CITE dataset works better than the other one. After training, we saved the pretrained model weights and preprocessing parameters. Next, we performed the second-round training of CLUE on the whole data of each dataset. The preprocessing parameters and training parameters followed the same settings in the first round. Since CLUE cannot well handle the batch effects within the same modality, we added harmony as the post-processing step to correct its final embedding across all experiments.

scVAEIT: We run scVAEIT following its pipeline (https://github.com/jaydu1/scVAEIT/blob/main/example.ipynb). We first selected highly variable genes or peaks and input the log-normalized data matrix into the method for training. Following the steps for processing ATAC features, we split the peak features into multiple chunks by their chromosome id and each chunk will be processed by a separate neural network. Note that scVAEIT can perform transductive imputation and inductive imputation. Specifically, we can train scVAEIT on the bridge batches and the batches with missing modality and perform inference on the training data. Also, we can train scVAEIT on the bridge batches first and perform inference on the batches with missing modality. We experimentally found that inductive setting led to better performance. The hyper parameters of training followed the default settings for different sequencing technologies.

TotalVI: We run TotalVI following its pipeline (https://docs.scvi-tools.org/en/stable/tutorials/notebooks/multimodal/totalVI.html). We first selected highly variable genes or peaks and input the count data matrix into the method for training. Similar to scVAEIT, TotalVi can also perform transductive and inductive imputation. We experimentally found that inductive setting led to better performance on the BM-CITE and CITE datasets. Most of training hyper parameters followed the default settings. When setting up anndata object, we did not use 'batch_key' parameter because we found that it would greatly decrease the method's performance.

MultiVI: We run MultiVI following its pipeline (https://docs.scvi-tools.org/en/stable/tutorials/notebooks/multimodal/MultiVI_tutorial.html). We selected highly variable genes or peaks and input the log-normalized data matrix into the method for training. Similar to TotalVI, we found that inductive imputation obtained better performance on the PBMC-Mult and Multiome datasets. Also, we did not use 'continuous_covariate_keys' parameter because it hurt model performance.

For ACE-align and ACE-spec, we developed two preprocessing pipelines tailored to different datasets:

1. For the raw count assays, we performed highly variable feature selection first. For the ATAC modality, we selected the top 50000 highly variable peaks using Scanpy's highly_variable_genes function [15], and for the RNA modality, we used the same function to select the top 10000 highly variable genes. We then performed normalization: for ATAC peak counts, we applied a TF-IDF transformation followed by log-normalization, while for the RNA and protein modalities, we used log-normalization. Singular value decomposition (SVD) was used to decompose the normalized RNA and ATAC data into low-dimensional representations. We set the dimensionality of output as 192 for the RNA modality and 256 for the ATAC modality. The output dimensionality followed MatchClot's settings [16]. Finally, we performed z-score scaling (zero-mean and standard variance) for each cell's representation within each modality.
2. As in the first pipeline, we performed feature selection for the RNA and ATAC modalities using the same parameters and applied identical normalization steps for both. For the protein modality, we used centered log-ratio normalization. Finally, we used principal component analysis (PCA) to decompose the RNA and ATAC data into low-dimensional representations. The output dimensionality was set as 50.

We used 'sklearn.decomposition.TruncatedSVD' from the scikit-learn package [17], an SVD implementation optimized for efficient handling of sparse matrices. We used the 'arpack' SVD solver because we empirically found that it was faster. For the PCA algorithm, we used the function provided by Scanpy, scanpy.pp.pca. We used the first pipeline for CITE, Multiome and DOGMA datasets. The second pipeline was used for BM-CITE, PBMC-Mult, CITE-ASAP and COVID-19 datasets.

#### Supplementary Note B

To confirm the modality gap phenomenon (i.e., intra-modality embeddings of different cells can be still closer than inter-modality embeddings of the same cell after training with the InfoNCE objective), we analyzed the distance between the intra-modality embeddings and inter-modality embeddings in the BM-CITE dataset where we observed modality gap phenomenon for the InfoNCE loss. Specifically, we trained ACE-align with the InfoNCE and our proposed loss function, respectively and used the modality-specific embeddings of the multi-modal batch (simultaneously measured with RNA and ADT) for analysis. For each cell's RNA embedding, we calculated its distance to the paired ADT embedding (denoted as RNA-ADT), and we also picked up its nearest neighbor from the RNA embeddings (only within the multi-modal batch) and calculated the distance (denoted as RNA-RNA). We did the same thing for the cell's ADT embedding (denoted as ADT-

RNA and ADT-ADT distance).

For the embeddings derived from InfoNCE and ours, we plotted the scatter of each cell's RNA-RNA distance and RNA-ADT distance (also for ADT-RNA and ADT-ADT distance), as shown in Supplementary Fig. S5. When the Manhattan distance metric was used and the temperature T was set to 0.01, we can observed in the InfoNCE' s results that the RNA-ADT distance was much larger than RNA-RNA distance and ADT-RNA distance was much larger than ADT-ADT distance (Supplementary Fig. S5a), which supported our statement that intra-modality embeddings of different cells can be still closer than inter-modality embeddings of the same cell after training with the InfoNCE loss. In contrast, our proposed loss function resulted in similar embedding distance. As T increased, the distance between inter-modality embeddings derived from the InfoNCE loss became closer to that of intra-modality embeddings. This aligns with the visualization in Supplementary Fig. S25, which shows the modality gap shrinking as T increases. For our proposed loss, increasing T didn't result in significant differences. When we changed the distance metric to the Euclidean distance, we still observed similar results (Supplementary Fig. S5b).

Moreover, we increased the neighborhood size to see if the observations remained consistent over a larger area. Specifically, for each cell's RNA embedding, we picked up its top 10 nearest neighbors from ADT embeddings and calculated the distance (RNA-ADT distance). Also, we picked up the top 10 nearest neighbors from RNA embeddings and calculated the distance (RNA-RNA distance). We did the same thing for ADT embeddings (ADT-RNA and ADT-ADT distance). When using the Manhattan distance metric and T as 0.01, we can observe in InfoNCE' s results that the distance between intra-modality embeddings was smaller than that of inter-modality embeddings in various neighborhood scales (Supplementary Fig. S6a). As for our results, the distance difference was moderate. Even if changing the distance metric and varying the temperature T, the observations remained consistent (Supplementary Fig. S6b).

Overall, the experimental evidence confirmed that intra-modality embeddings of different cells can be still closer than inter-modality embeddings of the same cell after training with the InfoNCE objective, while our proposed loss function can effectively reduce this modality gap.

#### Supplementary Note C

To investigate the impact of the UMAP parameters on the visualizations, we first obtained the embeddings of ACE-align, ACE-spec, scMoMaT and Cobolt in the BM-CITE and PBMC-Mult datasets. Then, we visualized these embeddings by varying hyper-parameters of UMAP, including number of neighbors and minimum distance, following the same setting in the UMAP hyperparameter experiment [18].

The visualization results were shown in Supplementary Figs. S18-S21. For both datasets, we could observe that varying the hyper-parameters did not affect the observed batch mixing quality. Specifically, ACE-align and ACE-spec well mixed three batches across all parameter settings whereas scMoMaT only mixed batches 1 and 3 for both datasets (Supplementary Figs. S18 and S20). For the PBMC-Mult dataset, Cobolt mixed batches 1 and 3 while a portion of batch 2 was isolated (Supplementary Fig. S20). This pattern was consistent across all parameter settings. However, the observed heterogeneity (cell types) was slightly affected by the UMAP parameters. Specifically, within each method's visualizations, increasing the number of neighbors would make each cluster of cell type look smaller (Supplementary Figs. S19 and S21). Consequently, those separated cell types look more isolated due to the increased inter-cluster distance. The partially overlapping cell types became increasingly difficult to distinguish due to the decreased scale of separation boundaries (similar to zooming out). Conversely, increasing the minimum distance would make each cluster of cell type look larger. The distance between each pair of cell types decreased, while the partially overlapping cell types became more distinguishable due to the increasing scale of separation boundaries (similar to zooming in). Therefore, adjusting the hyper-parameters of UMAP is similar to viewing cellular heterogeneity from different scales. However, the embedding outputs from integration methods basically determined the cellular heterogeneity that we can observe. For example, in the BM-CITE and PBMC-Mult datasets, Naïve B cells and Memory B cells are two slightly overlapping clusters in ACE-spec's visualization (min_dist=0.0125, n_neighbors=5, Supplementary Figs. S19 and S21). We could still distinguish between these two cell types after adjusting the hyper-parameters. However, these two cell types were mixed in scMoMaT's visualization and varying the parameters did not enable separation between them (Supplementary Figs. S19 and S21). In the PBMC-Mult dataset, CD4 TCM and CD4 TEM cells were separable in ACE-spec's visualizations while they were mixed in scMoMaT's visualizations (Supplementary Fig. S21). In addition, we observed that the small number of neighbors resulted in many isolated small clusters, which is a common phenomenon for ACE-align/spec, scMoMaT and Cobolt (Supplementary Figs. S19 and S21). This is unsurprising because small numbers of neighbors make model focus on small neighborhoods and the visualization become more dispersed.

Furthermore, we explored the hyper-parameters of UMAP for visualizing ACE-spec's embeddings on the CITE-ASAP and COVID-19 datasets. We observed that the visualizations with different hyper-parameters consistently aligned with the original cell-type annotations and the clustering labels (Supplementary Figs. S22 and S23). For example, clusters 11, 14, 15 and 16 in the CITE-ASAP dataset were rare clusters but were clearly separable across the parameter settings (Supplementary Fig. S22b). Clusters 14, 16, 17 and 20 were rare clusters in the COVID-19-RNA dataset but were also separable across the parameter settings (Supplementary Fig. S23b). These results again demonstrated the robustness of the observed cellular heterogeneity.

We concluded that the embedding outputs from integration methods primarily determined the cellular heterogeneity that we can observe. UMAP visualization is an approach to represent the heterogeneity in a 2-dimensional space. Adjusting the hyper-parameters of UMAP is analogous to zooming in or zooming out on the data and our hyper-parameter analysis illustrated the stability and robustness of the observed cellular heterogeneity.

#### Supplementary Note D

Ablation study of our proposed contrastive learning objective

To investigate the effectiveness of our proposed learning objective for ACE-align, we compared it with InfoNCE loss on BM-CITE and CITE datasets. First, by keeping other parameters the same, we evaluated the two loss functions for different settings of temperature values. Supplementary Fig. S24a shows that on BM-CITE dataset, our proposed loss function clearly outperforms InfoNCE for different temperature values with respect to bio-conservation and batch-correction metrics. Especially when the temperature is small, the improvement of our loss function is more significant. As for the modality alignment metrics, FOSCTTM and MS, two loss functions have similar performance. This is attributable to the fact that these two metrics solely quantify inter-modality relationships, neglecting intra-modality relationships. Consequently, our proposed loss function is not expected to exhibit enhanced performance. On the CITE dataset, the improvement of our loss function is also clear with respect to batch-correction metrics and bio-conservation metrics (Supplementary Fig. S24b). UMAP plots on two datasets also demonstrate that InfoNCE results in modality gap phenomena when T is smaller than 0.02, whereas our proposed loss function consistently mixes all batches and modalities (Supplementary Fig. S25). For the reason why the bridge batch mixed well with RNA-modal batch, this is because we only use RNA modality of bridge batch to infer its final embeddings.

However, we also noticed that the improvement of our loss function on CITE dataset is lower than on the BM-CITE dataset. We hypothesized that this is because the protein modality in CITE dataset contains more features (n=134) than in BM-CITE dataset (n=25). More protein features indicate that the information content between two modalities is more balanced. Thus, it's easier to avoid modality gap phenomenon on CITE dataset. To investigate how two loss functions perform with different number of protein features, we randomly selected 10, 20, 40, 80 protein features from CITE dataset and evaluated two loss functions in different cases. Each selection was repeated three times to avoid randomness. We set the T=0.1 for both losses to maximize the performance of InfoNCE. Supplementary Fig. S24c shows that for different numbers of protein features, our proposed loss function still outperforms InfoNCE with a clear margin of batch correction metrics. This indicates that our proposed loss function can better handle the cases with greatly imbalanced information content between modalities, thereby enhancing its generalization capabilities.

#### Supplementary Note E

Removing batch effects prior to model training has a notable impact on the model's performance [19]. We selected Harmony [20] as the batch correction method because it has been validated as a robust solution in numerous studies. To assess the impact of different batch correction methods on the model's performance, we compared three commonly used batch correction methods, Mutual Nearest Neighbors (MNN) [21], Scanorama [22], Harmony [20], and a new method, PARE [23] (as suggested by the reviewer), on the CITE, Multiome and DOGMA datasets, all of which contain batch effects within individual modality. We found that PARE required substantial computational resources, resulting in memory overflow on the CITE and Multiome datasets. To incorporate PARE for comparison, we subsampled cells from the CITE and Multiome datasets with sampling ratios of 5% and 10% (we named these datasets CITE-5%, CITE-10%, Multiome-5% and Multiome-10%). On the full CITE and Multiome datasets, we compared MNN, Scanorama and Harmony. On the DOGMA, CITE-5%, CITE-10%, Multiome-5% and Multiome-10% datasets, we compared all four batch correction methods. Results were shown in Supplementary Figs. S26-S31.

For the four subsampled datasets, ACE-spec combined with each of the four batch correction methods yielded similar NMI scores (ACE-align showing a similar trend), while the impact on ARI scores was more notable (Supplementary Fig. S26). For ACE-align, Scanorama led to more robust and higher ARI scores, followed by MNN, while PARE and Harmony had similar ARI scores. For ACE-spec, Scanorama and Harmony led to higher ARI scores. PARE led to the lowest ARI scores. In terms of batch correction metrics (modal iLISI and batch iLISI), Harmony led to the highest scores for both ACE-align and ACE-spec across the four subsampled datasets, except for the Multiome-10% dataset, where ACE-spec with PARE attained the highest modal iLISI score (Supplementary Fig. S26). ACE-spec with PARE consistently ranked in the top two for both modal iLISI and batch iLISI scores across all four datasets. For ACE-align, MNN, Scanorama and PARE had similar iLISI scores. In terms of modality alignment metrics (FOSCTTM and matching score), four batch correction methods' scores were close (Supplementary Fig. S26). To provide a more intuitive understanding of the effects of the four batch correction methods, we plotted their resulting UMAPs for the CITE-5% and Multiome-5% datasets, as shown in Supplementary Fig. S27. Harmony's output displayed a more uniform mixing of batches within each modality, with well separated cell types, which was consistent with the comparison results. The other three methods had similar visualization results, where batches were mixed and the cellular heterogeneity was preserved.

For the DOGMA dataset, Harmony attained the highest NMI and ARI for ACE-align and ACE-spec among the four batch correction methods, and outperformed other integration methods (Supplementary Fig. S28a). PARE achieved the highest modal iLISI and batch iLISI scores for ACE-spec and followed by Harmony, which were tied with the highest scores of other integration methods. ACE-align achieved the highest iLISI scores with Harmony. The four batch correction methods led to similar FOSCTTM scores for ACE-align and ACE-spec, and all surpassed other integration methods (Supplementary Fig. S28a). However, Harmony attained the highest matching score for ACE-align and ACE-spec, and significantly outperformed other integration methods. We visualized the batch corrected outputs from four batch correction methods using UMAP. Harmony mixed batches better within each modality, which helped to guide the modality alignment across batches (Supplementary Fig. S29). In terms of the overall scores, combining the four batch correction methods with ACE-spec all outperformed other state-of-the-art integration methods (Supplementary Fig. S28b). For ACE-align, Harmony, MNN and PARE also achieved competitive overall performance. For the CITE dataset, ACE-spec with MNN, Scanorama and Harmony achieved the highest overall scores among all integration methods (Supplementary Fig. S30). For the Multiome dataset, Harmony helped ACE-align and ACE-spec achieve the highest overall scores while MNN and Scanorama also achieved competitive performance (Supplementary Fig. S31).

Together, we conducted an exploration for the impact of four batch correction methods on our proposed framework. We concluded that using different batch correction methods did have an impact on the performance, and the performance differences were mainly reflected in the batch correction and modality alignment. Generally, Harmony had the best batch correction performance and consequently resulted in better modality alignment across batches. PARE achieved close or even higher batch correction scores as Harmony but still showed batch separation in the UMAP visualizations. Additionally, one limitation of PARE was that it took much longer to finish the integration than the other three methods. For instance, Harmony finished integration for DOGMA dataset within 2 minutes whereas PARE took 48 hours to finish on our server. However, from the perspective of overall scores, all four batch correction methods helped ACE-align and ACE-spec achieve top performance among all integration methods, which demonstrated the robustness and generalizability of our proposed framework.

#### Supplementary Note F

To assess whether different clustering algorithms may introduce biases in performance evaluation, we tested two additional clustering algorithms: Leiden and KMeans. Leiden is an improved shared nearest neighbor clustering (SNN)-based algorithm that addresses certain limitations of the Louvain algorithm, such as disconnected communities and instability in the results. KMeans, a widely used clustering method, is known for its computational efficiency and effectiveness with spherical, well-separated clusters, but it is less suited for data with irregular shapes or overlapping clusters. On the BM-CITE, PBMC-Mult, CITE, and Multiome datasets, we applied three clustering algorithms to the embeddings generated by each integration method and assessed the clustering performance using NMI, ARI. We also calculated two unsupervised metrics for each method's embeddings, silhouette coefficient and Davies-Bouldin index (DBI), which considers both within-cluster and between-cluster distance. Following the scIB benchmarking [24], the unsupervised metrics were calculated using cell type labels. Silhouette coefficient has a value range from -1 to 1, where -1 indicates strong misclassification, and +1 indicates dense and well-separated clusters. DBI has a value from 0 to infinity, and values closer to zero indicate a better cluster separation.

Using the Louvain and Leiden clustering algorithms, ACE-spec achieved the highest NMI and ARI scores on the BM-CITE, CITE, and Multiome datasets (Supplementary Fig. S32). Using Leiden algorithm, ACE-align ranked in the top three for NMI and ARI scores on BM-CITE, PBMC-Mult and Multiome datasets. Cobolt and CLUE using the Louvain and Leiden algorithms showed NMI scores comparable to ACE-spec while StabMap showed the lowest NMI and ARI scores among all methods. When using the KMeans clustering algorithm, the NMI and ARI scores of ACE-align and ACE-spec both dropped compared to using Louvain and Leiden (Supplementary Fig. S32). This is unsurprising because the UMAP visualizations suggested that many cell types in ACE-align's and ACE-spec's embedding space exhibited non-convex shapes (Supplementary Figs. S4 and S8), which are challenging for the KMeans algorithm to accurately capture. Nevertheless, ACE-spec still achieved competitive NMI and ARI scores among all methods while ACE-align's ARI scores were relatively lower. For other integration methods, we also observed performance drop when changing the clustering algorithm as KMeans, especially for ARI scores (Supplementary Fig. S32). However, CLUE and CLUE-harmony achieved higher NMI and ARI scores with KMeans on PBMC-Mult dataset. To compare the best clustering performance of each integration method, we selected the clustering algorithm that achieved the highest NMI score on each dataset and compared all methods' performance again. The results showed that ACE-spec achieved the highest NMI and ARI scores on the BM-CITE, CITE, and Multiome datasets, while ACE-align ranked among the top three for NMI and ARI on the BM-CITE, PBMC-Mult, and Multiome datasets (Supplementary Fig. S33).

In terms of silhouette scores, ACE-align ranked second on the BM-CITE dataset, behind Cobolt (Supplementary Fig. S34a). On the other datasets, CLUE achieved the highest silhouette scores, followed by Cobolt. Across all four datasets, ACE-spec's silhouette scores were lower than those of the top-performing methods. Regarding the DBI (lower score indicates better embedding quality), ACE-align achieved the lowest DBI on the BM-CITE dataset, followed by Cobolt and CLUE (Supplementary Fig. S34b). On the other datasets, ACE-align, Cobolt, and CLUE consistently ranked in the top three. However, ACE-spec exhibited higher DBI scores compared to the leading methods. The poor unsupervised metric scores of ACE-spec, particularly for the CITE and Multiome datasets, can be attributed to the distortion of many clusters in ACE-spec's embedding space. Based on the UMAP visualizations, these clusters appeared "stretched and squeezed" (Supplementary Fig. S8), which increased within-cluster distance and decreased the between-cluster distance. However, this deformation may reflect genuine developmental trajectories. For example, in both the CITE and Multiome datasets, there existed an erythrocyte development trajectory involving Hematopoietic stem cell (HSC), Megakaryocyte and Erythrocyte (MK/E) progenitor, Proerythroblast, Erythroblast, Normoblast and Reticulocyte [25]. In ACE-spec's embeddings, these cell types formed a continuous and stretched structure, consistent with the pseudotime that was annotated in the original study [25] (Supplementary Fig. S35). In contrast, Cobolt and other methods presented this trajectory less clearly. Therefore, although the stretched clusters resulted in lower unsupervised metric scores for ACE-spec, this pattern accurately captured the underlying biological trajectories.

Overall, ACE-align and ACE-spec achieved superior NMI and ARI scores with SNN-based clustering algorithms but showed decreased scores with the KMeans algorithm. ACE-align outperformed ACE-spec in terms of unsupervised metric scores, but both were relatively poorer than the best-performing methods.

### Supplementary Note G

Evaluation of ACE's performance in imputing missing molecular layers

The goals of mosaic integration are not only focused on projecting different batches into a consensus space, but also help impute raw features for those unmeasured modalities in batches as a natural biproduct of integration procedure [26, 27]. We realized that our cross-modal matching strategy can be applied to impute the missing molecular layers, as described in the Method section. To evaluate its performance, we organized two scenarios. In both scenarios, there exist bridge batches and multiple batches measured with the same one modality. In scenario 1, batch effects do not exist between bridge batches and single-modal batches. Accordingly, BM-CITE dataset and PBMC-Multiome dataset are used for scenario 1, and the methods are required to reconstruct the missing profiles from the available profiles, which includes four subtasks: predicting RNA from protein (ADT —> RNA), predicting protein from RNA (RNA —> ADT), and predicting RNA from ATAC (ATAC—>RNA), and predicting ATAC from RNA (RNA—>ATAC). In scenario 2, batch effects exist between bridge batches and single-modal batches. CITE dataset and Multiome dataset are used in this case, where the same four prediction subtasks are included. Following scVAEIT, we used Spearman correlation coefficient (SCC) and Pearson correlation coefficient (PCC) as evaluation metrics of RNA and protein imputation, and used area under the ROC curve (AUROC) as the evaluation metric of ATAC imputation. The overall score of each metric is computed by concatenating all cell's features into one vector. TotalVI, MultiVI, and scVAEIT were evaluated for comparisons. TotalVI and MultiVI are restricted to bimodal analysis, in which TotalVI jointly models RNA and proteins and MultiVI jointly models RNA and ATAC. Detailed method settings can be found in Supplementary Note A2.

In scenario 1, ACE and scVAEIT show similar overall scores of different metrics across four subtasks, and they outperform TotalVI/MultiVI with a distinct margin in subtasks of RNA—>ADT, ADT—>RNA, and ATAC—>RNA (Supplementary Fig. S36a). Evaluation on each feature separately also shows that ACE and scVAEIT achieve similar metric scores and they are more robust than ToTalVI/MultiVI (Supplementary Fig. S36b). Next, we visualized cells with their imputed features using UMAP. Supplementary Fig. S36c shows that ACE's imputed RNA and protein profiles on BM-CITE dataset well preserve the original cellular heterogeneity. When visualizing the imputed features and the ground truth of missing modalities together, it's observed that they overlap well (Supplementary Fig. S36d). Compared to the imputed features of scVAEIT, we found that ACE's imputations show better separation of cell types (Supplementary Fig. S36c). For instance, within ACE's imputed RNA profiles, CD16 Mono cells are distinct from GMP and cDC2 cells, whereas in scVAEIT's imputations, CD16 Mono cells overlap with these two cell types. Within the ground truth of missing RNA profiles, these three types are also separated. CD4 Naïve and CD8 Naïve are two separated clusters within ACE's imputed protein profiles while they are overlapped within scVAEIT's imputations. The ground truth of protein profiles also shows that these two types are separated. UMAP plots on PBMC-Multiome dataset also demonstrate ACE's capability in reconstructing raw omics features (Supplementary Fig. S37). Nevertheless, these visualizations also reveal that ACE's imputation is not smoothing enough compared to the ground truth, which is an inherent limitation of ACE due to discrete imputation process.

In scenario 2, batch effects exist between bridge batches and single-modal batches. Although we can correct the batch effects within the shared modality, it's highly possible that the imputations of missing modality still have batch effects with the ground truth. Evaluation results are shown in Supplementary Figs. S36a-b. ACE shows similar performance to scVAEIT and they generally outperform TotalVI/MultiVI. However, compared to scenario 1, the metric scores dropped clearly. For example, ACE's PCC of protein imputation on CITE dataset dropped by 11% compared to the BM-CITE dataset (PCC dropped by 17% for the RNA imputation). On the same dataset, scVAEIT's PCC of protein imputation dropped by 9% and PCC of RNA imputation dropped by 19%, indicating that batch effects have a notable impact on the quality of imputation. We visualized cells with the imputed features and cells with the ground truth of missing profiles together and observed that those cells with imputed features are separated to those cells using real features (Supplementary Figs. S38-S39). The obvious separation can also be observed in scVAEIT's results. We think this is a common problem for existing reconstruction methods and leave this problem for future work.
